# Uveal Melanoma Oncogene *CYSLTR2* Encodes a Constitutively Active GPCR Highly Biased Toward Gq Signaling

**DOI:** 10.1101/663153

**Authors:** Emilie Ceraudo, Mizuho Horioka, Jordan M. Mattheisen, Tyler D. Hitchman, Amanda R. Moore, Manija A. Kazmi, Ping Chi, Yu Chen, Thomas P. Sakmar, Thomas Huber

**Affiliations:** Laboratory of Chemical Biology and Signal Transduction, The Rockefeller University, New York, NY, USA; Human Oncology and Pathogenesis Program, Memorial Sloan Kettering Cancer Center, New York, NY, USA; Louis V. Gerstner Jr. Graduate School of Biomedical Sciences, Memorial Sloan Kettering Cancer Center, New York, NY, USA; Weill Cornell Graduate School of Medical Sciences, Cornell University, New York, NY, USA; Department of Discovery Oncology, Genentech Inc, South San Francisco, CA, USA; Department of Medicine, Memorial Sloan Kettering Cancer Center, New York, NY, USA; Department of Medicine, Weill Cornell Medical College, 1300 York Avenue, New York, NY, USA

**Author notes:** E.C. and M.H. contributed equally to this work. **Data and materials availability:** All data needed to evaluate the conclusions in the paper are presented in the paper or the Supplementary Materials. **Author Contributions:** E.C., M.H., J.M.M., T.D.H., A.R.M., P.C., Y.C., T.P.S., and T.H. designed the research; E.C., M.H., J.M.M., T.D.H., A.R.M., M.A.K. performed the research; E.C., M.H., J.M.M., T.D.H., A.R.M., P.C., Y.C., T.P.S., and T.H. analyzed and interpreted the data; E.C., M.H., J.M.M., T.P.S., and T.H. wrote the article.

## Abstract

The G protein-coupled receptor (GPCR) cysteinyl-leukotriene receptor 2 (CysLTR2) with a single amino acid mutation at position 3.43 (Leu replaced with Gln at position 129 in transmembrane helix 3) causes uveal melanoma in humans. The ability of CysLTR2-L129Q to cause malignant transformation has been hypothesized to result from constitutive activity. We show that CysLTR2-L129Q is a constitutively active mutant (CAM) that strongly drives Gq/11 signaling pathways in melan-a melanocytes and in HEK293T cells in culture. However, the mutant receptor only very weakly recruits beta-arrestins 1 and 2. The mutant receptor displays profound signaling bias while avoiding arrestin-mediated downregulation. The mechanism of the signaling bias results from the creation of a hydrogen-bond network that stabilizes the active G protein signaling state through novel interactions with the highly-conserved NPxxY motif on helix 7. Furthermore, the mutation destabilizes a putative allosteric sodium-binding site that usually stabilizes the inactive state of GPCRs. Thus, the mutation has a dual role of promoting the active state while destabilizing inactivating allosteric networks. The high degree of constitutive activity renders existing orthosteric antagonist ligands of CysLTR2 ineffective as inverse agonists of the mutant. CysLTR2 is the first example of a GPCR oncogene that encodes a GPCR with constitutive highly biased signaling that can escape cellular downregulation mechanisms.

## Introduction

The superfamily of G protein-coupled receptors (GPCR) is the largest gene family encoding cell signaling transmembrane proteins ^1^. About one-quarter of ∼400 non-olfactory GPCRs are therapeutic drug targets ^2^. Large-scale genomic analysis has revealed that one in five individuals carry a missense variant (MV) in a clinically-relevant GPCR gene. The rate of *de novo* germline MVs in a GPCR gene is one in every 300 newborns, and one in seven MVs is observed at functionally relevant sites ^3^. In addition, GPCR genes are commonly mutated in cancer and somatic mutations are found in 20% of tumor samples ^4,5,6^. The functional impact of most observed MVs is not known, leading to the terminology “variants of uncertain (or unknown) significance” (VUS) ^7, 8^.

We recently reported the discovery of a recurrent “hotspot” missense mutation of the gene *CYSLTR2*. The mutant *CYSLTR2* encodes CysLTR2-L129Q that carries a single amino acid substitution at a highly conserved residue in helix 3 (position 3×43) and serves as a driver oncogene in uveal melanoma (UVM) patients ^6^. More recently, the same mutation has also been identified as an oncogenic driver mutation in several other tumors ^9,,10,11^. A variety of mutations in other GPCRs also drive several benign endocrine neoplasms, and a virally-encoded GPCR drives Kaposi’s sarcoma ^12,,13,,14,15^. GPCRs are among the most commonly mutated genes in cancer, but the lack of specific “hotspot” variants makes it difficult to identify and validate individual receptors as driver oncogenes ^4, 5^.

Here we show the precise signaling mechanism of the oncoprotein CysLTR2-L129Q. We show that CysLTR2-L129Q is a constitutively active mutant (CAM) receptor that strongly couples to Gq/11 cellular signaling pathways. However, the receptor CAM only very weakly recruits β-arrestins and thereby avoids cellular down-regulation mechanisms. We propose a model of the molecular activation mechanism of CysLTR2-L129Q in which interaction of L(3×43)Q with the NPxxY motif differentially stabilizes an active state conformation. We also analyzed the genetic variants of CysLTR2 found in cancer and normal controls to identify other potentially constitutively active variants that could have tumor-promoting activity. We screened the impact of the mutations *in silico* by homology modeling of the mutant proteins using active and inactive state templates. In summary, the CysLTR2-L129Q CAM exhibits functional selectivity and escapes β-arrestin-dependent downregulation. Our working hypothesis is that the oncogenic potential of a *CYSLTR2* MVs is related to a gain-of-function in basal signaling through Gq/11 pathways.

## Results

### CysLTR2-L129Q signals through Gq/11-PLC-β pathways

The hallmark of CAM receptors is agonist-independent signaling. To determine the functional phenotype of CysLTR2-L129Q, we determined agonist dose-dependent signaling as a function of receptor gene dosage and compared CysLTR2-L129Q with CysLTR2 wild-type (CysLTR2 wt). CysLTR2 predominantly couples to Gq/11 when treated with the agonist leukotriene D4 (LTD4) (Suppl. Fig. S1). ^16^. We obtained a time-course of basal and LTD-4 dependent IP1 accumulation in HEK293T cells transiently transfected with plasmids for CysLTR2 wt, CysLTR2-L129Q and mock controls. In samples with LTD4-stimulated CysLTR2 wt, the IP1 accumulation increased over the first 100 minutes before reaching a plateau (Suppl. Fig. 2A), whereas the unstimulated CysLTR2 wt samples were indistinguishable from and the mock-transfected controls, both with and without LTD4 treatment. In comparison, the samples transfected with the same amount of DNA encoding for the CysLTR2-L129Q mutant show ligand-independent IP1 accumulation that parallels that seen in the LTD4-stimulated CysLTR2 wt samples, which slowed down the increase after 100 minutes, but continued to levels above the plateau seen in the wild-type receptor sample (Suppl. Fig. 2B).

**Fig. 1.**
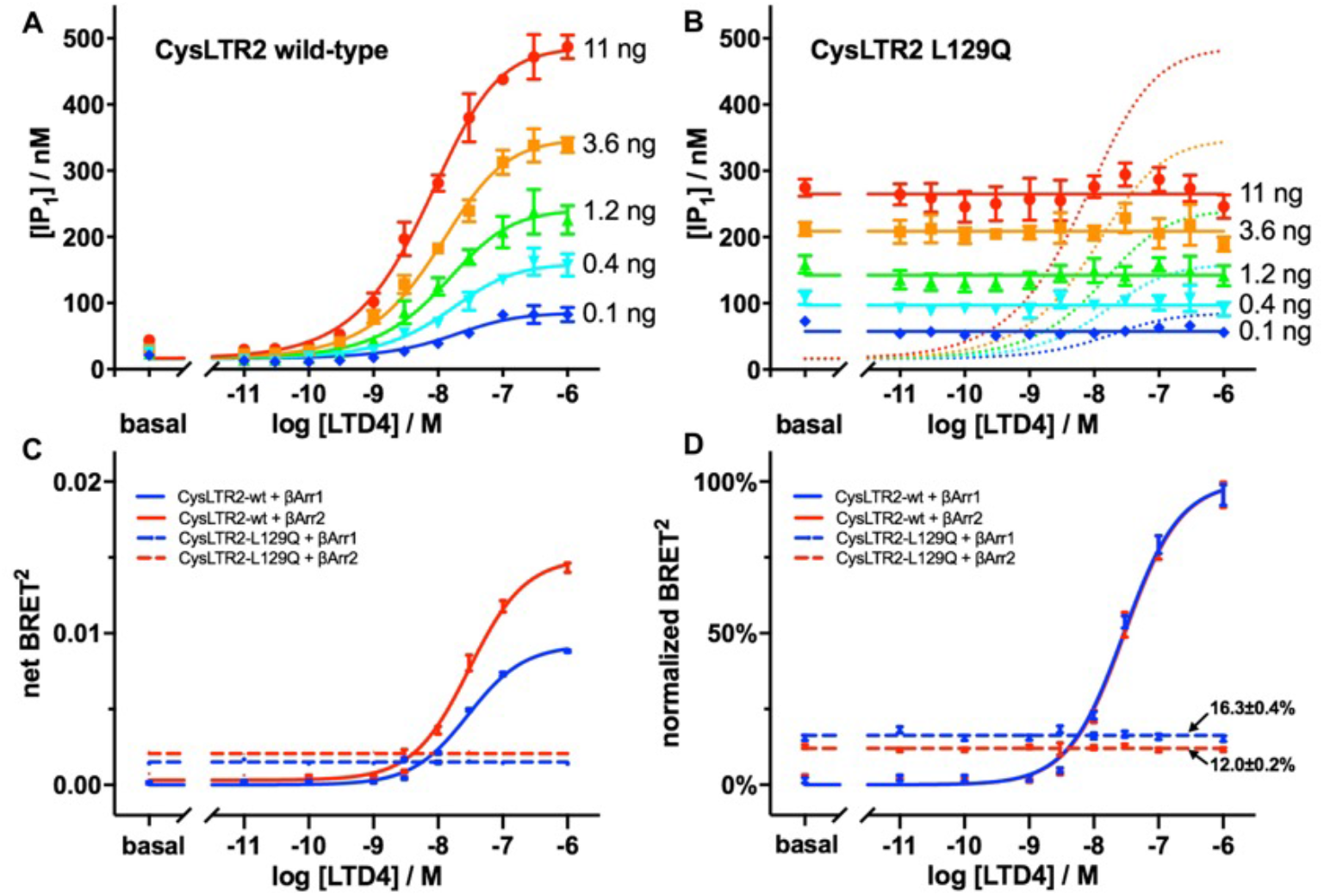
Oncoprotein CysLTR2-L129Q^3.43^ is a Gq-biased CAM that weakly recruits β-arrestins. (**A**) The agonist LTD4 leads to dose-dependent inositol monophosphate (IP1) accumulation in HEK293T cells expressing different levels of CysLTR2 wild-type as controlled by different amounts of receptor-encoding plasmid DNA transfected at constant total DNA (red, 11 ng; orange, 3.6 ng; green, 1.2 ng; cyan, 0.4 ng; blue, 0.1 ng). The curves are fits of the dose-response data to an operational model. (**B**) The corresponding experiment with the mutant CysLTR2-L129Q shows no significant dose-dependent response to LTD4, whereas the ligand-independent basal IP1 accumulation increases with increasing amount of CysLTR2-L129Q-encoding plasmid DNA. The data were fit to a horizontal line. The results show that CysLTR2-L129Q is a CAM with a basal activity corresponding to about 70% of the WT receptor maximally stimulated with agonist. Data are expressed as mean ± SEM of the concentration of IP1 accumulated (nM) and result from one experiment performed in four technical replicates. Wt and L129Q assays were performed in individual plates with mock-transfected cells as controls. (C,D) β-Arrestin-recruitment BRET^2^ assay with CysLTR2-GFP10 and β-arrestin-Rluc3. (**C**) The LTD4 dose-dependent increase of net BRET^2^ demonstrates recruitment of β-Arrestin 1 (red solid line and points) and β-Arrestin 2 (blue solid line and points) to wild-type CysLTR2. In comparison, the data for CysLTR2-L129Q indicate ligand-independent recruitment of β-Arrestin 1 (blue dashed line and open points) and β-Arrestin 2 (red dashed line and open points) to CysLTR2-L129Q. (**D**) The BRET^2^ data were independently normalized for either β-Arrestin 1 or 2 using the asymptotic endpoints of the sigmoidal fits for the wild-type receptor. The normalized data show nearly perfect overlap of the fitted curves for both β-arrestins binding the wild-type receptor. The ligand-independent β-arrestin recruitment for CysLTR2-L129Q is 16.3±0.4% and 12.0±0.2% for β-Arrestin 1 and 2, respectively. The dose-response data are the mean ± SEM from three independent experiments, with nine concentrations and three technical replicates, each.

LTD4 induces a dose-dependent increase in IP1 accumulation for CysLTR2 wt (Figure 1A), whereas the CysLTR2-L129Q mutant shows little or no response to treatment with LTD4 (Figure 1B). At the same time, CysLTR2-L129Q exhibits very large ligand-independent, basal signaling corresponding to about 70% of the IP1 accumulation obtained from maximally LTD4-stimulated CysLTR2 wt when compared at the same gene dosage. To further characterize the basal and ligand-dependent signaling of the different samples, we analyzed the agonist dose-response data using a sigmoidal fitting function (Suppl. Scheme S1). We compared the sigmoidal fit for each data set with a fit to a horizontal line function as an alternative hypothesis (Suppl. Table S1). The dose-response data for CysLTR2-L129Q in Fig. 1B are best described by a horizontal line, that is, they lack statistically significant response to the agonist LTD4. However, we have observed in other experiments with this mutant a small but statistically significant response to the agonist LTD4 (cf. Fig. 2 and Suppl. Fig. S5). Overall, the results suggest that CysLTR2-L129Q displays a loss-of-function (LoF) phenotype for agonist-dependent signaling as compared with the CysLTR2 wild-type receptor.

The potency of LTD4 at CysLTR2 wt increases with increasing receptor gene dosage (Suppl. Table S1), which corresponds to a left-shift of the sigmoidal agonist dose-response curves (cf. Fig. 1A). This increase in potency as a function of receptor can be described by the Black-Leff operational model ^17^ (Suppl. Scheme S1). We performed a global analysis of the CysLTR2 wt gene dosage-dependent, LTD4 dose-response experiment using individual “log τ” parameters for each gene dosage and shared fitting parameters for all other parameters (Suppl. Table S2). The curves shown in Fig. 1A result from this global analysis. The value of the τ parameter from this analysis suggests that at the highest gene dosage of 11ng DNA per well (1.57pg DNA/cell), the total receptor concentration is approximately 1.7-fold higher than the concentration necessary to reach half-maximal signaling. Therefore, the IP1 accumulation obtained by the fully LTD4-stimulated receptor reaches about 64% of the maximum. However, the Black-Leff operational model is not suitable for the description of agonist dose-response data for receptors with high constitutive activity, such as CysLTR2-L129Q. Even for CysLTR2 wt, the low concentration endpoints of the curves from the global analysis with the Black-Leff model are below the experimental unstimulated data points indicated as the basal tick in Fig. 1A. The sigmoidal fits for the CysLTR2 wt samples from the same data set (Suppl. Table S1) show a small increase of the *bottom* parameter, corresponding to the low concentration asymptote of the sigmoidal curve.

Next, we applied the Slack-Hall operational model ^18, 19^ to quantify the increase in basal signaling that cannot be described by the Black-Leff operational model, which is commonly used to quantify *biased agonism* or *ligand bias* ^20^. The key result is that the Slack-Hall model can be used to quantify the agonist-independent, inherent pathway bias of the receptor referred to as *receptor bias* ^19^. The Slack-Hall model splits the τ parameter of the Black-Leff model into a product of two parameters, χ and ε. The basal response is determined by χ and it is defined as the ratio of [R]*_t_*, the total receptor concentration, and *K_e_*, the receptor concentration producing half-maximal effect in the *absence* of an agonist. In contrast, the τ parameter in the Black-Leff model is the ratio of [R]*_t_* and a different *K_e_*, which is defined as the receptor concentration producing half-maximal effect in the *presence* of a saturating agonist concentration. The ε parameter measures the *intrinsic efficacy* of the ligand. Suppl. Table S4 shows the parameters for the Slack-Hall operational model fitted to the experiments shown in Fig. 1A, B. The difference of log χ calculated from the CysLTR2 wt and CysLTR2-L129Q data and averaged over all receptor gene dosages is −1.22±0.03. Therefore, the constitutive activity of the L129Q mutation is 17-fold higher than that of the wild-type receptor, assuming that the receptor densities are the same. The difference of the log epsilon parameters is 1.51±0.19, which suggests that the intrinsic efficacy of the agonist LTD4 is 33-fold higher for the wild-type receptor as compared to the L129Q mutant receptor.

We tested the functionality of the GFP10-fusion constructs that we developed for the β-arrestin-recruitment assays in an LTD4 dose-response IP1 accumulation assay with different gene dosages (Suppl. Fig. S3A,B, Suppl. Tables S1–S3). The difference of the log χ parameters from the Slack-Hall operational model results for the CysLTR2-GFP10 wt samples and the CysLTR2-GFP10-L129Q samples averaged over all gene doses was −1.68±0.10, which corresponds to 48-fold higher constitutive activity of the L129Q mutant *versus* wild-type. The difference of the log ε parameters was 1.65±0.28, suggesting that the intrinsic efficacy of LTD4 is 45-fold higher for the wild-type receptor as compared to the mutant.

### Characterizing the β-Arrestin-recruitment to CysLTR2-L129Q

Signals from active GPCRs normally get terminated by β-arrestin-dependent mechanisms including desensitization, sequestration, and down-regulation ^21^. We next asked the question of how CysLTR2-L129Q is capable of sustained strong signaling at a level comparable to the fully agonist-stimulated wild-type receptor. CysLTR2 has been shown to bind β-arrestin2 in response to several agonists ^22^. However, little is known about the β-arrestin-dependent desensitization and downregulation of CysLTR2 and CysLTR2-L129Q. We designed a bioluminescence resonance energy transfer (BRET) experiment to quantify the basal and agonist-dependent binding of β-arrestins to CysLTR2 variants.

We generated fusion constructs of CysLTR2 wt and CysLTR2-L129Q with a version of green fluorescent protein (GFP10) that can be used in BRET^2^ assays ^23^ in combination with β-arrestins fused to an engineered variant of *Renilla* luciferase, β-arrestin1-RLuc3 and β-arrestin2-RLuc3 ^24^. Next, we characterized the agonist-dependent β-arrestin recruitment to the wild-type receptor. We performed an initial optimization of the gene dosage and cell density for the BRET2 assay with HEK293T cells transiently expressing wild-type CysLTR2-GFP10 and β-arrestin-RLuc3 to maximize the LTD4-dependent increase in the BRET^2^ ratio. We then performed a time-course experiment to characterize the agonist-dependent β-arrestin-recruitment. The results show that the BRET^2^ ratio increases for approximately ten minutes after addition of the agonist LTD4, before starting to decrease again slowly (Suppl. Fig. S3C,D). The slope of the initial increase increases with higher concentrations of the agonist. The shape of the time-course was similar comparing samples expressing β-arrestin1-RLuc3 and β-arrestin2-RLuc3, but the peak increase seen for β-arrestin2-RLuc3 was almost twice that of β-arrestin1-RLuc3. Such a biphasic BRET β-arrestin-recruitment time-course is typical for GPCRs with “class A” β-arrestin-recruitment phenotype that have transient, weak interactions with β-arrestin and these receptors rapidly recycle after internalization ^25^.

The LTD4 dose-dependent increase of the BRET^2^ ratio for samples transfected with CysLTR2-GFP10 wt and β-arrestin-RLuc3 substantiate the finding from the time-course assay that the agonist-dependent increase of BRET^2^ is larger for β-arrestin2-RLuc3 as compared with β-arrestin1-RLuc3 (Fig. 1C). Even though the agonist-dependent increase was different, the midpoints of the sigmoidal fits of the agonist dose-dependent data for both β-arrestins were identical (Fig. 1D).

### CysLTR2-L129Q poorly recruits β-arrestins

To characterize the effect of the L129Q mutation on β-arrestin-recruitment, we included a set of samples expressing CysLTR2-L129Q-GFP10 in the BRET^2^ experiments. The results from the LTD4 dose-response experiment show a ligand-independent net BRET^2^ ratio of 0.00151±0.0004 for β-arrestin1 and 0.00208±0.0003 for β-arrestin1 (Fig. 1C). To compare these values to the agonist-dependent net BRET^2^ ratio for the wild-type receptor at saturating concentrations, we normalized the data of the L129Q mutant receptor using the *top* and *bottom* parameters for the sigmoidal fits of the wild-type data (Fig. 1D). Interestingly the normalized data reverse the order and show that β-arrestin1 with 16.3±0.4% of the agonist-dependent recruitment by the wild-type is slightly preferred over β-arrestin2 with only 12.0±0.2%. Next, we quantify the constitutive activity for β-arrestin-recruitment to estimate the receptor bias of the L129Q mutation for the Gq/11 and β-arrestin pathways.

To quantify the constitutive activity for β-arrestin-recruitment, we applied a modified version of the Slack-Hall operational model to the BRET^2^ experiments, which enables the calculation of receptor bias comparing Gq/11 and β-arrestin. We noticed that in the absence of a ligand, the Slack-Hall model reduces to the mathematical form of a one-site saturation-binding function ^26^. We performed saturation-binding BRET^2^ experiments ^27^ to measure CysLTR2-GFP10 gene dosage-dependent β-arrestin-RLuc3 recruitment. For each β-arrestin, we compare the L129Q mutant with the wild-type receptor with and without LTD4 stimulation (Suppl. Fig. S3E,F). We use the GFP10 fluorescence in each sample to quantify the total receptor concentration [R]*_t_* ^28^. As compared with the data for β-arrestin1-RLuc3, the fit of the data for β-arrestin2-RLuc3 to LTD4-stimulated CysLTR2-GFP10 wt to a one-site saturation-binding function gives a relatively tight estimate of the dissociation constant *K_d_* of 11065±966 in arbitrary GFP10 fluorescence units, and for the saturating net BRET^2^ ratio (*B_max_*) of 0.0251±0.0009. The gray shaded areas on the graphs indicate the location of the asymmetric 95% confidence intervals for the GFP10 signal of LTD4-stimulated CysLTR2-GFP10 wt giving half-maximal BRET^2^ ratios, as determined by the parameter *K_d_*, and the BRET^2^ ratios for the zero and infinite concentration end points, as determined by the parameters background and *B_max_* (Suppl. Fig. S3E,F). The GFP10 fluorescence of the samples transfected with the highest amount of CysLTR2-GFP10 wt encoding DNA is about twice the *K_d_*. We noticed that the samples expressing CysLTR2-GFP10-L129Q showed smaller GFP10 fluorescence as compared to samples expressing CysLTR2-GFP10 wt, which correlated with increasing degree of cell death for the cells with the oncogenic CAM (Suppl. Fig. S4). Most dead cells are removed by the media change right before the BRET^2^ assay have little impact on the experiment. The cell death induced by the oncogenic CAM most likely results from ERK-mediated apoptosis and autophagy known in HEK239T cells ^29^, which mirrors the ERK-mediated cell proliferation in UVM driven by CysLTR2-L129Q.

Tight independent estimates of *K_d_* and *B_max_* are not required since at low concentrations only the ratio *B_max_*/*K_d_* determines the concentration-dependent binding, which can be estimated from the initial slope. The initial slopes are well defined by samples even at low expression levels of receptors and avoid the need for very high receptor concentrations to reach saturation. This approximation enables a direct comparison of the slopes of the net BRET^2^ ratio versus GFP10 fluorescence data for unstimulated CysLTR2-GFP10 wt and CysLTR2-GFP10-L129Q. Alternatively, one can globally fit the three data sets for each β-arrestin with a shared parameter for *B_max_* to eliminate the effects of the statistical dependence of *B_max_* and *K_d_* in the subsequent calculations of Δlog χ, the differences of log χ for the wild-type and mutant receptors. The resulting *K_d_* values are 117169±12499 for CysLTR2-GFP10-L129Q and 435140±97751 for CysLTR2-GFP10 wt. The resulting Δlog χ is −0.57. We obtained a similar value for β-arrestin1. Compared to the Δlog χ value of −1.22±0.03 for activation of the Gq/11 signaling pathway by the wild-type versus mutant receptor, the double difference ΔΔlog χ shows that the receptor bias of CysLTR2-GFP10-L129Q is 4.5-fold towards Gq/11 activation and away from β-arrestins, suggesting that CysLTR2-L129Q shows high receptor bias for the Gq/11 signaling pathway.

**Fig. 2.**
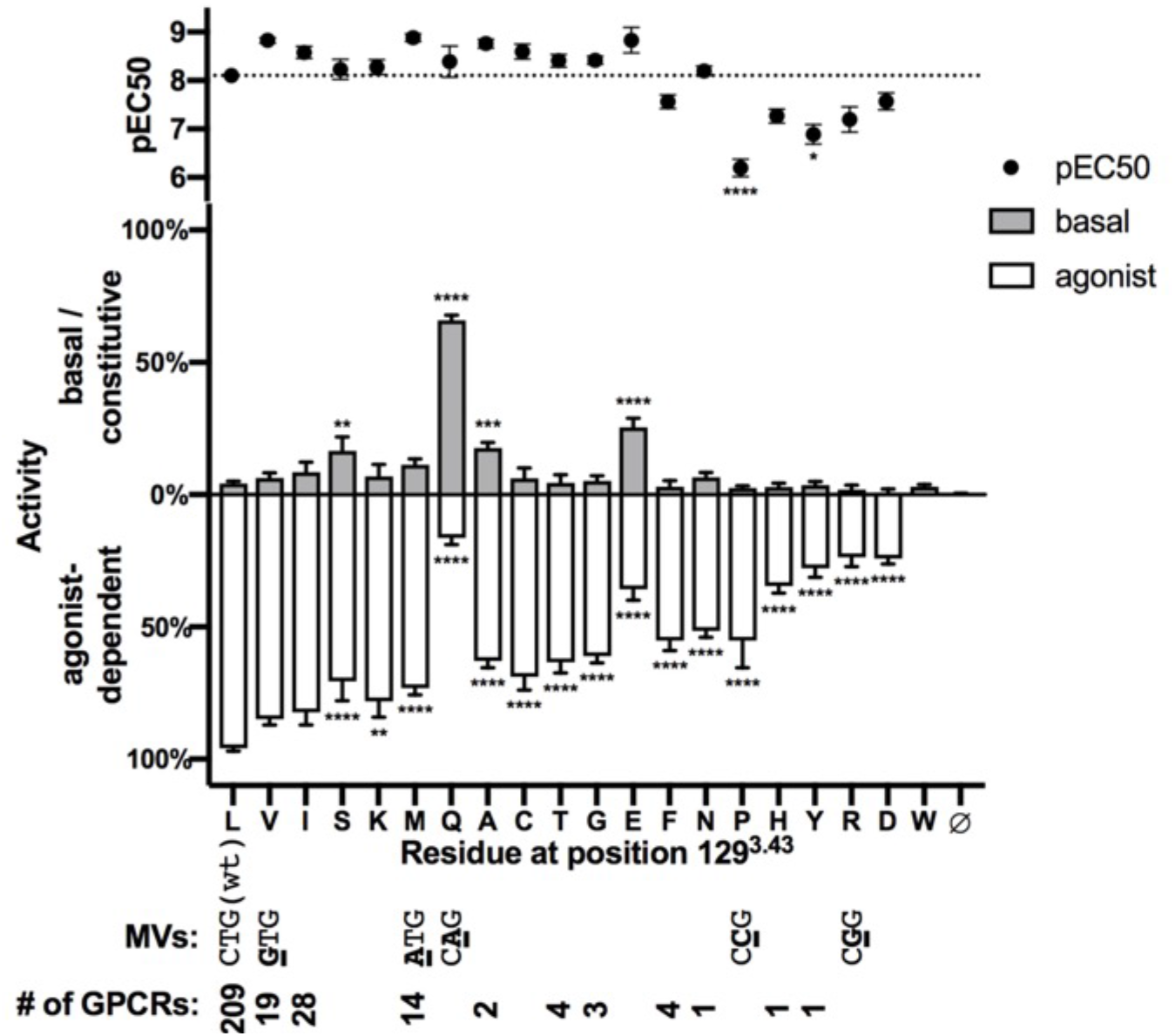
Site-saturation mutagenesis of CysLTR2-L129^3.43^. The basal and agonist-dependent signaling of all 19 natural amino acid variants of residue CysLTR2-L129^3.43^ in HEK293T cells were analyzed. The LTD4 dose-dependent IP1 accumulation for each mutant was fit with a three-parameter sigmoidal function and normalized (cf. Suppl. Fig. S5). We quantify and plot the agonist-dependent activity (white bars) as the span between the normalized bottom and top endpoints, and the basal or constitutive activity (grey bars) as the normalized bottom endpoint. The pEC50 plots (black dots) correspond to the midpoint of the dose-response curve for each mutant. The graph shows the mutants in descending order of total activity defined as the sum of basal and agonist-dependent activity, which corresponds to the height of the white and grey bars combined. The Gln (Q), Glu (E), Ala (A), and Ser (S) variants show significantly increased basal activity. All other residues except Val (V) and Ile (I) in place of Leu (L) show significantly reduced agonist-dependent activity. Pro (P) and Arg (R) show significantly reduced LTD4 potencies (pEC50). The dose-response data are the mean ± SEM from two independent experiments with ten concentrations and five technical replicates, each. The parameters of each mutant are compared with those of the wild-type’s and assessed for significant difference using a two-way ANOVA with Dunnett’s multiple comparison test. The missense variants (MVs) accessible by single-nucleotide exchange from the wild-type CTG codon encode for Val (V, codon GTG), Met (M, codon ATG), Gln (Q, codon CAG), Pro (P, codon CCG), and Arg (R, codon CGG). All other single-nucleotide variants of this codon are synonymous and encode for Leu (L). The conservation of the amino acids at the generic residue position 3.43 can be seen from the number of GPCRs (# of GPCRs) having the particular residue type among a subgroup of 286 rhodopsin-like GPCRs in the GPCRdb.

### Site saturation mutagenesis at position 3.43

To get insight into the role of the glutamine substitution in CysLTR2-L129Q and to study the functional impact of structural variants of the highly conserved residue L^3.43^, we performed a mutational analysis on the constitutive and agonist-dependent activation of the Gq-PLCβ signaling pathway. We determined the LTD4 dose-response in the IP1 accumulation assay for all twenty amino acid variants of position 129^3.43^ in CysLTR2 transiently transfected into HEK293T cells (Suppl. Fig. S5). We analyzed the data with the sigmoidal model and the horizontal line as alternative model (cf. Suppl. Table S1). The basal activity (grey bars, up), the agonist-dependent activity (open bars, down), and the potency (pEC50, black circle) are shown for all variants sorted in descending order of total activity (given as combined height of grey and open bars) (Fig. 2). Compared with the wild-type (leucine, L), all other residues except valine (L) and isoleucine (I) show significantly reduced agonist-dependent activity, most dramatic in the case of tryptophan (W) that showed no agonist-dependent response. Proline (P) and tyrosine (Y) showed significantly lower agonist potencies. Three variants, glutamine (Q), glutamic acid (E), alanine (A), and serine (S) had significantly increased basal activity as compared to wild-type.

### Two-state allosteric model suggests ground state equilibrium of CysLTR2-L129Q is largely shifted to the active state

To quantify the impact of the mutations on the pharmacological observables, we use a two-state allosteric model (Suppl. Scheme S1D) to describe the effect of a mutation as a change of the equilibrium constant *K_q_* for the agonist-independent equilibrium of the receptor with inactive (R_i_) and active (R_a_) states. Our model assumes that the mutation does not change the affinity of the agonist for the inactive state, *K_A_*, and the affinity for the active state *αK_A_*, which are related by the term *α*. Note that in our model the ligand-free active state receptor (R_a_) and the ligand-bound active state receptor (AR_a_) are both transduced into the observable effect E with the same logistic function parameters (*E_max_* and *K_e_*).

We performed a global analysis of the site-saturation mutagenesis of CysLTR2-L129^3.43^ data with the two-state allosteric model (Supple. Fig. S6). Two fitting parameters, *τ* and *K_q_*, are used describe each mutant, and one fitting parameter, α, is used for all mutants. We heuristically fixed the other parameters, *basal*, *E_max_*, and *K_A_*. Note that α and *K_A_* are linearly dependent for *K_A_* below 10^5^, and we fixed *K_A_* to this upper limit. Above that limit, the model fails to describe for mutants with large right-shifted dose-response curves. The resulting fits for a representative set of mutants (Supple. Fig. S6) illustrate the strength of the model. As compared to the empirical sigmoidal model that uses three parameters for each mutant, the two parameters of the two-state allosteric model capture most of the variations in the end and midpoints of the dose-response data. The key advantage of the model is that it enables a thermodynamic relation of receptor pharmacology to structure and dynamics, that is, changes in *K_q_* are directly related to the free-energy difference ΔΔG(active)-ΔΔG(inactive) obtained from the mutational effects on the active and inactive state receptor structures. Moreover, the model is compatible with the extended ternary complex model (Suppl. Scheme S1F) that allows inclusion of further biochemical details, such as the G protein concentration dependency of *K_q_*. The ranking of the constitutive activity of the mutants (Suppl. Fig. S6B) as calculated from the Kq suggests that Q, E, A, M, V, C, I, S, G, T, N, and K all stabilize the active state, whereas F, D, H, R, Y, P, and W all promote the inactive state.

**Fig. 3.**
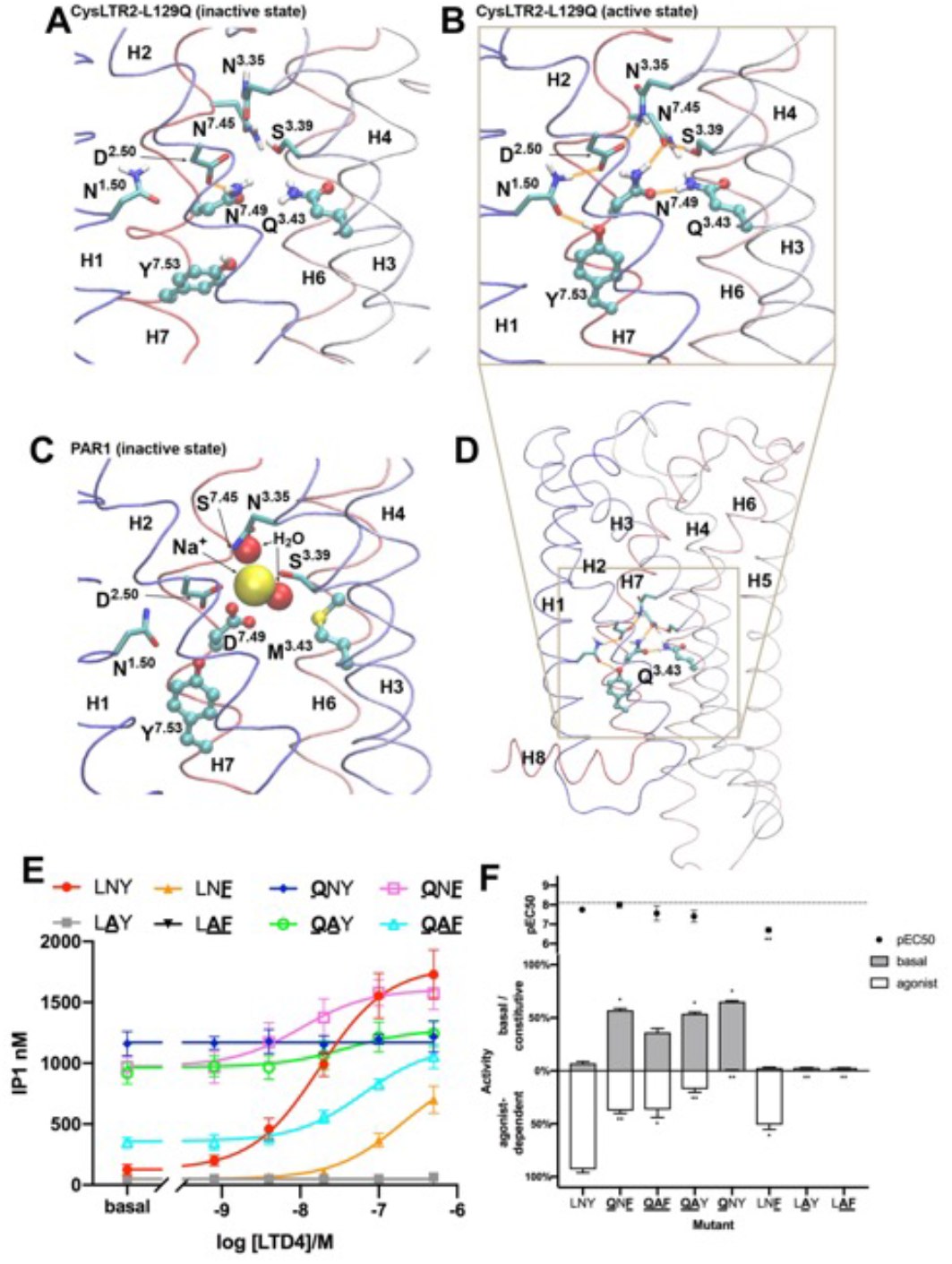
Hydrogen bond network stabilizes active state in CysLTR2-L129Q^3.43^. (**A**) CysLTR2-L129Q mutation modeled in an “inactive state” homology model of CysLTR2 using Rosetta. (**B**) A novel hydrogen-bond network stabilizes the CysLTR2-L129Q in the “active state” structure homology model. CysLTR2-L129Q also disrupts a conserved sodium ion binding site that stabilizes inactive structures of GPCRs. (**C**) Sodium ion binding site in the PAR1 receptor inactive state. (**D**) Overall location of the Q3.43 residue in receptor model. (**E**) The LTD4 dose-dependent IP1 accumulation was assayed for 8 combinations of mutations of 3.43/7.49/7.53. The single mutants (L129Q, **<u>Q</u>**NY; N301A, L**<u>A</u>**Y; Y305F, LN**<u>F</u>**), the double mutants (L129Q/N301A, **<u>QA</u>**Y; L129Q/Y305F, **<u>Q</u>**N**<u>F</u>**; N301A/Y305F, L**<u>AF</u>**) and the triple mutant (L129Q/N301A/Y305F, **<u>QAF</u>**) at 3.43 and 7.49 and 7.53 of the NPxxY motif were compared with wild-type (LNF). (**F**) Bar graph showing the basal (grey bars) and agonist-dependent activity (white bars) normalized relative to maximally stimulated wild-type receptor and mock transfected control. The LTD4 potencies (pEC50) are shown as points. The results from the 8 variants are ordered in descending order of total activity (basal plus agonist-dependent) corresponding to the combined height of the grey and white bars. The data are the mean ± SEM from two independent experiments with six concentrations and four technical replicates, each. The obtained parameters of each mutant are compared to those of the wild-type’s in a two-way ANOVA with Dunnett’s multiple comparison test.

### Molecular activation mechanism of CysLTR2 mutations at position 3.43

The mutation CysLTR2-L129Q is located at the generic position (3.43) following the GPCRdb/Ballesteros-Weinstein numbering system. Position (3.43) is highly conserved with 96% hydrophobic residues (Leu, Ile, Val, Met, and Phe) in 286 class A GPCRs from the GPCRdb (cf. bottom panel of Fig. 2). L(3.43)Q mutations have been shown to induce disease-causing constitutive activity in CysLTR2 ^6^ and thyroid stimulating hormone receptor (TSHR) ^30^. Other constitutively activating mutations of L(3.43) have been described for the luteinizing hormone (LH)/chorionic gonadotropin (CG) receptor (LHCGR), follicle stimulating hormone receptor (FSHR), β_2_-adrenergic receptor (β_2_-AR), and M_1_ acetylcholine receptor (M_1_AChR) ^13, 31, 32^.

To elucidate the molecular mechanism of constitutive activation due to L(3.43)Q mutations, we used an *in silico* computational approach. Crystal structures of CysLTR2 are not yet available. Instead, we decided to use homology models of CysLTR2 in the active and inactive state. Ideally, to generate homology models for multiple conformational states, the structural templates for each state should come from structures of one particular GPCR crystallized in two conformational states, and that GPCR should have high sequence homology with CysLTR2. However, structures of GPCRs in the active and inactive states have been solved for only a handful of receptors, viz., μ- and κ-opioid, NTS1, CB_1_ cannabinoid, β_2_- and β_1_- adrenergic, M_2_ acetylcholine, A_2A_- and A_1_-adenosine, 5-HT_2C_ receptors, and rhodopsin. These receptors with known active and inactive state structures are more phylogenetically distant to CysLTR2, and none of these receptors is from the δ-branch of rhodopsin-like GPCRs as determined by phylogenetic analysis ^33^, which still has a poor structural coverage. Alternatively, we could use receptors with known structures that are more closely related to CysLTR2, but where either only an inactive or an active structure has been solved. Homology models based on those receptors might give a better idea about potential changes in helix packing of δ-branch rhodopsin-like class A receptors.

We decided to use the active and inactive state homology models of CysLTR2 from the GPCRdb ^34^, which are based on structural templates from the structures of the C-C chemokine receptor 5 (CCR5) for the inactive state ^35^ and the lysophosphatidic acid receptor 6 (LPA6) for the active state ^36^. We introduced the L129Q^3.43^ mutation with the Rosetta software to optimize the resulting models by local side-chain repacking and constrained energy minimization, and to predict the difference in free energies ΔΔG of the wild-type and mutant structures ^37^. In addition, we prepared models of the L(3.43)Q mutation introduced in all the GPCRs with known active and inactive state structures. Next, we analyzed the packing around the L129Q mutation.

### A novel hydrogen-bond network stabilizes the CysLTR2-L129Q active state structure

We reasoned that as an activating mutation, it should *stabilize the active state* and/or *destabilize the inactive state*. We observed in models of several GPCRs that the L(3.43)Q side chain is in hydrogen-bonding distance of the side chains of N(7.49) or Y(7.53) in the highly conserved NPxxY motif. We only see these interactions in models based on templates in the active state, but not in the inactive state. This finding is highly relevant as a potential mechanism for a CAM. The hallmark feature of GPCR activation is the outward movement of the cytoplasmic end of helix 6. Less prominently featured are concomitant changes with an inward movement of the cytoplasmic end of helix 7. We hypothesize that L(3.43)Q stabilizes the active state receptor by direct interaction with NPxxY. We predict that many class A GPCRs will be activated by the L(3.43)Q mutation, given the high conservation of the conformational change and the residues involved in this interaction, and further supported by the complete absence of a glutamine residue at the 3.43 position in all known GPCR sequences. However, other receptors might lack the receptor bias away from β-arrestin and the efficiency of β-arrestin-dependent desensitization and downregulation mechanisms will determine the ultimate phenotype of the mutation.

The active and inactive state models of CysLTR2-L129Q (Fig. 3) reveal an extended hydrogen bond network linking Q^3.43^ to N^7.49^, N^7.49^ to N^7.45^, N^7.45^ to S^3.39^, N^3.35^ to D^2.50^, D^2.50^ to N^1.50^, and N^1.50^ to Y^7.53^ exclusively present in the active state (Fig. 3B). None of these residues are hydrogen bonded to each other in the inactive state (Fig. 3A). Comparison with the inactive state structure of PAR1 ^38^ shows that most of these residues are involved in binding of the hydrated sodium ion (Fig. 3C).

### CysLTR2-L129Q also disrupts a conserved sodium ion binding site that stabilizes inactive structures of GPCRs

Residue L129^3.43^ is part of the allosteric sodium-binding site ^39^. Negative allosteric modulatory (NAM) effects of sodium ions and amilorides have been observed for some class A GPCRs. Amilorides are analogs of the diuretic amiloride, and they are small organic cations that can bind in the sodium-binding site ^40^. The sodium-binding site collapses upon receptor activation, and the NAM effect of the sodium ions and amilorides can be explained by selectively stabilizing the inactive state.

Our alternative hypothesis is that L129Q^3.43^ blocks sodium binding and destabilizes the inactive state. It is currently unknown if CysLTR2 is controlled by sodium ions, but the conservation of the residues suggests that the receptor has a functional sodium-binding site. The only crystal structure of a δ-group GPCR with clear evidence of a bound sodium is PAR1 (Fig. 3C), which has D^7.49^ as an additional acidic residue in the sodium pocket ^38^. Note that CysLTR1 also has D^7.49^, but CysLTR2 has the common N^7.49^. We further speculate that the sodium-binding pocket might accommodate a small molecule drug specific for CysLTR2-L129Q and virtual screening of compound libraries should be possible once a high-resolution crystal structure of CysLTR2 becomes available.

### Reciprocal mutagenesis of positions L^3.43^, N^7.49^, Y^7.53^ (L129 and the conserved NPxxY motif)

Our homology models predicted novel interactions, such as the hydrogen-bonding interactions between L129Q^3.43^ and N301^7.49^ and potentially between L129Q^3.43^ and Y305^7.53^. To test the role of these interactions, we investigated these interactions by mutagenesis using single, double and triple mutants in an effort to disrupt a particular stabilizing interaction and measure the functional outcome. We focused on two variants for each site, the wild-type residue together with one mutation. We used leucine (L) and glutamine (Q) at position 129^3.43^, asparagine (N) and alanine (A) at position 301^7.49^, and tyrosine (Y) and phenylalanine (F) at position 305^7.53^. We designate the eight possible combinations by three letters highlighting the mutated residues as LNY, LN**<u>F</u>**, L**<u>A</u>**Y, L**<u>AF</u>**, **<u>Q</u>**NY, **<u>Q</u>**N**<u>F</u>**, **<u>QA</u>**Y, and **<u>QAF</u>**. The LTD4 dose-response for each of these mutants in the IP1 accumulation (Fig. 3D) compared to the wild-type (LNY) sample shows a significant loss of agonist-dependent activity for all mutants, a significant gain of constitutive activity for **<u>Q</u>**N**<u>F</u>**, **<u>QA</u>**Y, and significantly lower pEC50 for **<u>Q</u>**NY (Fig. 3E). Interestingly, introducing either N301A, Y305F, or N301A/Y305F into the L129Q mutant partially reverts the Loss-of-Function in agonist-dependent signaling, but without reverting the Gain-of-Function in basal activity. The same mutations introduced into the wild-type receptor let to a Loss-of-Function in agonist-dependent signaling, which was partial for Y305F and complete for N301A and N301A/Y305F. Together, these findings underline the importance of the highly conserved NPxxY motif in active state formation.

### Screening for additional oncogenic driver mutations in CYSLTR2

More than 200 variants of uncertain significance (VUS) are known for *CYSLTR2.* The GPCRdb lists 119 germline missense variants (MVs) for *CYSLTR2*. The COSMIC database lists 76 somatic MVs from tumor samples, and *The Cancer Genome Atlas (TCGA)* lists 79 somatic MVs. To identify other potentially oncogenic MVs, we compared the two sets of somatic MVs with the set of germline MVs. We found in total 218 non-redundant MVs in all three databases, with 98 germline-only MVs (“normal”, **Suppl. Tab. S5**), 18 germline and somatic MVs (“both”, **Suppl. Tab. S6**), and 102 somatic-only MVs (“cancer”, **Suppl. Tab. S5**). Interestingly, we found 18 new recurrent or “hotspot” mutations when merging the somatic-only MVs from COSMIC and TCGA. Next, we annotated MVs with the generic modified Ballesteros-Weinstein numbering system used in the GPCRdb for structural comparison with homologous GPCRs in order to identify functionally relevant sites. Here we found that many of the “cancer” MVs are located in functionally relevant sites, such as, the sodium ion pocket (Na), the microswitch (MS), and the G protein binding interface (GP) ^3^. Next, we performed a virtual phenotypic screening of a GPCR cancer genomics database *in silico* to identify potentially activating MVs.

**Fig. 4.**
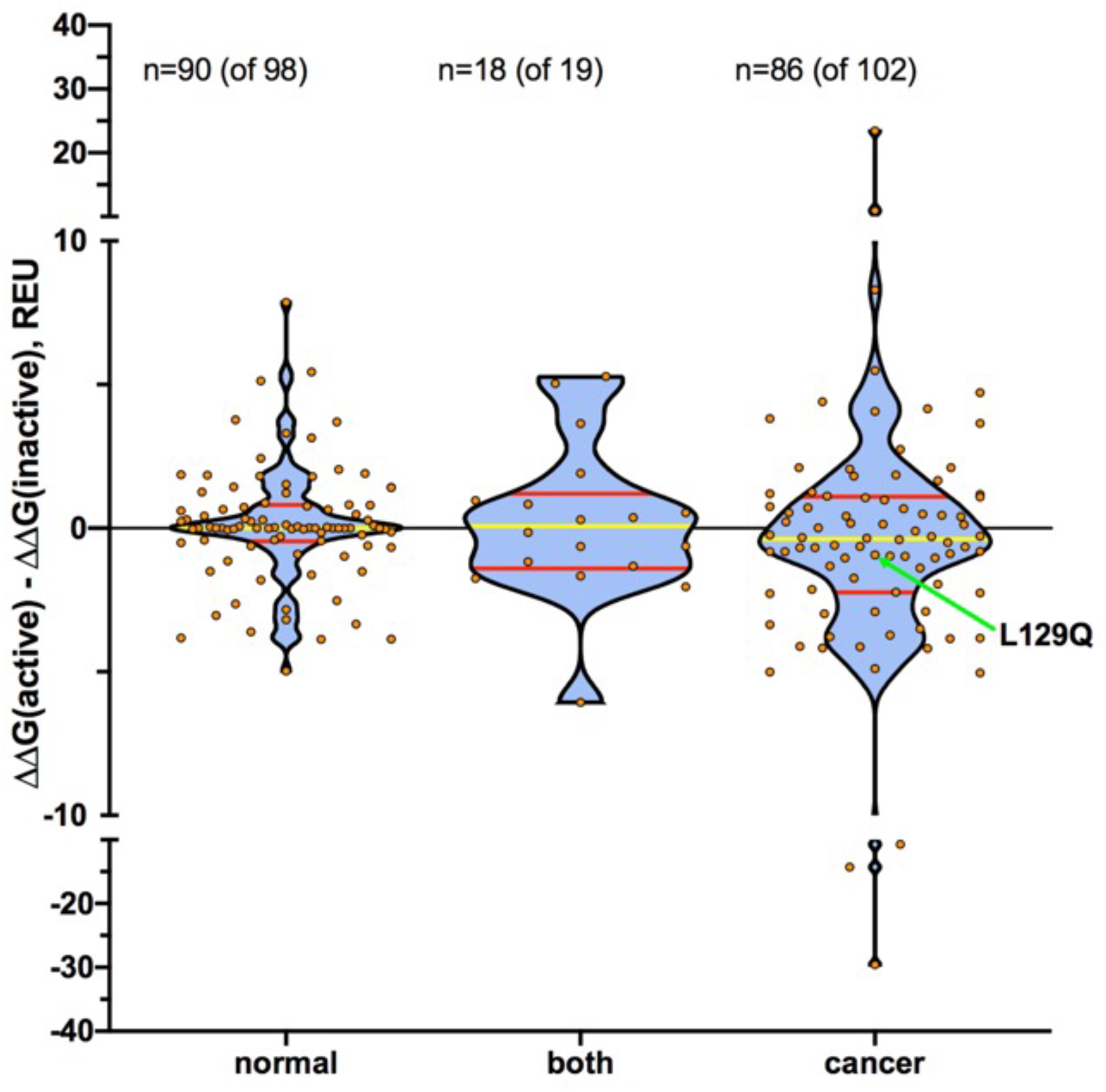
Prediction of active state stabilization in CysLTR2 MVs from cancer genomes. More than 200 variants of uncertain significance (VUS) are known for *CYSLTR2*. We used Rosetta for a virtual screening of the free energy change of the mutations in active relative to inactive state structural models of CysLTR2. Negative values predict stabilization of the active state conformation. Each data point corresponds to one mutation. The normal group are variants only known as germline mutations, whereas the cancer group are variants exclusively found as somatic mutations in cancer samples. The remaining variants are found both as germline and somatic variants.

### Rosetta structural modeling to predict active state stabilization of VUSs

To predict the structural and energetic impact of *CYSLTR2* MVs, we used the Rosetta ddG algorithm to model the structure and to predict the free energy change (ΔΔG) of 194 different amino acid substitutions encoded by the 194 MVs that map to residues included in the CysLTR2 homology models. The algorithms used by Rosetta are easily scalable to analyze large sets of mutations, and they provide a good balance of computational cost-to-performance ratio. We calculated the active state stabilization (ΔΔΔG) as the free-energy difference ΔΔG(active)-ΔΔG(inactive) obtained from the mutational effects on active and inactive state structures (cf. Suppl. Tables S5-S7). A negative value indicates stabilization of the active state. The distributions of active state stabilization (ΔΔΔG) are illustrated as violin plots for “normal”, “both”, and “cancer” MVs (Fig. 4). The free energies are given in Rosetta Energy Units (REU). Interestingly, the “normal” MVs have little impact on active state stabilization and most data points are tightly clustered around zero, which is also reflected in the locations of the median value (0.003 REU) and the 25^th^ percentile (−0.459 REU) and 75^th^ percentile (0.807 REU) tightly around zero. MVs in the “both” groups have a median at 0.290 REU and a slightly wider distribution with the 25^th^ percentile at 1.320 and the 75^th^ percentile at 1.060. In contrast, the “cancer” MVs show a much wider distribution (25% percentile at −2.240 REU and 75% percentile at 1.105 REU) that is shifted to negative with a median value of −0.400 that indicates the majority of these variants stabilize the active state.

**Fig. 5.**
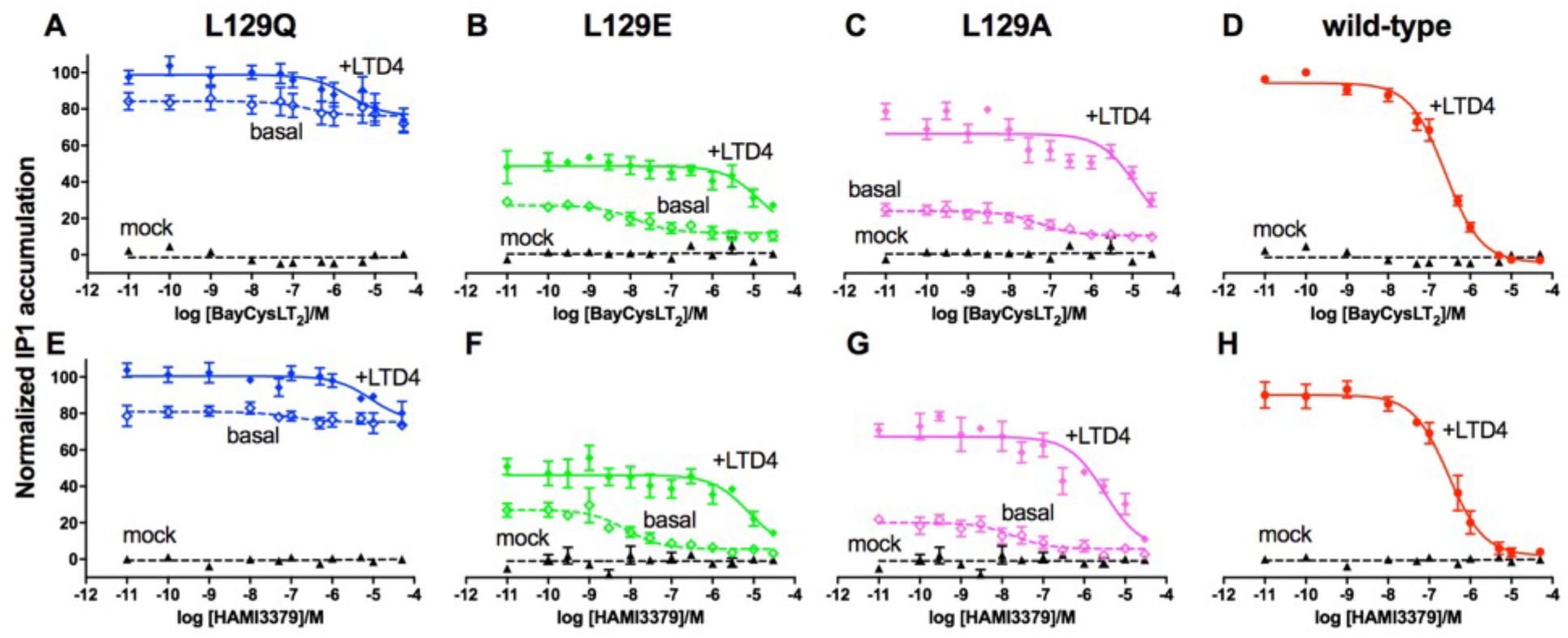
CysLTR2 antagonists BayCysLT2 and HAMI3379 have limited efficacy as inverse agonists. The CysLTR2 antagonists BayCysLT2 (**A-D**) and HAMI3379 (**E-H**) and show only minimal reduction of the constitutive activity of the three CAMs, L129Q, L129E, and L129A (basal, dashed lines and open symbols), and full inhibition of the LTD4-dependent IP1 stimulation of the CAMs and wild-type receptor (+LTD4, solid lines and symbols). CysLTR2-L129Q (open blue squares). The data are presented as the percentage of IP1 accumulation minus empty vector over the maximal response exhibited by CysLTR2 wt following 100 nM LTD4 stimulation and represent the mean ± SEM of at least 3 independent experiments, each carried out in at least triplicate.

### Known CysLTR2 antagonists have limited efficacy as inverse agonists

To test if known antagonists at CysLTR2 have inverse agonist activity, we compared the antagonist dose-dependent effects on IP1 accumulation in cells expressing CysLTR2-L129Q, -L129E, and - L129A. BayCysLT2 (Fig. 5A-C) and HAMI3379 (Fig. 5E-G) both result in small reduction of the basal signaling of the three CAMs with larger effects on the weaker CAMs as compared to L129Q. Both compounds also inhibit the LTD4-dependent increase in IP1 accumulation for the CAMs and the wild-type receptor (Fig. 5D,H). We have also used a modified two-state allosteric model including two competing ligands (Suppl. Scheme S1E). A global fit of the model suggests that HAMI3379 slightly stronger stabilizes the inactive state (with log β = −0.43±0.05) and is more potent (with log K_b_ = 7.63±0.08) as compared to BayCysLT2 (log β = −0.30±0.05 and log K_b_ = 7.17±0.09). Together, these findings suggest that the two CysLTR2 antagonists HAMI3379 and BayCysLT2 act as neutral antagonists, but they have limited efficacy as inverse agonists targeting the oncogenic CAM CysLTR2-L129Q.

### Oncogenic signaling pathways in uveal melanoma

**Fig. 6.**
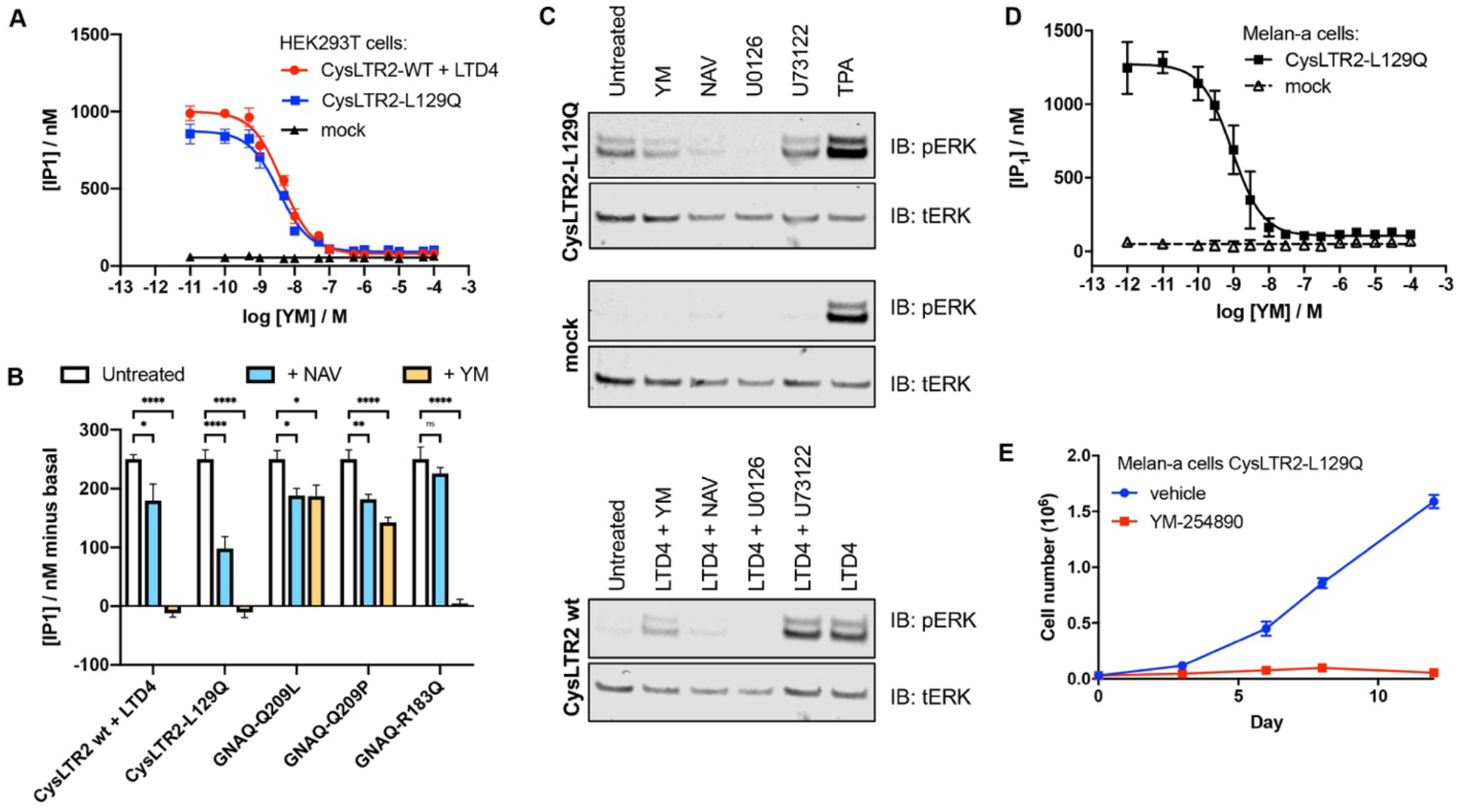
Targeting the signaling of CysLTR2-L129Q and GNAQ CAMs with Gq- and Arf6-inhibitors. **(A)** Dose-dependent effect of YM-254890 on IP1 accumulation comparing HEK293T cells expressing CysLTR2-L129Q with LTD4-stimulated CysLTR2 wt and mock-transfected cells. **(B)** Effects of YM and NAV on the PLCβ-dependent signaling of different constitutively active mutations (CysLTR2-L129Q, Gq-Q209L, Gq-Q209P, and Gq-R183Q) and LTD4-stimulated CysLTR2 wt transiently transfected in HEK293T cells. (**C**) ERK/MAPK activation in HEK293T cells transfected with CysLTR2-L129Q as compared to mock transfected controls and LTD4-stimulated wild-type receptor. We tested inhibitors for Gq/11 (YM), Arf6 (NAV), MEK (U0126), PLC (U73122) on phosphor-ERK (pERK) formation indicating ERK activation. Total ERK (tERK) serves as a loading control. (**D**) YM-254890 dose-response on IP1 accumulation in melan-a melanocytes stably expressing the receptor mutant expressing CysLTR2-L129Q. (**E**) Inhibitory effect of YM-254890 on the growth of melan-a expressing CysLTR2-L129Q.

#### Targeting CysLTR2 and GNAQ CAM signaling pathways with Gq (YM) and Arf6 (NAV) inhibitors

To identify suitable drugs to target the downstream signaling of the oncogenic CysLTR2-L129Q CAM, we first investigated the effect of the Gq/11 inhibitor YM-254890. YM-254890 is known to block LTD4-stimulated calcium signaling of CysLTR2 ^41^. We hypothesized that CysLTR2-L129Q couples to Gq/11 and inhibition of Gq/11 blocks activation of the downstream effector PLC-beta. We tested the dose-dependent effect of YM-254890 on IP1 accumulation comparing cells expressing CysLTR2-L129Q with LTD4-stimulated CysLTR2 wt and mock-transfected cells (Suppl. Figure S7, Fig. 6A). The Gq/11 inhibitor completely blocks the CysLTR2- and CysLTR2-L129Q-dependent IP1 accumulation. Therefore, both CysLTR2 wt and CysLTR2-L129Q signal through Gq/11 activation to PLC-beta.

We next tested the effect of the Arf6 inhibitor NAV-2729. Inhibition of Arf6 by NAV-2729 has been shown to block all known signaling pathways (PLC/PKC, Rho/Rac, YAP, and beta-catenin) of oncogenic Gq/11 and it drives a redistribution of Gq/11 from cytoplasmic vesicles to the plasma membrane ^42^. Arf6 is activated in GEP100-dependent manner by Gq/11 and drives internalization and signaling of Gq/11-PLC from endosomal compartments, and inhibition of Arf6 reduces Gq/11-PLC signaling to the contribution from the plasma membrane compartment. We hypothesized that Arf6 is involved in Gq/11-dependent signaling of CysLTR2-L129Q. We analyzed the dose-dependent effect of NAV-2729 on CysLTR2-L129Q-dependent IP1 accumulation (see Suppl. Figure S7). The compound results in almost complete inhibition of IP1 accumulation, only limited by the practical upper limit of the concentration in the assay of ten micromolar. We conclude that Gq/11, Arf6 and the Arf6-GEF GEP100 are potential therapeutic targets for CysLTR2-L129Q-driven UVM.

To compare the effects of YM and NAV on the PLC-beta-dependent signaling of different driver mutations found in UVM and related conditions, we tested a fixed concentration of each inhibitor as a function of the gene dosage for a set of constitutively active mutations (CysLTR2-L129Q, Gq-Q209L, Gq-Q209P, and Gq-R183Q) and LTD4-stimulated CysLTR2 wt transiently transfected in HEK293T cells. The gene dosage-dependent IP1 accumulation was well approximated by as a semilogarithmic function in the range of 0.14ng to 11ng plasmid DNA. To compare the different genes at a comparable overall signaling level, we interpolated the gene dosage-dependent IP1 accumulation data to 250 nM IP1 for the untreated samples. We then used the interpolated value of the gene dosage to obtain the IP1 accumulation for the samples treated with NAV and YM. The results are shown as a bar graph (Fig. 6B). Interestingly, treatment of the different samples with NAV indicated that the contribution of Arf6 activation to the observed IP1 accumulation differs substantially. NAV has the largest effect on CysLTR2-L129Q-dependent signaling, whereas the effect on the signaling from LTD4-stimulated CysLTR2 was much smaller. The two oncogenic mutants, Gq-Q209L and Gq-Q209P, show a comparable reduction of IP1 accumulation with NAV, but the small reduction we see for the driver mutant Gq-R183Q in benign melanocytic tumors is insignificant. Next, we compared the effects of YM. The results show that the IP1 accumulation from LTD4-stimulated CysLTR2 wt, and from the constitutive activity of CysLTR2-L129Q and Gq-R183Q is fully inhibited by YM-254890, whereas only incomplete inhibition can be obtained in samples expressing Gq-Q209L and Gq-Q209P, consistent with earlier reports on Gq-Q209L and Gq-R183C ^41^. We conclude that CysLTR2-L129Q is more sensitive to the Gq/11 and Arf6 inhibitors than any of the Gq driver mutants tested in HEK293T cells.

Next, we characterized the activation of the mitogen-activated protein kinase (MAPK)/extracellular signal-regulated kinase (ERK) pathway by the oncogenic CAM CysLTR2-L129Q. Activation of the ERK/MAPK pathway plays a pivotal role in uveal melanoma^6^. We tested in HEK293T cells the inhibitory effects of the Gq/11 inhibitor YM-254890, the Arf6-inhibitor NAV-2729, the MEK inhibitor U0126, and the PLC inhibitor U73122 (Fig. 6C). Stimulation of PKC with the tumor-promoting phorbol ester, 12-O-tetradecanoylphorbol-13-acetate (TPA), results as expected in ERK activation and served as positive control. We found that YM, NAV and U0126 strongly inhibit the activation of ERK by the oncogenic CAM as seen by a reduction of phosphor-ERK (pERK) compared to the untreated control.

We next tested the effects of YM in a UVM model based on melan-a cells. Melan-a cells, a murine melanoblast line, require TPA for growth. CysLTR2-L129Q, but not wild-type CysLTR2, stably expressed in melan-a cells can drive TPA-independent cell proliferation in vitro and tumor formation in vivo ^6^. The tumor-promotor TPA mimics the action of DAG and activates protein kinase C (PKC), which in turn can stimulate the ERK/MAPK pathway that drives melanocyte proliferation. DAG and IP3, indirectly through Ca2+, stimulate PKC. DAG and IP3 are the second messengers generated by PLC. We hypothesized that Gq/11-dependent activation of PLC-beta by CysLTR2-L129Q stimulates PKC, and results in PKC-dependent ERK/MAPK signaling and cell proliferation. In UVM cells, PKC phosphorylates the Ras guanylyl-releasing protein 3 (RasGRP3) and activates the Ras–Raf–MEK–ERK pathway. The melan-a cell model recapitulates the role of PKC-dependent activation of RasGRP3 as the link of Gq/11 signaling to ERK activation normally found in UVM cells. We observed YM-254890 dose-dependent inhibition of the IP1 accumulation in melan-a cells stably transduced with CysLTR2-L129Q (Fig. 6D). Moreover, YM-254890 efficiently suppresses TPA-independent growth of these cells (Fig. 6E). We conclude that blockage of the CysLTR2-L129Q-dependent activation of Gq/11 is necessary and sufficient to suppress phorbol ester-independent melanocyte proliferation. Therefore, Gq/11 is a suitable therapeutic target for UVM driven by CysLTR2-L129Q.

## DISCUSSION

UVM is the most common intraocular malignancy and is associated with a high rate of metastasis with short survival time for patients, the liver being the most common site for secondary tumors. A hallmark feature of UVM is an aberrant activation of Gq protein-dependent signaling cascades. The signaling pathways relevant for UVM (Scheme 1) suggest potential targets for pharmacotherapy of the disease. The CAM in the receptor CysLTR2 drives the formation of the active state (R*). One of our goals is to identify an inverse agonist, a small molecule drug that blocks the receptor activation. The activation of the G protein (Gαq•GDP•βγ) by R* results in nucleotide exchange and dissociation to give Gαq•GTP and βγ. In tumors driven by CAMs of Gαq, the activation occurs independently of the receptor. GPCR kinase (GRK) is an effector of Gαq•GTP and mediates formation of a phosphorylated receptor (R*-P). β-arrestin (βArr) binds R*-P and results in the desensitized receptor–β-arrestin complex (R*-P• βArr). The receptor-β-arrestin binds the adapter protein 2 (AP2) and targets the complex to clathrin-coated pits for internalization to endosomal vesicles. Analogous clathrin-independent internalization pathways are not shown. Depending on the ubiquitination of the receptor and β-arrestin, the endosomal sorting results in recycling to the cell surface or degradation ^43, 44^. Little is known about the β-arrestin-dependent trafficking of CysLTR2. Our observation of receptor bias away from β-arrestin recruitment suggests that the oncogenic CAM CysLTR2-L129Q escapes β-arrestin-dependent downregulation. We envision future antibody-based therapies against CysLTR2-L129Q that stimulate receptor internalization and downregulation.

**Scheme 1.**
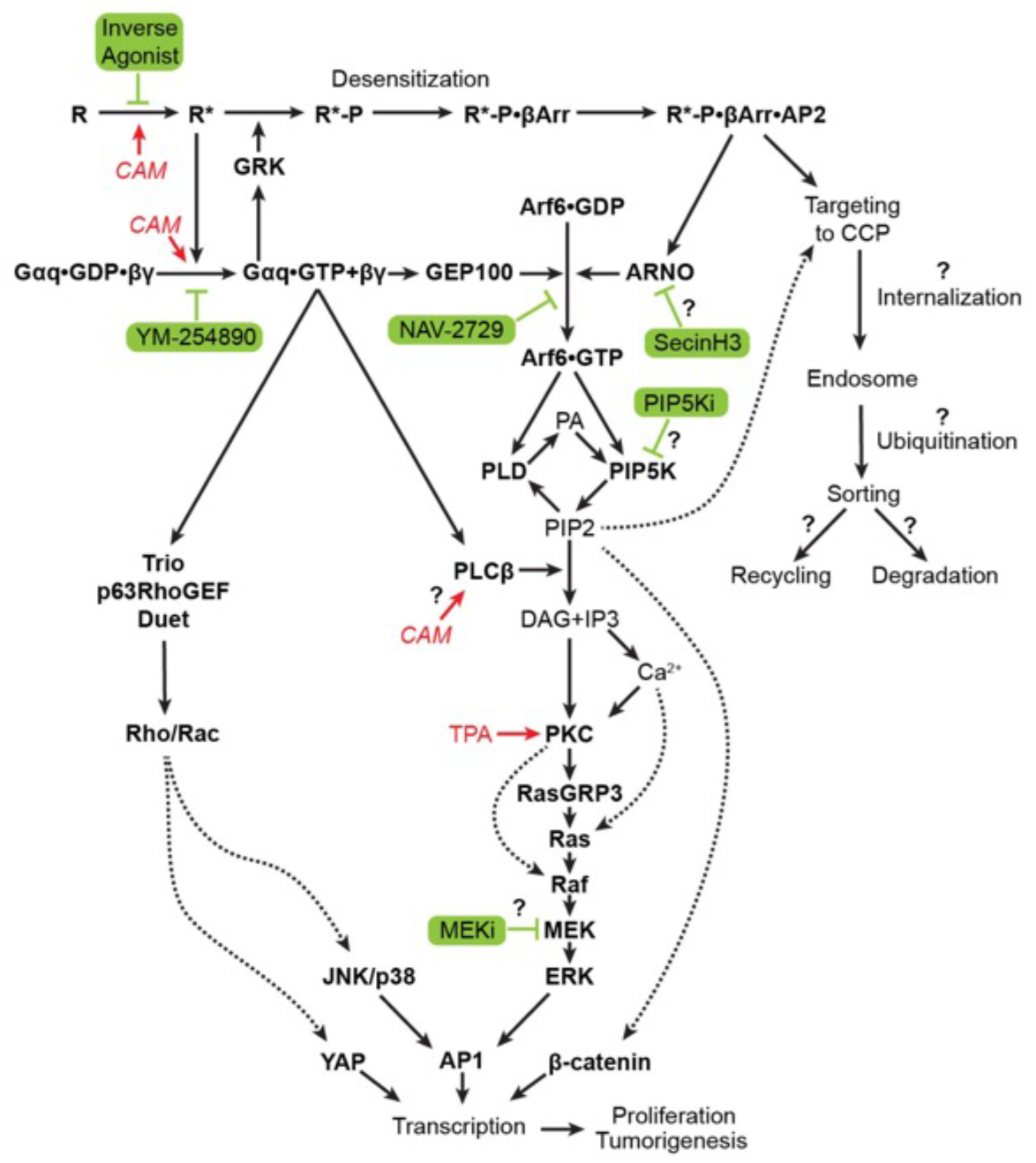
Targeting the oncogenic signaling pathways in uveal melanoma. Constitutively activating mutations (CAMs) of CYSLTR2, GNAQ and GNA11 result in strong stimulation of second messengers (DAG, IP3, and Ca^2+^).

While GRK is an effector of Gαq•GTP with negative feedback on the input from the activated receptor, the other effectors of Gαq•GTP stimulate signal transduction pathways culminating in activation of transcriptional programs for proliferation and tumorigenesis. Phospholipase C-β (PLC-β) is the classical effector of Gq/11. It has a high catalytic activity to cleave the substrate phosphatidylinositol 4,5-bisphosphate (PIP2) into the second messengers, diacylglycerol (DAG) and inositol 1,4,5-trisphosphate (IP3). It has been suggested that PLC-β-dependent IP1 accumulation also includes contributions from the IP2 formed by direct hydrolysis of PI4P, instead of IP2 formed by the dephosphorylation of IP3 formed by hydrolysis of PIP2 ^45^. Since this novel pathway does not depend on PIP2, which is predominantly found at the plasma membrane, the PI4P-dependent IP1 accumulation could have a substantial contribution to PLC-β signaling from the endosomal compartment. ADP-ribosylation factor-(Arf) Guanine Exchange Protein 100-kDa (GEP100) is a guanine-nucleotide exchange factor (GEF) that promotes binding of GTP to the Arf6. Arf6•GTP stimulates phospholipase D (PLD) and phosphatidylinositol 4-phosphate 5-kinase (PIP5K) ^46^. PLD uses abundant lipids to generate phosphatidic acid (PA) ^47^. PIP5K synthesizes PIP2 from the precursor phosphatidylinositol 4-phosphate (PIP). PIP2 is only a minor lipid component in the cell and its supply limits the second messenger output from Gαq•GTP–PLC-β. We hypothesize that the key function of the Arf6 in UVM is to ensure ample supply of the PLC-β substrate PIP2. Our experiments show additive effects of the Gq/11 inhibitor YM-254890 ^41^ and the Arf6 inhibitor NAV-2729 ^42^ abolishing the CysLTR2-L129Q-dependent IP3 formation. We predict that the Gq/11 inhibitor will also abolish binding of its effectors Trio, p63RhoGEF, and Duet/Kalirin, which activate Rho/Rac-dependent JNK/p38 and YAP pathways ^48,49,50^. Arf6 signaling through its effector PIP2 is also required for clathrin-dependent and -independent endocytosis ^51^ of receptor–β-arrestin complexes, which in turn stimulate PIP2 formation through the Arf6-GEF ARNO ^52, 53^. PIP2 is also required for β-catenin signaling, which is another oncogenic pathway relevant to UVM ^54^. We predict that a PIP5K inhibitor (PIP5Ki) will block CysLTR2-L129Q-dependent signaling.

Protein kinase C (PKC) is stimulated by DAG and indirectly through Ca^2+^ by IP3. The tumor promoter 12-O-tetradecanoylphorbol-13-acetate (TPA) targets PKC and stimulated the mitogen-activated protein kinase (MAPK)/extracellular signal-regulated kinase (ERK) pathway. In UVM cells, PKC phosphorylates the Ras guanylyl-releasing protein 3 (RasGRP3) and activates the Ras–Raf–MEK–ERK pathway ^55^. Activation of this pathway is essential for growth of UVM cells, and also for the TPA-dependent growth of melan-a cells. Other cell lines, such as HEK293T, utilize different Ca^2+^- and PKC-dependent mechanisms to activate ERK ^56^. Some GPCRs stimulate ERK through β-arrestin-mediated signaling complexes (“signalosomes”) ^57, 58^.

Although the CysLTR2-L129Q CAM is highly biased toward Gq signaling, we show that CysLTR2-L129Q does recruit β-arrestin, albeit much less efficiently than the agonist-stimulated wild-type receptor. It is possible that the β-arrestin-bound receptor forms ERK-activating signalosomes. Since certain GRKs can mediate G protein-independent receptor phosphorylation and β-arrestin-recruitment, it would be possible that ERK-activating signalosomes might result in G protein-independent activation of ERK. Such a mechanism explains the incomplete suppression of CysLTR2-L129Q-dependent ERK activation using the Gq/11 inhibitor YM-254890, while inhibitors of Arf6 and MEK both showed stronger reduction of pERK as compared to the Gq/11 inhibitor in HEK293T cells. Therefore, we propose that MEK inhibitors (MEKi) might be useful in addition to Gq/11 and Arf6 inhibitors in targeting CysLTR2-L129Q signaling. Such combination therapy might require much lower dosage as compared with monotherapy and reduce side effects.

The discovery of GPCR CAMs as potential driver oncogenes suggests that the study of CAM hotspot mutation sites and the mapping of gene overexpression in tumor samples might be able to identify GPCRs as novel cancer therapeutic targets. The observation that known neutral antagonists might also have potential inverse agonist activity toward CAMs, also suggests that existing drugs might be able to target GPCR CAMs in human cancers. Thus, novel therapeutic strategies targeting specifically CAM GPCRs could benefit patients who are being treated according to the genetic signatures in their tumor.

In conclusion, we characterized a *CYSLTR2* GPCR oncogene in UVM and established the proof-of-principle that an inverse agonist should able to inhibit the persistent signaling from the CysLTR2 L129Q CAM. We establish that the CysLTR2 L129Q CAM is highly biased toward Gq/11 cellular signaling pathways and fails to recruit significantly β-arrestin. The constitutively biased signaling pattern of CysLTR2 L129Q explains why it can persistently activate Gq and avoid cellular downregulation mechanisms. Furthermore, we provide a structural model showing how the mutation at position 3.43 facilitates the formation of novel hydrogen-bonds through the conserved NPxxY motif that stabilizes an activated state of CysLTR2, while at the same time disrupts a conserved sodium binding site that would normally be expected to stabilize the inactive state of the receptor. Finally, we show how the structural model can be used to predict the functional effects of other mutations found in GPCRs in tumor databases.

## Acknowledgments

We thank the Rockefeller University High-Throughput and Spectroscopy, Bio-Imaging, and Flow-Cytometry Resource Centers for training, discussions, and assistance with the experiments. E.C. was funded by the François Wallace Monahan Fellowship and the Tom Haines Fellowship in Membrane Biology. M.H., J.M.M. were supported by the Tri-institutional Training Program in Chemical Biology, and M.H. by T32 GM115327. We also acknowledge grant support from the Robertson Foundation, the Crowley Family Fund, and the Danica Foundation to T.H. The work was further supported by MSKCC Support Grant/Core Grant (P30 CA008748) and grants from the NCI (K08CA140946 Y.C.; R01CA193837, Y.C.; P50CA092629, Y.C.; P50CA140146, P.C.; K08CA151660, P.C.; DP2 CA174499, P.C.), US DOD (W81XWH-10-1-0197, P.C.), Prostate Cancer Foundation (Y.C.), Starr Cancer Consortium (Y.C., P.C.), Geoffrey Beene Cancer Research Center (Y.C., P.C.), Gerstner Family Foundation (Y.C.), Bressler Scholars Fund (Y.C.), Cycle for Survival (Y.C.).

## Detailed Materials and Methods

### Materials

HAMI3379, BayCysLT2 and LTD4 were from Cayman Chemical (Ann Arbor, MI). NAV-2729 was from Tocris (Bristol, UK) and YM-254890 was from Wako Pure Chemical Industries (Richmond, VA) and U0126 was from Abcam (Cambridge, UK). The IP-One HTFR assay kit was from Cisbio (Codolet, France). Dulbecco’s Modified Eagle’s Medium GlutaMAX (DMEM), FluoroBrite DMEM, Roswell Park Memorial Institute (RPMI) 1640, and Dulbecco’s phosphate-buffered saline without calcium and magnesium (DPBS) were from Fisher Scientific (Hampton, NH). Penicillin /Streptomycin (10,000 U/mL), L-Glutamine and Lipofectamine 2000 were from ThermoFisher Scientific (Waltham, MA). Fetal bovine serum (FBS) was from Gemini Bio-Products (West Sacramento, CA). Poly-D-lysine, TPA, lithium chloride (LiCl), propidium Iodide (PI, 50 μg/ml), sodium orthovanadate, aprotinin and phenylmethylsulfonyl fluoride (PMSF) were from Sigma-Aldrich (St. Louis, MO). Dodecyl-D-maltopyranoside (DM) was from Anatrace (Berkshire, UK). HEK293T cells were from ATCC (Manassas, VA) and Melan-A cells were provided by D. Bennett (St. George’s Hospital, University of London). White low-volume 384-well plates and black CELLSTAR 96-well microplates (polystyrene wells flat bottom) were from Greiner (Monroe, NC). Bovine serum albumin (BSA) fraction V, fatty acid-free, and cOmplete protease inhibitor tablets was from Roche (Basel, Switzerland). NEBuilder Hifi DNA Assembler, Dpn1, T4 DNA Ligase, Q5 Hot Start High-Fidelity DNA Polymerase, and dNTPs were from New England BioLabs (Ipswich, MA). Quikchange Site-Directed Mutagenesis Kit was from Agilent Technologies (Santa Clara, CA) and TagMaster Site-Directed Mutagenesis Kit was from GM Biosciences Inc. (Frederick, MD). QIAGEN Plasmid Maxi Kits and QIAprep Spin Miniprep Kit were from QIAGEN (Germantown, MD). BRET substrate methoxy e-Coelenterazine was from NanoLight Technology (Pinetop, AZ). The synthetic vector for the G alpha q protein cloned into pcDNA3.1(+) was from the cDNA resource center (www.cdna.org) (#GNA0Q00000). The rabbit anti-phospho-p44/42 MAPK (Erk1/2) (Thr202/Tyr204) polyclonal antibody was from Cell Signaling technology ((#9101), Danvers, MA), the mouse total ERK2 (D-2) monoclonal antibody was from Santa Cruz ((#sc-1647), Dallas, TX), the secondary antibodies Goat anti-rabbit IRDye 800CW for channel 800nm and Goat anti-mouse IRDye 680RD for channel 700nm were from LI-COR (Lincoln, NE). NuPage protein gels and SeeBlue Plus2 pre-stain protein standards were from ThermoFisher Scientific, Odyssey Blocking buffer was from LI-COR, and Immobilon-P PVDF membranes were from Merck Millipore (Burlington, MA).

### IP1 Assays

#### Plasmids

The synthetic vector encodes human CysLTR2 cDNA in pcDNA3.1(+) fused to a N-terminal FLAG tag (DYKDDDDK) and a C-terminal 1D4 epitope tag (TETSQVAPA) ^6^. The FLAG tag was deleted by site-directed mutagenesis using TagMaster Site-Directed Mutagenesis Kit according to the manufacturer’s instructions. CysLTR2 3.43, 7.49, and 7.53 mutants and G alpha q protein mutants were generated by site-directed mutagenesis using QuikChange Lightning Site-Directed Mutagenesis Kit. The PCR reactions were performed according on the manufacturer’s instructions with some modifications. To generate CysLTR2 3.43 and G alpha q mutants, the specified reactions volumes were modified to half reactions using 25 ng template DNA. Similarly, to generate CysLTR2 7.49, and 7.53 mutants, volumes were modified to quarter reactions using 15 ng of template DNA. Quikchange primers and TagMaster primers were used to introduce the various mutant constructs and are listed in Suppl. Table S8. All constructs were confirmed by sequencing.

#### Cell Culture and Transfection

HEK293T cells from the American Type Culture Collection (ATCC) were maintained in DMEM GlutaMAX supplemented with 10% FBS (passage numbers 5 to 14). Melan-A cells were provided by D. Bennett (St. George’s Hospital, University of London) and were grown in RPMI with 10% FBS and 200 nM TPA unless specified otherwise. All cells were maintained at 37 °C under 5% CO_2_.

HEK293T cells were transiently transfected directly ‘in-plate’ in low-volume 384-well plates using lipofectamine 2000 according to manufacturer’s instructions with some modifications. The total DNA amount was kept constant at 11 ng per well using empty vector pcDNA3.1(+). Briefly, the appropriate amount of plasmid DNA was mixed with DMEM GlutaMAX (no FBS). In a separate mixture, the total Lipofectamine 2000 (2.5 µL per µg DNA) was mixed in DMEM GlutaMAX (no FBS) and incubated for 5 minutes. The appropriate amount of Lipofectamine 2000/ DMEM GlutaMAX mixture is mixed with the DNA/ DMEM GlutaMAX and incubated for 20 minutes. Cells were then trypsinized, re-suspended in DMEM GlutaMAX supplemented with 20% FBS and counted. Cells were mixed with the DNA/ Lipofectamine 2000/ DMEM mixture, and directly plated onto 0.01% poly-D-lysine coated, white, clear-bottom, tissue culture treated low volume 384-well plates at a density of 8 000 cells per well in 7 µL. All assays were conducted 24 hours after the transfection. All plasmids were prepared from QIAGEN Plasmid Maxi Kits, resulting in a high quality and high concentration plasmid solution (about 1000 - 4000 ng/µL) unless otherwise specified. For specific experiments, the plasmids were prepared from QIAprep Spin Miniprep Kit resulting in a 10-fold lower concentration.

#### IP1 Accumulation Assay

The CisBio IPone homogeneous time-resolved immunoassay quantifies D-myo-inositol-1-phosphate (IP1), a degradation product of the second messenger D-myo-inositol-1,4,5-trisphosphate (IP3), to measure activation of PLC-β by Gq-coupled GPCRs ^59^. The IP1 assay is a competitive homogenous time resolved fluorescence (HTRF) assay where the d2-labeld IP1 analog acts as the fluorescence acceptor and the terbium cryptate-labeled anti-IP1 mAb acts as the fluorescence donor. The terbium-cryptate is a long-lifetime fluorescence donor that can be excited by UV light. Lithium chloride is added during the stimulation period of the assay to block further degradation of IP1 by the enzyme inositolmonophosphatase (IMPase). It has been suggested that PLC-β-dependent IP1 accumulation also includes contributions from the IP2 formed by direct hydrolysis of PI4P, instead of IP2 formed by the dephosphorylation of IP3 formed by hydrolysis of PIP2 ^45^. Since this novel pathway does not depend on PIP2, which is predominantly found at the plasma membrane, the PI4P-dependent IP1 accumulation could have a substantial contribution to PLC-β signaling from the endosomal compartment.

All assays stimulated and incubated cells with agonists and inhibitors in the presence of LiCl. Following incubation, cells were lysed by addition of d2-labeled IP1 analog and terbium cryptate-labeled anti-IP1 mAb diluted in the kit-supplied lysis and detection buffer, and the plates were incubated overnight at RT. All time-resolved fluorescence signals were read using the BioTek Synergy NEO plate reader (BioTek Instruments, Winooski, VT) in the Rockefeller University’s High Throughput and Spectroscopy Resource Center (HTSRC). The plate is first subjected to flash lamp excitation at 320 nm, and then the fluorescence is measured at wavelengths centered at 620 nm and 655 nm simultaneously.

#### LTD4 Dose-Response of CysLTR2 DNA Titration

HEK293T cells were transiently transfected with a serial dilution of 11, 3.6, 1.2, 0.4, and 0.1 ng of wt and L129Q mutant of CysLTR2-1D4 and CysLTR2-GFP10-1D4 (BRET^2^ construct described later) per low volume 384-well as described above. 24 hours after transfection, the assay plate was placed on an aluminum heating block maintained at 37 °C, and cells were treated with 7 µL/ well of various concentrations of LTD4 (final concentrations from 10^-6^ M to 10^-11^ M) diluted in pre-warmed stimulation assay buffer provided by the manufacturer (10 mM HEPES, 1 mM CaCl_2_, 0.5 mM MgCl_2_, 4.2 mM KCl, 146 mM NaCl, 5.5 mM glucose, 50 mM LiCl, pH 7.4) supplemented with 0.2% BSA and 50 mM LiCl to prevent IP1 degradation. The plate was incubated at 37 °C for 2 hours. Following incubation, cells were lysed by addition of 3 µL/ well of d2-labeled IP1 analog and 3 µL/ well of terbium cryptate-labeled anti-IP1 mAb diluted in the kit-supplied lysis and detection buffer. The HTRF assay was read after overnight incubation in the dark.

#### Time-course

HEK293T cells were transiently transfected with 11 ng of CysLTR2-1D4 (wt and L129Q) per low volume 384-well. 24 hours after transfection, the assay plate was placed on an aluminum heating block as described above, and the cells were treated every 20 minutes over 180 minutes with 3.5 µL/ well of Lithium Chloride (LiCl) diluted in pre-warmed stimulation assay buffer at a final concentration of 50 mM. Cells were incubated at 37 °C. 1 hour after the start point, 3.5 µL/well of LTD4 diluted in pre-warmed DMEM GlutaMAX at a final concentration of 100 nM (agonist stimulated) or 3.5 µL/ well of DMEM GlutaMAX alone (basal) were added in appropriate wells and incubated for 2 hours. The reaction was then stopped by successively adding 3 µL/ well of the d2-labeled IP1 analogue and the terbium cryptate-labeled anti-IP1 mAb in reverse chronological order.

#### Dose-Response and Site-Saturation of CysLTR2 3.43, 7.49 and 7.53 Mutants

HEK293T cells were transiently transfected with 11 ng of wt and L129^3.43^ mutants of CysLTR2-1D4 or the 8 combinations of CysLTR2-3.43/7.49/7.53 mutants per low volume 384-well as previously described. 24 hours after transfection, cells were treated with 7 µL/ well of various concentrations of LTD4 diluted in the supplemented assay buffer (final concentrations from 1 µM to 10 pM) for the dose-response assay. For the site-saturation experiments, cells were treated with 7 µL/ well of buffer alone or LTD4 at a final concentration of 100 nM or buffer alone. Following 2 hours incubation at 37 °C, cells were lysed as described above. The plates were read and IP1 concentrations were determined, as before. All the plasmids for the CysLTR2-L129^3.43^ mutants and CysLTR2-N301^7.49^ and/or CysLTR2-Y305^7.53^ mutants except wt and L129Q^7.49A^ (namely LAY and QAY) were prepared from QIAprep Spin Miniprep Kit. Miniprep elutions have DNA concentrations in the range 100-500 ng/µL.

#### Inverse Agonist Competition Assay

HEK293T cells were transiently transfected with 11 ng of FLAG-CysLTR2-1D4 wt, L129Q, L129E and L129A per low volume 384-well as previously described. CysLTR2-L129A and - L129E were prepared from QIAprep Spin Miniprep Kit. 24 hours after transfection, cells were treated with 3.5 µL/ well of various concentrations of HAMI-3379 or BayCysLT2 (final concentrations from 30 µM to 10 pM) or equivalent amounts of DMSO diluted in the supplemented assay buffer. Following 1 hour incubation, 3.5 µL/ well of the assay buffer with or without 10^-7^ M LTD4 were added for 1 hour and 45 minutes and cells were lysed as described above.

#### Gene Dosage of CysLTR2 with GNAQ Mutants and Inhibitors

HEK293T cells were transiently transfected with a serial dilution of 11, 3.6, 1.2, 0.4, and 0.1 ng of CysLTR2-1D4 wt or L129Q and 11 ng of G alpha q proteins mutants GNAQ-Q209L, Q209P and R183Q per low volume 384-well as described above. 24 hours after transfection, cells were treated with 3.5 µL/ well of 30 µM NAV-2729 or 1 µM YM-254890 diluted in pre-warmed DMEM GlutaMAX. Following 1 hour incubation, 3.5 µL/ well of pre-warmed supplemented stimulation buffer supplemented were added for 2 hours prior to cells lysis. For all experiments involving YM-254890 and NAV-2729, the amount of DMSO was kept constant.

#### Dose-Response of YM-254890 and NAV-2729

HEK293T cells were transiently transfected with 11 ng of CysLTR2-1D4 wt or L129Q constructs per low volume 384-well. After 24 hours, cells were treated with 3.5 µL/ well of various concentrations of YM-254890 or NAV-2729 (final concentrations from 30 µM to 1 pM) diluted in DMEM GlutaMAX and incubated for 24 hours (48 hours after transfection) or 3 hours (27 hours after transfection) at 37 °C. 2 hours prior to cell lysis, 3.5 µL/ well of supplemented assay buffer were added to each well. Cells were lysed as described above.

#### Dose-Response of YM-254890 in HEK293T cells and Melan-a Cells

HEK293T cells were transfected as previously described. Melan-a cells expressing CysLTR2-L129Q or empty vector MSCV-PURO were seeded in low volume 384-well plates at a density of about 5 000 cells/well. After 24 hours, 3.5 µL/ well of various concentrations of YM-254890 (final concentrations from 10^-4^ M to 10^-12^ M) diluted in the supplemented assay buffer were added to each well. Following 1 hour incubation, 3.5 µL of assay buffer was added for 2 hours prior to cell lysis. Following 1 hour incubation, 3.5 µL/ well of supplemented stimulation assay buffer were added for 2 hours prior cells lysis. Cells were lysed as described above. The plates were read and IP1 concentrations were determined, as before.

#### Data Reduction

The raw signals were transformed into a fluorescence ratio, 665 nm/620 nm, and IP1 concentrations were interpolated from a standard curve prepared using the supplied IP1 calibrator. The IP1 standard curve was fit to a sigmoidal curve using the equation y = Bottom + (Top - Bottom)/ (1 + 10^(Log IC50 - x). The fluorescence ratio was then converted into the corresponding IP1 concentration (nM) using the equation IC50x(y-top)/(Bottom-y) and the standard curve. In some cases, these concentrations were further analyzed to get a normalized IP1 value. First, the mock concentrations of IP1 (obtained from the average of the basal levels of accumulation for pcDNA-transfected cells) are subtracted from the raw IP1 concentrations to give the corrected IP1 concentrations. These data, in nM, are then converted into picomoles/ well, from a working volume of 20 µL per well. Each data set is then normalized relative to the unstimulated mock-transfected cells (set to 0%) and to the fully stimulated wild-type receptor (set to 100%).

IP1 concentrations or normalized IP1 data were fit to specific operational models introduced in Supplementary Scheme S1. For the LTD4 dose-response of CysLTR2 DNA Titration, the IP1 concentrations, in nM, were plotted against concentration of LTD4. CysLTR2 wt data are then fit to a sigmoidal dose-response function using the equation y =Basal + (EffectMax - Basal)/(1 + operate) where operate = (((10^log K_A_) + (10^x))/(10^(log τ + x)))^n. In this fit, the Basal (basal offset at zero agonist concentration), EffectMax (the maximal IP1 concentration at infinite agonist concentration), log K_A_ (affinity of the agonist for the receptor), and n were shared between all gene dosage curves. Individual gene dosage curves have unique log τ parameters, which represent total receptor concentration. As CysLTR2-L129Q did not follow this dose dependency, these data were fit to a horizontal line, y = Mean + 0(x). The Prism 8 software used to analyze these data did not allow for a fitting without parameter x, so we multiplied this by zero to negate its influence. Effectively, this fit is a horizontal line plotting the mean IP1 concentrations for all LTD4 doses.

For the time-course, the corrected IP1 concentrations, in nM, were plotted against time, in minutes. These data are then fit to a one-phase decay model using the equation y = (y_0_ - Plateau)*exp(-K*x) + Plateau. Y_0_ is the IP1 concentration at time zero while Plateau is the IP1 concentration at infinite time. K is the rate constant of the decay.

For the dose-response and site-saturation of CysLTR2 3.43 mutants, the normalized IP1 accumulation, in percentages, were plotted against concentrations of LTD4 and fit to a sigmoidal dose-response model using the equation y = basal + (Emax – basal)/ (1 + 10^(-pEC50 – x)). From these fits, the basal, agonist dependent activity (Emax), and pEC50 were compiled and plotted in descending order of total activity defined as the sum of basal and agonist-dependent activity. Furthermore, the two-state allosteric model was used to fit this data using the equation y = basal + (Effectmax Basal)*(10^(log τ + log K_q_)*(1+10^(log α + log β + x + log K_A_)))/(1 + (10^log K_q_)*(1 + 10^log τ) + (10^(x +log K_A_))*(1 + (10^(log α + log K_q_))*(1+10^(log β + log τ)))). The parameters basal, Emax, log α, log β, and K_A_ are fixed for all data. This model assumes that mutations don’t change the affinity of the agonist for the inactive state (K_A_ is fixed) and for the active state αK_A_ (α is fixed). The two fitting parameters, τ and K_q_, are unique for each mutant and describe the total receptor concentration, and the equilibrium constant for the agonist-independent equilibrium of the receptor with inactive and active states, respectively.

For the dose-response for the 8 combinations of CysLTR2 3.43/ 7.49/ 7.53 mutants, the IP1 concentrations, in nM, were plotted against LTD4 concentrations. These data are then fit to a sigmoidal fitting function and a horizontal line function, as described previously. The fits to these two functions are compared and the Akaike Information Criterion (AIC) was used to select the better model (Suppl. Table S1). From these fits, the basal, agonist dependent activity (Emax), and pEC50 were compiled and plotted in descending order of total activity defined as the sum of basal and agonist-dependent activity, as before.

For the inverse agonist competition assay, the normalized IP1 accumulation, in percentages, were plotted against concentrations of antagonists BayCysLT_2_ and HAMI3379, and fit to a sigmoidal dose-response model using the equation y = Bottom + (Top - Bottom)/ (1 + 10^(Log EC50 - x) as described above. In this fit, the Bottom or basal IP1 accumulation is shared for both the LTD4 stimulated and unstimulated curves, such that the maximal effect of the antagonists at infinite concentration converge for both curves. The pcDNA is fit to a horizontal line as described above.

For the gene dosage of CysLTR2 with GNAQ mutants and inhibitors, the corrected IP1 concentrations, in nM, were plotted against the logarithm of the DNA concentration. This gene dosage-dependent IP1 accumulation was fit to the semi-logarithmic function y = mlog(x) + c. In order to compare the overall signaling levels, the results were interpolated to a dosage resulting in 250 nM IP1 accumulation for the untreated condition. In the same way, the gene dosage was interpolated for IP1 accumulation of 250 nM for the samples treated with YM-254890 and NAV- 2729. These interpolated IP1 accumulation concentrations were then plotted and compared to each other using Tukey’s multiple comparisons test, to determine which means amongst a set of means differ from the rest. Those that are determined to be significantly different are given a p-value, shown above the bar graphs.

Lastly, for the dose-response of YM-254890 and NAV-2729, the IP1 concentrations, in nM, were plotted against concentrations of inhibitors, YM-254890 and NAV-2729. These data were fit to a sigmoidal dose-response model using the equation y = Bottom + (Top - Bottom)/ (1 + 10^(Log EC50 - x) as described above. All data analysis was conducted on Prism 8 software.

### BRET^2^ Assays

#### Designing Plasmids for PCR of Fragments

The NEBuilder HiFi DNA Assembly Tool was used to assemble the BRET^2^ acceptor constructs CysLTR2-GFP10-1D4. These were assembled from three parts: pcDNA 3.1(+) backbone from construct HA-CLIP-CLR ^60^, CysLTR2 and full-length C-terminal 1D4 epitope tag from FLAG-CysLTR2-1D4 mentioned above, and GFP10 from YB124_CXCR4-GFP10 ^24^. The primers were designed using the NEBuilder Assembly Tool on the NEB website. These primers were designed with a specific sequence to prime to the gene of interest for template priming (3’ end), as well as an overlap sequence to aid in assembly (5’ end). These oligonucleotides were purchased from IDT (Coralville, IA), purified at the standard desalting grade, and are shown in Suppl. Table S9.

#### PCR

The fragments introduced above were PCR amplified using Q5 Hot Start High-Fidelity DNA Polymerase and fresh dNTPs purchased from NEB. Briefly, the PCR reactions were performed in 25 µL total volume containing: 1x Q5 Reaction Buffer, 0.2 mM dNTPs, 0.5 µM of the forward primer, 0.5 µM of the reverse primer, 1 ng template DNA, and 1 unit of Q5 Hot Start High-Fidelity DNA Polymerase. The PCR thermocycle was as follows: initial denaturation at 98 °C for 30 secs, followed by 25 cycles of denaturation (98 °C, 10s), annealing (varied from 50-70 °C, 30 s), and elongation (72 °C, 3 min), and ending by a final elongation (72 °C, 2 min). The recommended annealing temperature calculated on the NEBuilder Assembly Tool was used for each primer pair. Following PCR, 1 unit of DpnI was added and the mixture was incubated at 37 °C for 30 minutes to digest any remaining template DNA. This was cleaned up, and any enzymes were removed using a DNA Clean and Concentrator (Zymo Research). The concentrations of all PCR-amplified fragments were determined using a NanoDrop.

#### Assembly

The NEBuilder Hifi DNA Assembler includes three enzymes; the exonuclease to create 3’ overhangs to aid annealing of neighboring fragments sharing a complimentary overlap region, the polymerase to fill the gaps of each annealed fragment, and the DNA ligase to seal nicks in the assembled DNA. The assembly reaction was performed in 20 µL total volume, with 50 ng of vector, 100 ng of insert(s), and 10 µL of the NEBuilder HiFi DNA Assembly Master Mix. This was incubated at 50 °C for 60 minutes, and 2 µL of the assembled product was used to transform NEB 5-alpha Competent *E. coli* cells. Cells were spread on LB-Amp plates and colonies were picked and confirmed by sequencing.

After the assembled product was confirmed by sequencing, the second NotI site that is flanked by two XhoI sites, which had been part of the pcDNA 3.1(+) backbone in HA-CLR-CLIP, was removed. This was simply done by digesting at the XhoI sites and self-ligating the vector using T4 DNA Ligase. We then sequenced the NotI-removed CysLTR2-GFP10-1D4 in its entirety to check for any erroneous modifications or linkages.

β-Arrestin1-RLuc3 and β-Arrestin2-RLuc3, the BRET^2^ donors paired to these acceptor constructs, were constructed previously by C-terminally fusing the coding sequence of β-Arrestin to RLuc3^24^.

#### Cell Culture and Transfection

HEK293T cells were maintained as described above, and transiently co-transfected with β-Arrestin1-RLuc3 or β-Arrestin2-RLuc3 and CysLTR2-GFP10-1D4 wt or L129Q directly ‘in-plate’, in 96-well plates, using lipofectamine 2000 as described above with slight modifications to account for the larger well volume. The total DNA amount was kept constant at 205 ng per well using empty vector pcDNA3.1(+). Briefly, a master-mix of the β-arrestin-RLuc3 was made in FluoroBrite DMEM (DMEM without phenol red and suitable for fluorescence experiments) and the CysLTR2-GFP10-1D4 DNA were added to these after appropriate distribution. In a separate mixture, the total Lipofectamine 2000 was mixed in FluoroBrite DMEM and incubated for 5 minutes. The appropriate amount of Lipofectamine 2000/ FluoroBrite DMEM mixture is mixed with the DNA/ FluoroBrite DMEM and incubated for 20 minutes. Cells were then trypsinized, re-suspended in FluoroBrite DMEM, 20% FBS, 30 mM HEPES, and 8 mM glutamine, and counted. Cells were mixed with the DNA/ Lipofectamine 2000/ FluoroBrite DMEM mixture, and directly plated onto 0.01% poly-D-lysine coated, black, clear-bottom, tissue culture treated 96-well plates at a density of 40 000 cells per well in 100 µL FluoroBrite DMEM 10% FBS, 15 mM HEPES, 4 mM glutamine. All assays were conducted 24 hours after the transfection. All plasmids were prepared from QIAGEN Plasmid Maxi Kits, resulting in a high quality and high concentration plasmid solution (about 1000 - 4000 ng/µL).

#### DNA Titration Assay

HEK293T cells were transiently transfected with 5 ng of β-Arrestin1 or 2-RLuc3 and 0, 12.8, 32, 80, and 200 ng of CysLTR2-GFP10 wt or L129Q per 96-well. 24 hours after transfection, media were aspirated carefully from all wells. 40 µL of pre-warmed BRET buffer (DMEM FluoroBrite, 15 mM HEPES, 0.1% BSA, 4 mM Glutamine) was added to each.

All BRET^2^ measurements were taken on the BioTek Synergy Neo2 microplate reader (HTSRC) using filter set 109 (center wavelength/band width) of 410/80 nm (donor) and 515/30 nm (acceptor). First, the GFP fluorescence was read using the monochromator (ex: 395 nm, em: 510 nm +/- 20 nm from bottom, auto gain) to quantify total expression levels. Following this, the cell-permeable substrate methoxy e-Coelenterazine (Me-O-e-CTZ/ Prolume Purple) was added to each well at a final concentration of 5 µM and the luminescence at the two wavelengths were read simultaneously.

#### Time Course Assay

HEK293T cells were transiently transfected with 5 ng of β-Arrestin1 or 2-RLuc3 and 80 ng of CysLTR2-GFP10 wt or L129Q per 96-well. 24 hours after transfection, media were aspirated carefully from all wells. 30 µL of pre-warmed BRET buffer was added to each well, and the GFP fluorescence was read. Methoxy e-Coelenterazine was added to 2 columns at 5 µM final concentration, followed by addition of 0 nM, 30 nM, and 1000 nM of LTD4 to appropriate wells in the two columns. The plate was quickly placed into the microplate reader so that there is as little lag time between addition of the ligand and BRET readings as possible. The two columns take about 45 seconds to read, and this was repeated 30 times such that we have a BRET reading every 45 seconds for about 23 minutes.

#### Agonist Dose-Response Assay

HEK293T cells were transiently transfected with 5 ng of β-Arrestin1 or 2-RLuc3 and 80 ng of CysLTR2-GFP10 wt or L129Q per 96-well. 24 hours after transfection, media were aspirated and 30 µL of pre-warmed BRET buffer was added to each well. Various concentrations of LTD4 (final concentrations from 10^-6^ M to 10^-11^ M) were added to appropriate wells and incubated for 10 minutes at room temperature. Following the incubation, GFP and BRET^2^ signals were obtained as described above.

#### Data Reduction

Raw BRET^2^ ratios were determined by calculating the ratio of the light intensity emitted by the GFP10 (515 nm) over the light intensity emitted by the RLuc3 (410 nm). The BRET^2^ signals are normalized to basal BRET^2^ signals (β-Arrestin-RLuc3 only signals) to give corrected BRET^2^ ratios.

For the DNA titration assays, the BRET^2^ ratios were plotted against GFP10 fluorescence readings to create a DNA titration curve ^61^. These data are fit to a one-site saturation-binding function using the equation y = B_max_ (x / K_d_ + x) + NS(x) + background, where NS (non-specific binding) is set to zero, and both B_max_ (the maximal recruitment at infinite agonist concentration) and background (basal offset at zero agonist concentration) are shared for all three curves (wt, L129Q, and wt + LTD4). The K*_d_* is varied for the three curves, and this gives GFP10 signal of the receptor giving half-maximal BRET^2^ ratios.

For the time course assays, the BRET^2^ ratios were plotted against time, in minutes, to assess the time-dependence of the LTD4 stimulated β-Arrestin recruitment. These data are fit to a two-phase decay model using the equation y = y_0_ + If (x > x_0_, Plateau*(-exp(-(10^log K_Fast_)*(x - x_0_)) + exp(-(10^log K_Slow_)*(x - x_0_)))*(10^log K_Fast_)/((10^log K_Fast_)-(10^log K_Slow_)), 0). K_fast_ describes the initial recruitment of β-Arrestin dependent on the concentration of active receptor, and is represented as the initial increase in signal. K_slow_ describes the disassembly of the receptor-β-Arrestin complex and is represented by the decay of the signal over time. For this fit, x_0_, y_0_ (the BRET2 signal at time zero), and log K_slow_ are shared for all three curves (0 nM, 30 nM, and 1000 nM LTD4). Log K_fast_ is varied for the three curves, and gives the rate constant of β-Arrestin recruitment for the three conditions.

Lastly, for the agonist dose-response assays, the BRET^2^ ratios were plotted against concentrations of LTD4. The cysLTR2-GFP10 wt data were fit to a dose response sigmoidal curve using the equation y = Bottom + (Top - Bottom)/ (1+10^(Log EC50 - x). As CysLTR2-L129Q-GFP10 did not follow this dose dependency, these data were fit to a horizontal line, y = Mean + 0(x), as described previously. These data were then normalized using the top and bottom parameters for the sigmoidal fits of the wild-type data. This time, when fitting the CysLTR2-GFP10 wt using the same sigmoidal curve mentioned previously, the top was constrained to equal 100, and the bottom to equal zero. The fitting of CysLTR2-L129Q-GFP10 is the same but with normalized values.

### Flow Cytometry

The HEK-293T cells were transiently transfected using lipofectamine 2000 with 2 pg/cell total DNA and 0, 0.02, 0.2, or 2 pg/cell receptor-encoding DNA using previously described in-plate transfection protocol. The total DNA amount was kept constant at 2 µg per well using empty vector pcDNA3.1(+), and 600 000 cells were seeded in 2 mL per 6-well in DMEM GlutaMAX, 10% FBS.

Cells were resuspended in the well media then centrifuged at 300 × g for 5 min. The cell pellets were gently washed three times in 1X Dulbecco’s PBS without calcium and magnesium supplemented with 0.1% BSA (300 × g, 5 min), then suspended in 250 µl PBS (to a concentration of ∼2.0 × 10^6^ cells/ml). Cells were mixed 1:1 with 110 ng/ml (2X) PI, the optimal PI concentration for HEK293T cells previously determined by titration experiments (data not shown). Samples were filtered through a cell strainer cap (Falcon, Ref 352235) then place on ice for immediate analysis of GFP10 expression and live-dead discrimination (PI).

Analyses were performed by flow cytometry using Cytek Aurora spectral flow cytometer equipped with four lasers (405, 488, 561 and 640 nm) and SpectroFlo software (version 2.0.1). Forward and side scatter parameters were used to eliminate debris from analysis. For each sample 20,000 events in the SSC singlet gate were collected. Spectral unmixing was performed using unstained pcDNA transfected cells, single stained PI pcDNA transfected cells, and CysLTR2-WT-GFP10 cells as the unstained, PI positive, and GFP10 positive controls, respectively. Unmixed data was prepared using FlowJo (version 10.5.3).

### Live cell imaging

The HEK293T cells were transiently transfected using lipofectamine 2000 with 2 pg /cell DNA using the previously described in-plate transfection protocol. In 35 mm glass bottom microwell dishes (MaTek; Cat#P35G-1.5-14-C), coated with 0.2% gelatin Type A (Sigma; EC No 232-554-6) fixed with 0.5% glutaraldehyde (Electron Microscopy Sciences Cat 16316), 600 000 cells in 2 ml DMEM 10% FBS were seeded. Cells were incubated at 37°C and 5% CO_2_ for ∼22h. Cell media was aspirated from each well and replaced with 2 ml FluoroBrite media supplemented as described above. Z-stacks of cells were acquired using a DeltaVision Image Restoration Microscopy System (Applied Precision) and Resolve3D softWoRx-Acquire (version 6.5.2 Release RC1). Samples were kept in an enclosed environmental chamber kept at 37°C during imaging. An Olympus 60X/1.3 objective was used. Excitation at 390nm with 18nm bandpass. Emission at 525nm with 50nm bandpass. CoolSNAP HQ/ICX285 CCD camera was used. Deconvolution of z-stacks were performed in softWoRx.

### ERK Phosphorylation Assay

HEK293T cells were transiently transfected with 1 µg of CysLTR2 wt or CysLTR2-L129Q using lipofectamine 2000 as previously described. The total amount of DNA was kept constant at 2 µg/well using pcDNA3.1(+). Briefly, 1 million cells/ well were seeded in 2 mL DMEM Glutamax, 10% FBS in PDL-coated 6-well plates. 20 hours post transfection, the growth media was replaced by 1.5 mL of pre-warmed DMEM GlutaMAX (no FBS) to subject cells to 3 hours of FBS starvation. 500 µL of the following compounds diluted in pre-warmed DMEM GlutaMAX were added to the appropriate wells: 1 µM YM-254890, 30 µM NAV-2729, 10 µM U0126, 10 µM U73122, 200 nM TPA or corresponding volumes of DMSO for control basal wells. Cells were incubated for 3 hours at 37°C. Cells expressing CysLTR2 wt were then stimulated with 200 µL of 100 nM LTD4 for 3 minutes. The reaction was stopped by placing the plates on ice, and cells were harvested and washed once with cold PBS. Cells were incubated with 200 µL solubilization buffer (50 mM Tris-HCl, 100 mM NaCl, 1 mM MgCl_2_) supplemented with 1% (w/v) DM, 1 µg/µl aprotinin, 1mM PMSF, cOmplete protease inhibitor, and 1 mM sodium orthovanadate for 1 hour at 4°C. Following incubation, cells were centrifuged at 20 000 x g for 20 min at 4°C. Total protein was determined by Protein DC assay according to manufacturer’s instructions (Bio Rad).

#### Immunoblotting

Briefly, 30 µg of lysate were mixed with LDS-NuPage loading buffer supplemented with 100 mM dithiothreitol (DTT). Denatured lysates were resolved by SDS-PAGE on NuPAGE 4-12% Bis-Tris gel (1.5mm) at 120V for 1h45 and transferred electrophoretically onto an Immobilon-P PVDF membrane using a semi-dry transfer (45min at 18V). Membranes were blocked for 1 hour in Odyssey Blocking Buffer at room temperature before being incubated overnight at 4°C with primary antibodies: rabbit anti-phospho-ERK1/2 polyclonal pAb and mouse anti-total ERK2 mAb diluted at 1:1 000 in the Odyssey blocking buffer. Membranes were washed three times for 10 minutes in PBS-T (PBS supplemented with 0.1% Tween). Secondary antibodies, goat anti-rabbit IRDye 800CW and goat anti-mouse IRDye 680RD, diluted at 1:20 000 in Odyssey Blocking Buffer supplemented with 0.1% Tween and 0.01% SDS, were added and the membranes were incubated for1 hour at room temperature, protected from light. Following three washes in PBS-T for 10 minutes, the membranes were scanned with a LI-COR Odyssey SA imager to visualize the phosphorylated ERK1/2 signal (800 nm channel detection) and the total ERK2 signal (700 nm channel detection) at the HTSRC.

### Molecular Modeling

We established our protocols initially using receptors with crystal structures in active and inactive conformations (rhodopsin and µ opioid receptor) to test the *in silico* mutagenesis methods. In these initial tests, we also tried the *mp_relax* and *mp_mutate_relax* programs of Rosetta ^62^, but found that the membrane mimetic introduced some artifacts when comparing multiple conformations. The CysLTR2 homology models in the Active and Inactive states were taken from the version of the GPCRdb dated 2018-07-10. The models were optimized with the *relax* program from the Rosetta3 software suite ^63^ release 2018.33 using the *beta_nov16_cart* all-atom sore function ^64^ with constraints to maintain the overall conformation close to the starting structure. The mutations were introduced by the *cartesian_ddg* program of Rosetta and the local neighborhood within 0.9 nm of the mutated residue was repacked and optimized using backbone constraints. Five structures (‘iterations’) with two hundred cycles of optimization were calculated for each mutant to obtain a score (‘free energy change’) of the mutation for each iteration. The minimum score is reported as “ddG Activation” or “ddG Inactivation” depending on the starting structure used. The difference of these two scores “Activation – Inactivation” is a prediction of the free energy change ΔΔG(active)-ΔΔG(inactive) due to the mutation, which predicts the change of the active – inactive state equilibrium (R_i_–R_a_). Structures were visualized with vmd1.9.3 ^65^.

## Supplementary Figures and Tables

**Supplementary Figure S1.**
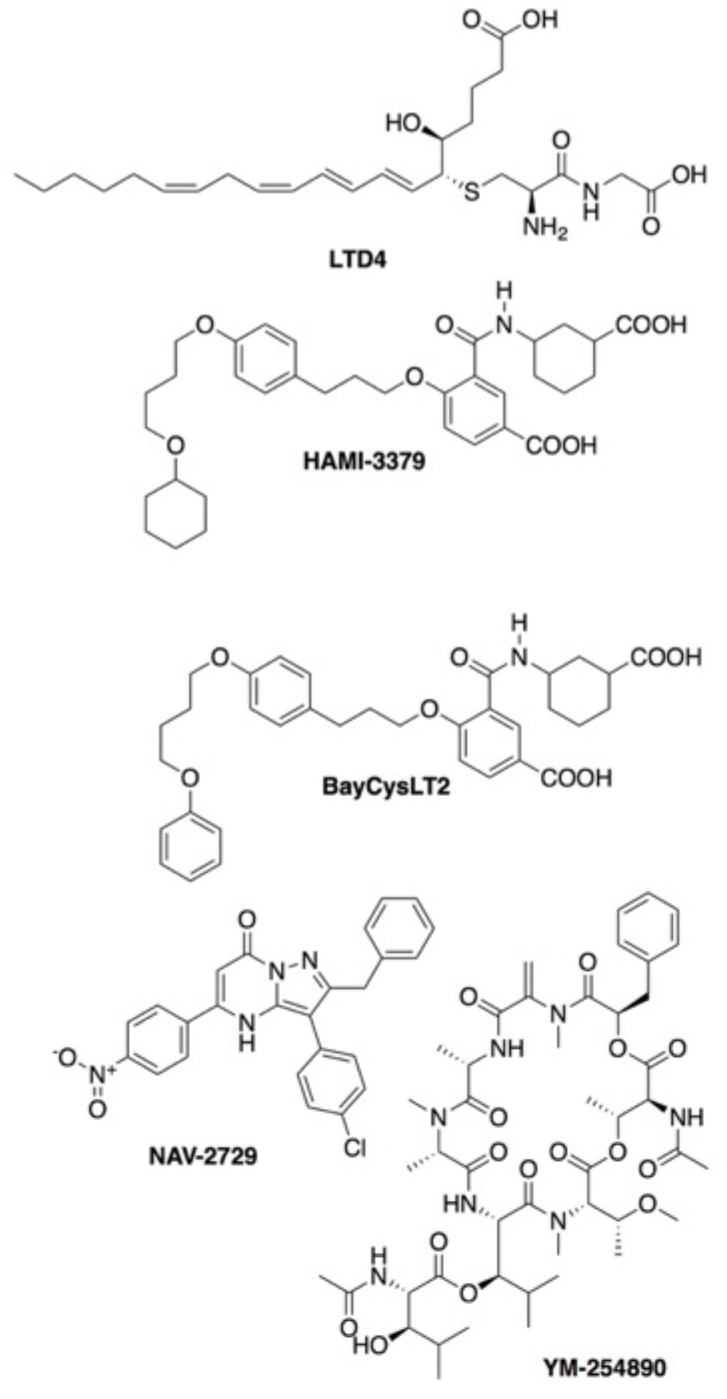
Molecular structure of the diverse compounds tested for IP1 accumulation inhibition studies. Shown are the CysLTR2 agonist LTD4 and antagonists HAMI-3379 ^66^ and BayCysLT2, the Arf6 inhibitor NAV-2729 ^42^ and the Gq/11 inhibitor YM-254890 ^41^.

**Suppl. Fig. S2.**
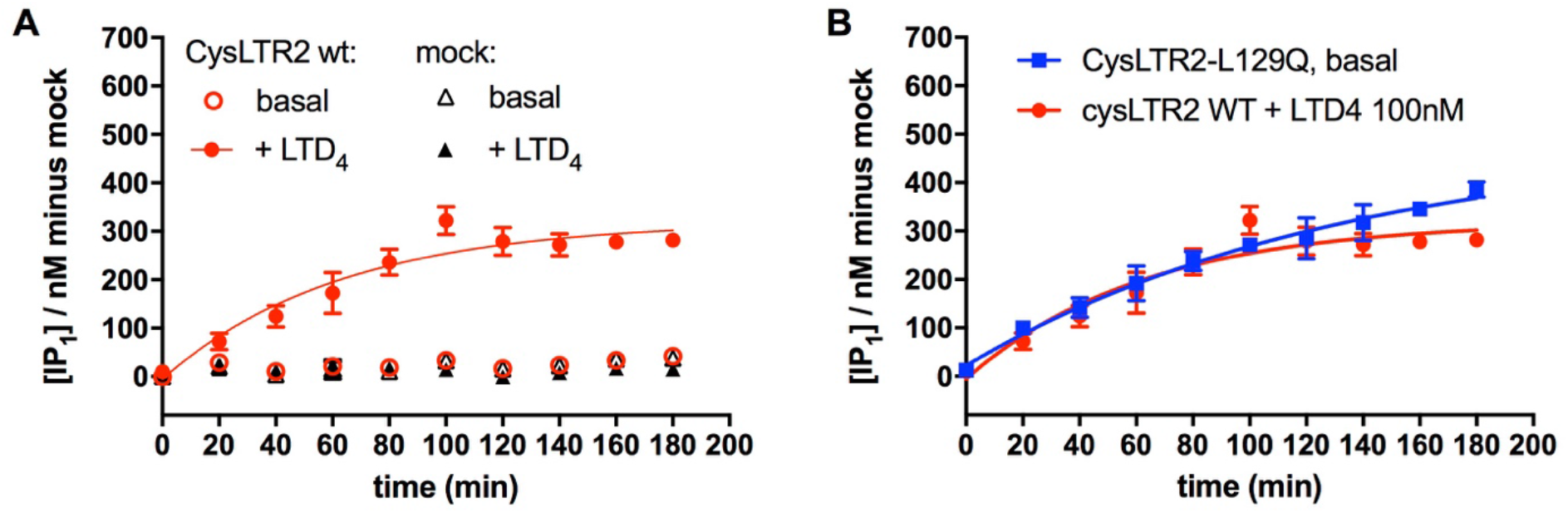
Lithium-dependent accumulation of IP1 differs for agonist-induced and constitutive receptor activity. Lithium chloride (LiCl) is added during the stimulation period of the assay to block further degradation of IP1. (**A-B**) Time-course of the effect of 50 mM LiCl on the basal and LTD4-induced IP1 accumulation for CysLTR2 wt (**A**) and L129Q (**B**) transfected HEK293T cells over 180 minutes. (**A**) Basal IP1 accumulation of CysLTR2 wt (open red circles) is comparable to mock with (solid black triangles) or without (open black triangles) LTD4 stimulation and is not affected by the addition of LiCl. CysLTR2 wt stimulated by LTD4 exhibits an increasing IP1 accumulation over 100 minutes after addition of LiCl, before reaching a plateau (solid red circles). (**B**) CysLTR2-L129Q (blue squares) shows a LTD4-independent IP1 accumulation similar to the wt receptor that continues to increase over 180 minutes. The data are expressed as the mean ± SEM of IP1 (nM) minus mock and result from one experiment performed in four technical replicates.

**Suppl. Fig. S3.**
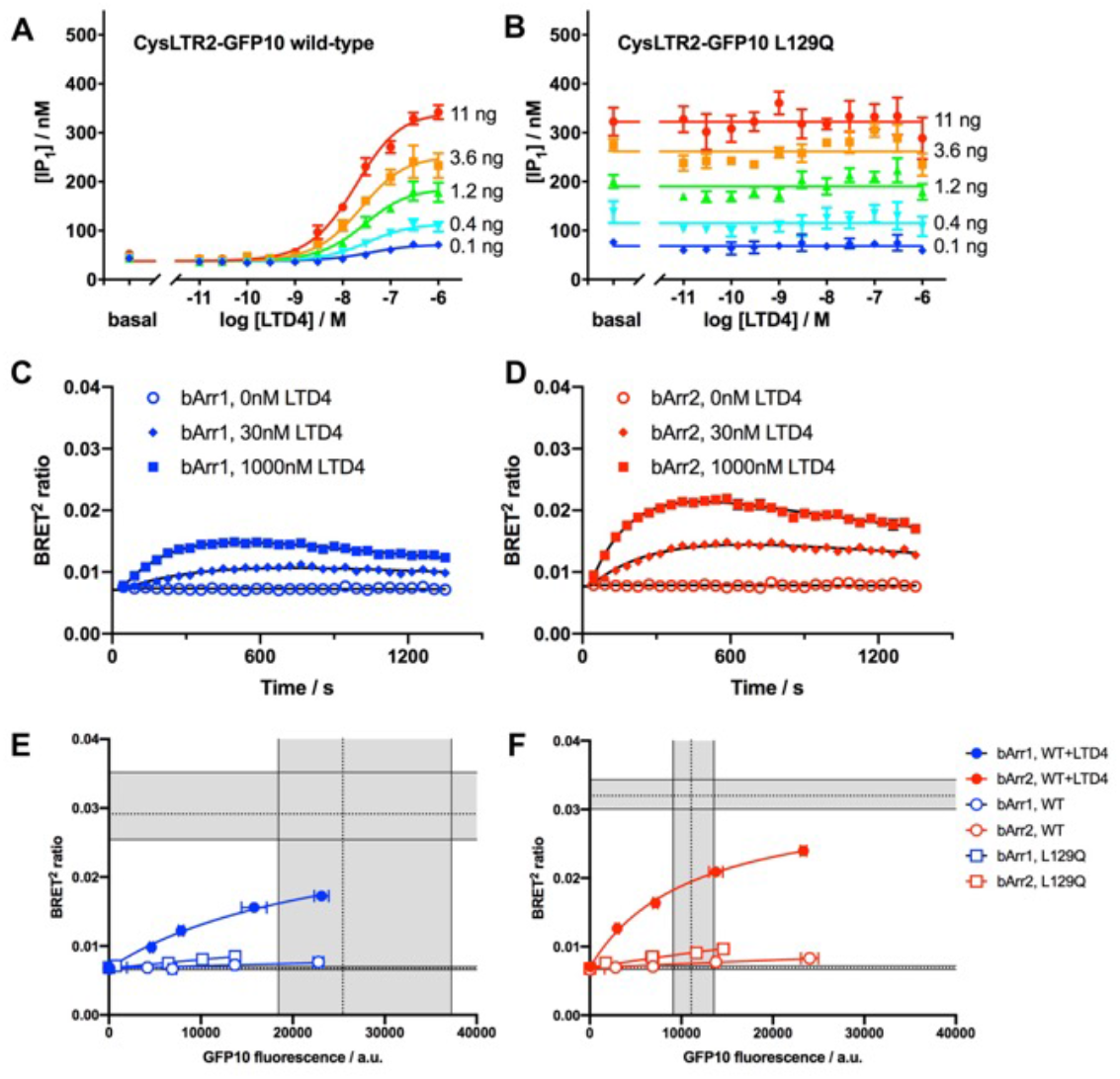
Development of the CysLTR2 BRET^2^ assay. (**A**) The agonist LTD4 leads to dose-dependent inositol monophosphate (IP1) accumulation in HEK293T cells expressing different levels of CysLTR2-GFP10 wild-type as controlled by different amounts of receptor-encoding plasmid DNA transfected at constant total DNA (red, 11 ng; orange, 3.6 ng; green, 1.2 ng; cyan, 0.4 ng; blue, 0.1 ng). The curves are fits of the dose-response data to an operational model. (**B**) The corresponding experiment with the mutant CysLTR2-GFP10-L129Q shows no significant dose-dependent response to LTD4, whereas the ligand-independent basal IP1 accumulation increases with increasing amount of CysLTR2-GFP10-L129Q-encoding plasmid DNA. The results show that CysLTR2-GFP10-L129Q is a CAM with a basal activity corresponding to about 100% of the WT receptor maximally stimulated with agonist. Data are expressed as mean ± SEM of concentration of IP1 accumulated (nM) and result from one experiment carried out in four technical replicates. Each plot represents one individual assay plate. (**C,D**) Time-course of LTD4-stimulated β-Arrestin-recruitment measured by the BRET^2^ assay with CysLTR2-GFP10 and β-arrestin-RLuc3. (**C**) The time-dependent increase of net BRET^2^ demonstrates recruitment of β-arrestin 1, and was measured for two LTD4 concentrations (30 nM, blue diamond; 1000 nM, blue square) and vehicle (0 nM, blue open circle). (**D**) Time-course of β-arrestin 2 recruitment for two LTD4 concentrations (30 nM, red diamond; 1000 nM, red square) and vehicle (0 nM, red open circle). The time-dependent data in (**C,D**) were globally fitted with a double exponential function using shared slow kinetic rates and starting values. They are the mean ± SEM from two independent experiments, with two sets of four technical replicates. (**E,F**) Saturation-binding BRET^2^ experiment with CysLTR2-GFP10 wild-type stimulated with 1000 nM LTD4 (circle), unstimulated CysLTR2-GFP10-L129Q (open square), and unstimulated CysLTR2-GFP10 wild-type (open circle). The data are fit to a one-site saturation binding function and are the mean ± SEM from five technical replicates on one plate. (**E**) Recruitment of β-arrestin-1-Rluc3 (blue). (**F**) Recruitment of β-arrestin-2-Rluc3 (red).

**Suppl. Fig. S4.**
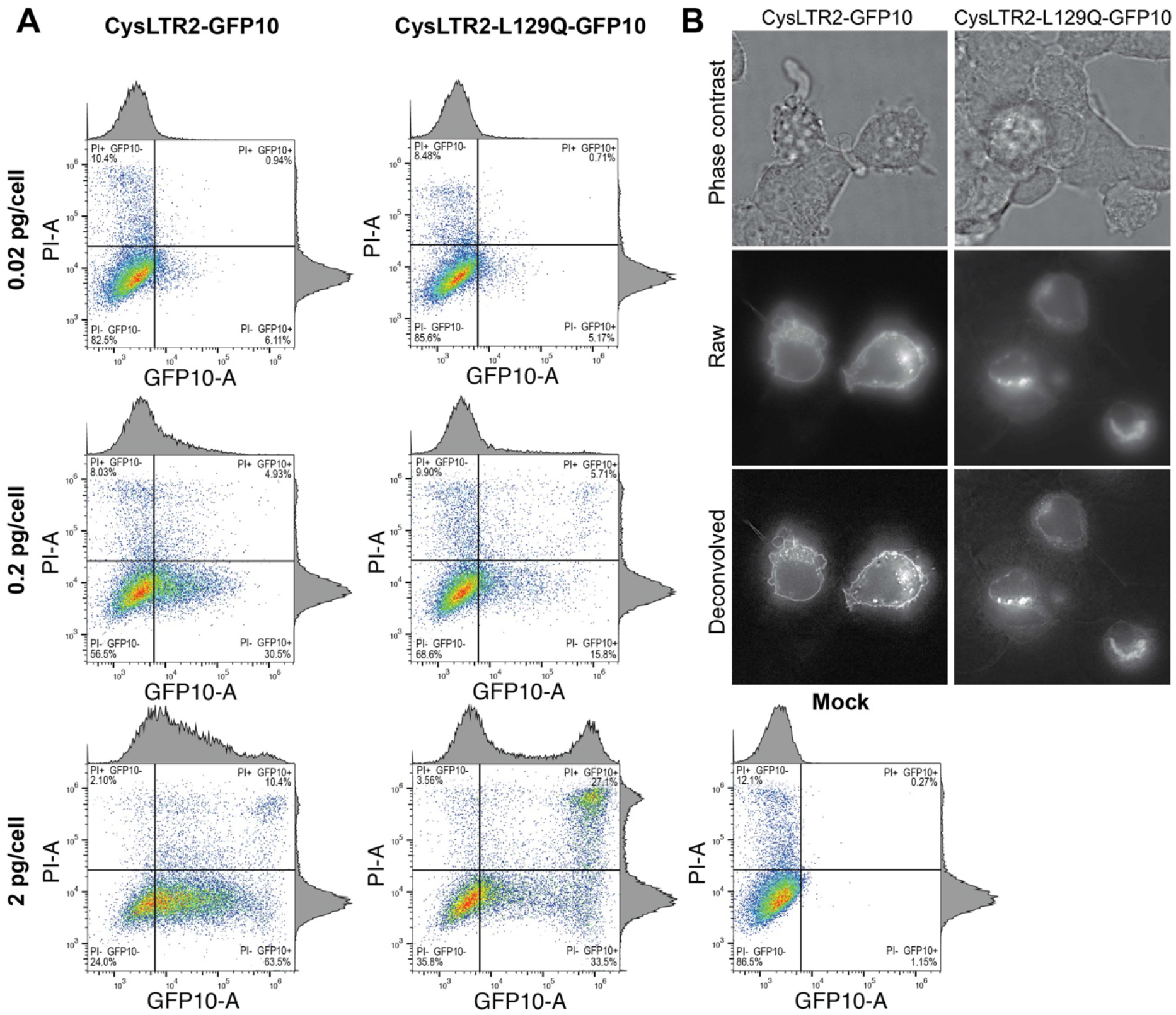
High levels of CysLTR2-L129Q result in cell death. (**A**) The toxicity of CysLTR2-L129Q-GFP10 and CysLTR2-WT-GFP10 was assessed using flow cytometry for GFP10 and the live-dead cell marker propidium iodide (PI). Data shown has been unmixed to correct for spectral overlap between GFP10 and PI. Representative dot plots with gating for HEK293T cells transfected with different amounts of DNA (0.02, 0.2 and 2.0 pg/cell) of CysLTR2-GFP10 wt and CysLTR2-GFP10-GFP10, keeping the total DNA fixed at 2 pg/cell, and mock transfected cells, all in the presence of PI staining. Single parameter histograms representing expression of GFP10 and PI are shown above and to the left of each dot plot, respectively. Each plot represents a single replicate where 20,000 events in the SSC singlet gate were collected. Quadrant gating was created using untransfected control cells and single stained sample cells. High levels of receptor expression as monitored by GFP10 intensity correlates with cell death as monitored by PI staining for CysLTR2-GFP10-L129Q transfected samples but not for CysLTR2-GFP10 wt transfected samples. (**B**) Live cell imaging of CysLTR2-GFP10 wt and CysLTR2-GFP10-L129Q constructs transfected at 2 pg/cell. Cellular context of cells is shown using phase contrast. Raw and deconvolved images are optical sections extracted from the confocal stacks for wt and L129Q receptors, respectively.

**Suppl. Fig. S5.**
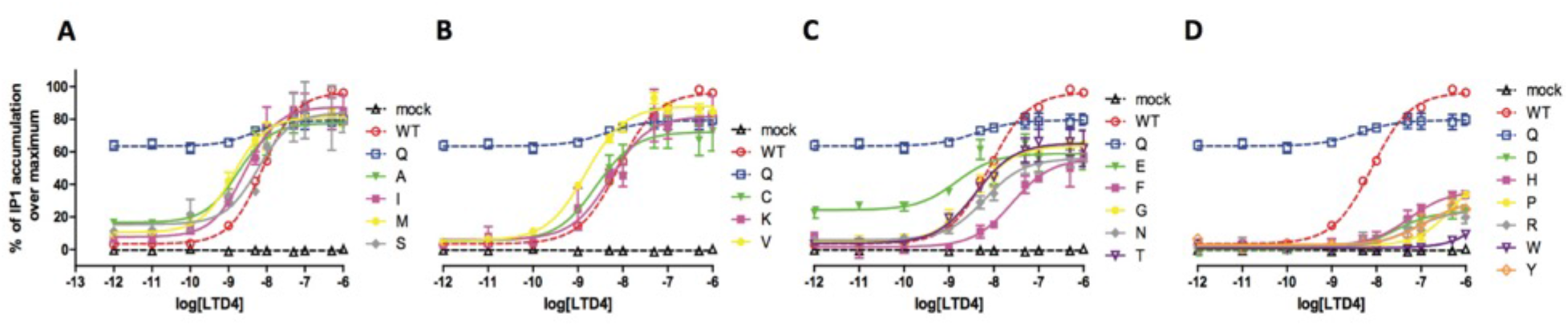
Site-saturation mutagenesis of CysLTR2-Leu129. The receptor activities were examined using agonist concentration-response curves for all twenty amino acids at residue L129 in the CysLTR2-1D4 construct transfected in HEK293T cells. (**A-D**) To reduce the complexity of the graphs, data from four to six different mutants are shown in each plot together with the empty vector transfected control (black dashed line and open triangle), CysLTR2 wild-type with native leucine (L) at position 129 (red dashed line and open circle, WT), and CysLTR2-L129Q with the oncogenic variant glutamine at position 129 (blue dashed line and open square, Q). (**A**) Compares alanine (green line and down triangle, A), isoleucine (magenta line and square, I), methionine (yellow line and circle, M), and serine (gray line and diamond, S) variants. (**B**) Compares cysteine (green line and down triangle, C), lysine (magenta line and square, K), and valine (yellow line and circle, V) variants. (**C**) Compares glutamate (green line and down triangle, E), phenylalanine (magenta line and square, F), glycine (yellow line and circle, G), asparagine (gray line and diamond, N), and threonine (purple line and open down triangle, T) variants. (**D**) Compares aspartate (green line and down triangle, D), histidine (magenta line and square, H), proline (yellow line and circle, P), arginine (gray line and diamond, R), tryptophan (purple line and open down triangle, W), and tyrosine (orange line and open diamond, Y) variants. The mutants are grouped in panels (**A-D**) according to their maximum signaling capacity. Data from experiments are shown as the normalized IP1 accumulation and are represented as the mean ± SEM of at least 2 independent experiments, each carried out in four technical replicates. The experiments were performed on four individual assay plates, which all included mock, CysLTR2-WT, -L129Q and five CysLTR2-L129^3.43^-transfected cells. Each data set was fitted to a sigmoidal dose-response model with three parameters.

**Suppl. Scheme. S1.**
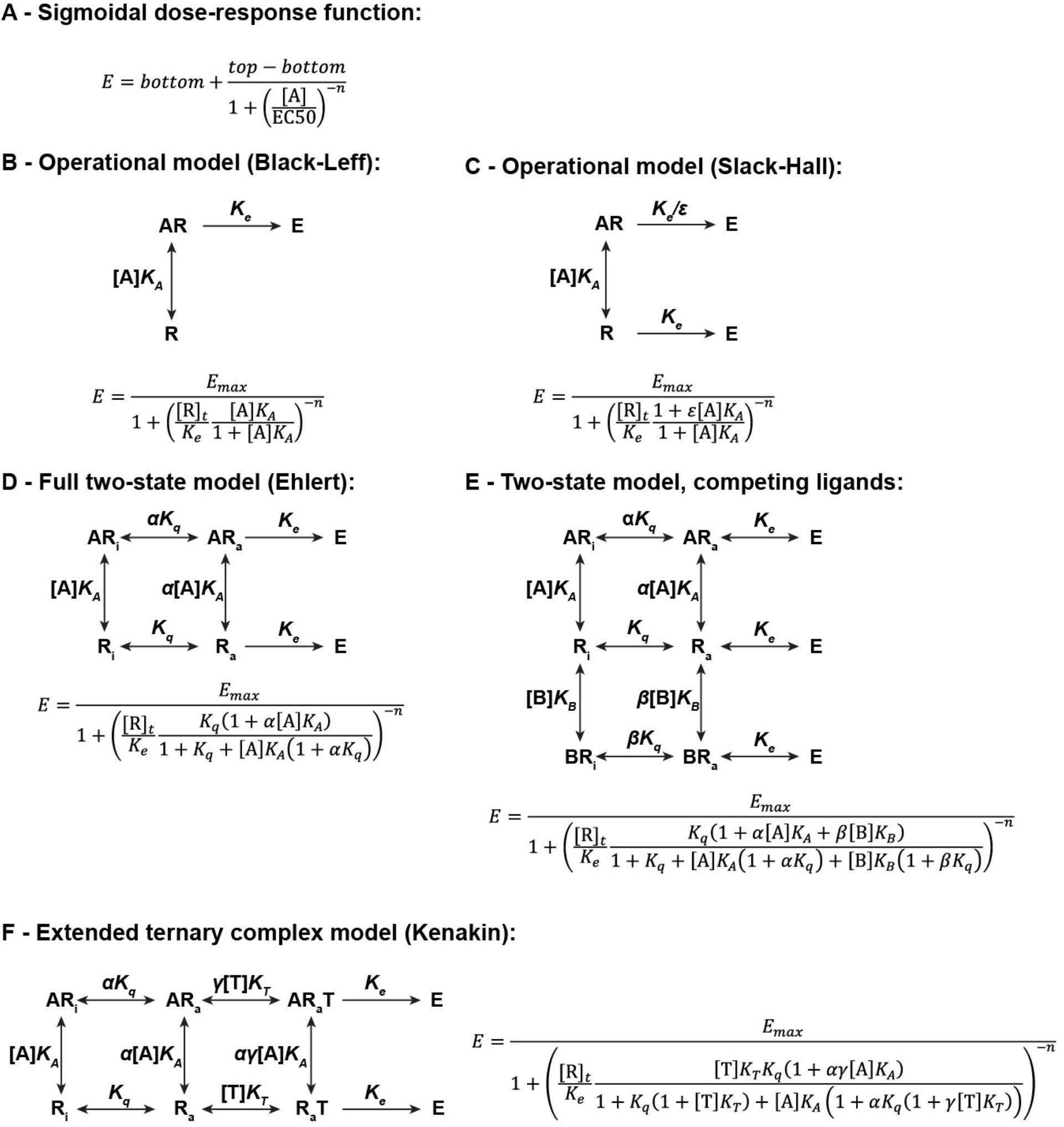
Modeling the pharmacology of constitutively active receptors. The different models and corresponding equations proposed to describe the pharmacology of the constitutively active CysLTR2-L129Q receptor are described here. (**A**) The sigmoidal dose-response model, (**B**) the Black-Leff operational model ^17^, (**C**) the Slack-Hall operational model ^18, 19^, (**D**) the Ehlert two-state allosteric model ^67^, (**E**) the modified two-state allosteric model including two competing ligands and (**F**) the extended ternary complex model ^68, 69^.

**Suppl. Fig. S6.**
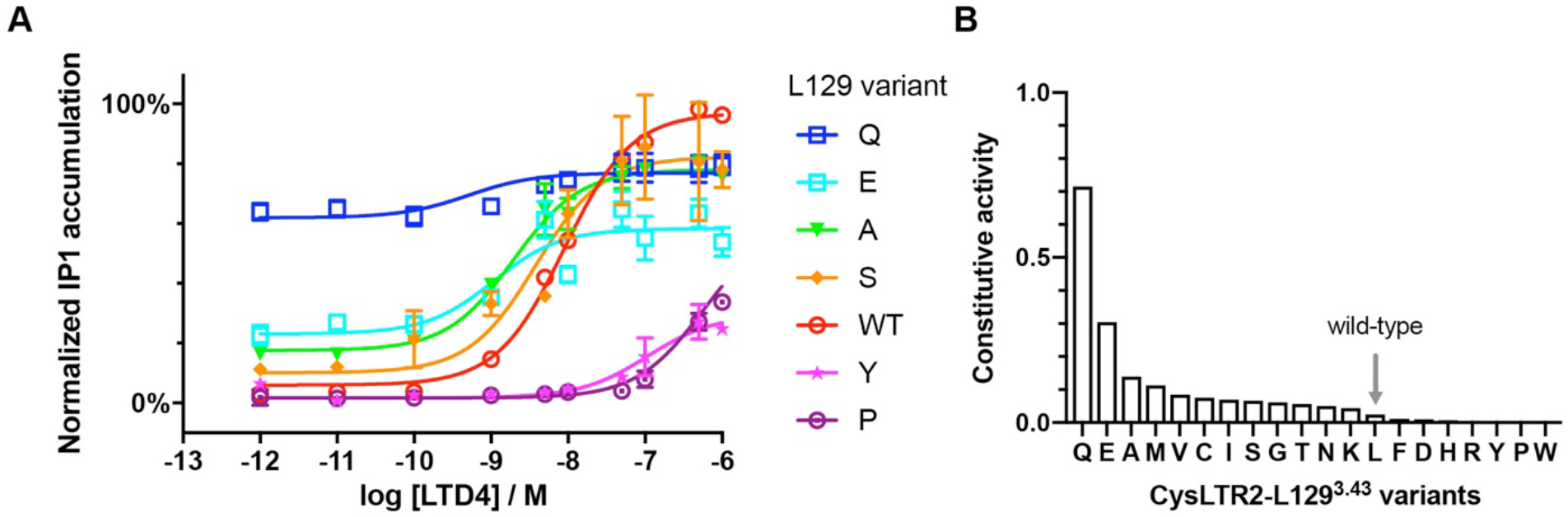
Two-state allosteric model suggests ground state equilibrium of CysLTR2-L129Q is largely shifted to the active state. (**A**) Data for selected L129 variants are shown together with curves obtained from the Ehlert two-state allosteric model using two free parameters for each mutant (τ and *K_q_*). The data are expressed as normalized IP1 and are expressed as the mean ± SEM of 2 independent experiments described in Fig. Suppl. S5, each carried out in four technical replicates. (**B**) Ranking of mutants in order of decreasing constitutive activity calculated as CA=*K_q_*/(1+*K_q_*). L129Q is the strongest CAM, and E, A, M, V, C, I, S, G, T, N, and K show constitutive IP1 activity. The grey arrow indicates CysLTR2 wild type basal level.

**Supplementary Figure S7.**
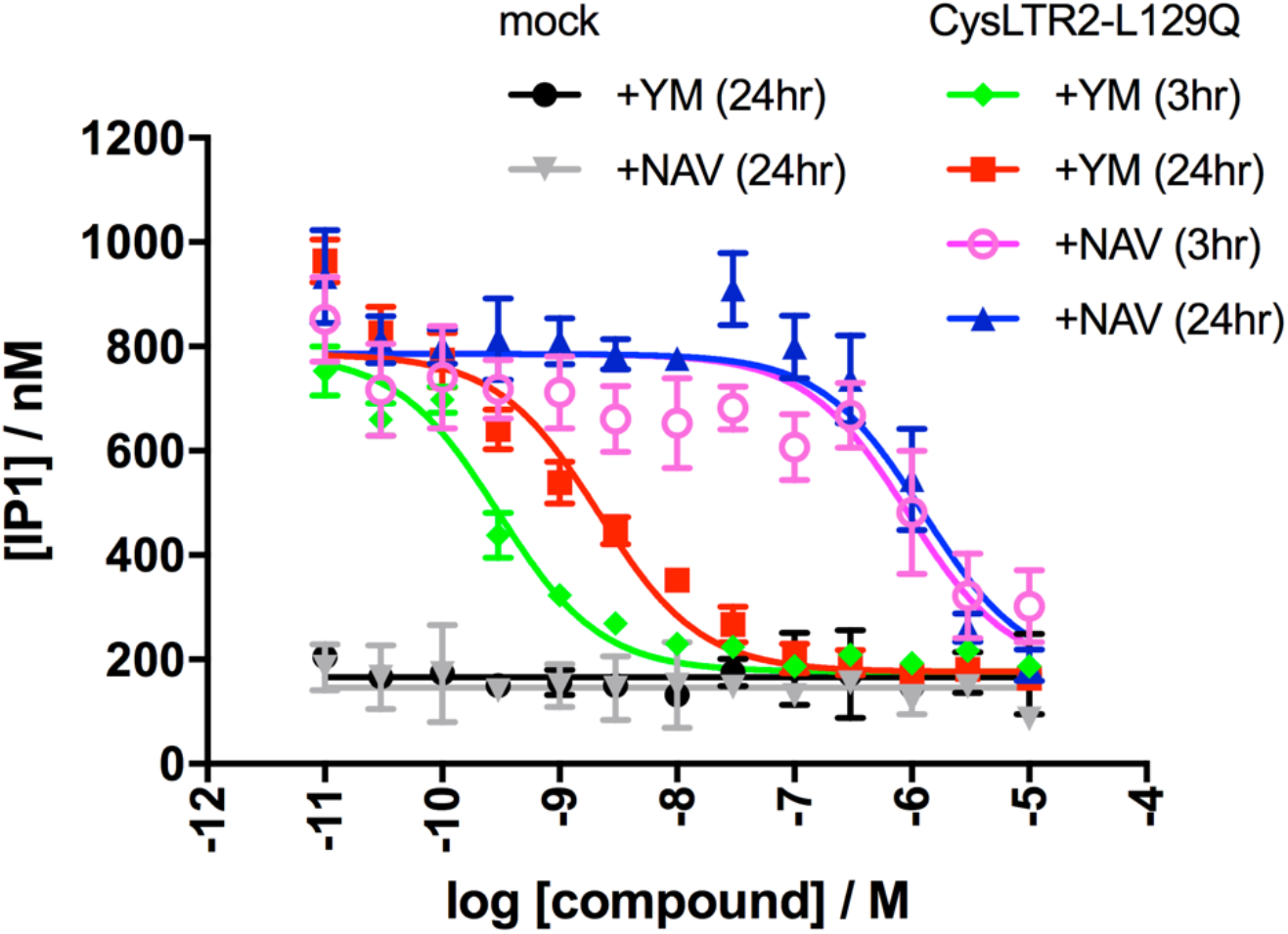
CysLTR2-L129Q signaling is blocked by Gq-inhibitor YM-254890 and Arf6-inhibitor NAV-2729. YM-254890 and NAV-2729 dose-dependently inhibits IP1 accumulation of CysLTR2-L129Q in HEK293T cells. YM-254890 fully and dose-dependently inhibits the IP1 signaling of CysLTR2-L129Q to the basal level of mock (black circles) following treatment for 3 hours (green diamonds) or 24 hours (red squares). The Arf6 inhibitor NAV-2729 strongly inhibits the constitutive IP1 signaling of CysLTR2-L129Q at doses above 1 µM, after 3 hours (pink open circle) or 24 hours treatment (blue triangle). Mock-transfected cells are not altered by any of the treatments (black circles, grey down triangle). The data are expressed as the mean ± SEM of IP1 concentration (nM) from one experiment with four technical replicates. They are fit to a sigmoidal-dose response model.

**Suppl. Table S1.** Sigmoidal versus horizontal line

|  | Bottom | Top | log EC<sub>50</sub> | Amplitude | Mean | Dof <sup>a)</sup> |
| --- | --- | --- | --- | --- | --- | --- |
| CysLTR2 wt-1D4 (Fig. 1A) |  |  |  |  |  |  |
| 11ng | 36.5±7.8 | 475.3±10.7 | -8.167±0.061 | 438.9±12.5 | **45.0±212.5** | 27 |
| 3.6ng | 31.4±5.7 | 337.5±8.5 | -8.026±0.067 | 306.1±9.6 | **45.0±147.0** | 18 |
| 1.21ng | 19.0±6.5 | 229.9±9.7 | -8.040±0.111 | 211.0±11.0 | **45.0±99.2** | 13 |
| 0.4ng | 19.1±3.9 | 161.9±6.9 | -7.748±0.107 | 142.8±7.5 | **45.0±66.5** | 8 |
| 0.1ng | 13.9±2.9 | 87.1±5.3 | -7.727±0.157 | 73.2±5.7 | **45.0±38.0** | 5 |
| CysLTR2-L129Q-1D4 (Fig. 1B) |  |  |  |  |  |  |
| 11ng | **257.9±9.7** | **274.7±11.9** | **-8.376±1.927** | **16.9±14.4** | 265.1±6.0 | 46 |
| 3.6ng | **208.1±8.9** | **209.8±8.4** | **-8.866±14.670** | **1.7±11.5** | 209.0±4.8 | 47 |
| 1.21ng | **137.3±6.4** | **149.1±7.9** | **-8.347±1.782** | **11.8±9.6** | 142.4±3.9 | 47 |
| 0.4ng | **93.7±4.9** | **100.8±5.4** | **-8.571±2.130** | **7.1±6.8** | 97.0±2.8 | 47 |
| 0.1ng | **56.1±2.0** | **61.9±4.4** | **-7.419±1.474** | **5.8±4.6** | 57.7±1.4 | 47 |
| CysLTR2-GFP10 wt (Fig. S3A) |  |  |  |  |  |  |
| 11ng | 43.7±4.2 | 339.5±7.6 | -7.730±0.056 | 295.8±8.2 | **141.0±16.9** | 47 |
| 3.6ng | 42.5±5.3 | 244.0±9.8 | -7.700±0.104 | 201.5±10.5 | **107.8±11.9** | 47 |
| 1.21ng | 39.0±3.8 | 185.9±7.7 | -7.537±0.107 | 147.0±8.1 | **82.6±8.4** | 47 |
| 0.4ng | 35.4±2.8 | 114.4±5.9 | -7.493±0.149 | 79.0±6.2 | **58.3±4.7** | 47 |
| 0.1ng | 36.3±1.7 | 75.1±4.4 | -7.253±0.202 | 38.8±4.4 | **46.1±2.3** | 47 |
| CysLTR2-GFP10-L129Q (Fig. S3B) |  |  |  |  |  |  |
| 11ng | **N.C.** | **322.3±8.1** | **N.C.** | **N.C.** | 322.3±7.9 | 47 |
| 3.6ng | 246.5±8.6 | 278.6±9.5 | -8.586±0.828 | 32.1±12.1 | **261.6±5.3** | 47 |
| 1.21ng | 176.3±7.2 | 205.9±8.1 | -8.552±0.758 | 29.6±10.2 | **190.0±4.6** | 47 |
| 0.4ng | **107.5±7.7** | **124.8±10.0** | **-8.277±1.492** | **17.4±11.9** | 114.7±4.9 | 47 |
| 0.1ng | **64.9±5.8** | **70.6±4.6** | **-9.215±2.713** | **5.6±7.0** | 68.2±2.8 | 47 |
| pcDNA | **-0.3±0.9** | **-0.7±0.6** | **-9.398±7.626** | **-0.3±1.0** | -0.6±0.4 | 77 |
| WT | 3.5±0.8 | 96.5±0.9 | -8.103±0.024 | 93.0±1.1 | **48.5±4.3** | 79 |
| Q | 63.5±1.9 | 79.3±1.8 | -8.391±0.319 | 15.9±2.5 | **72.0±1.3** | 79 |
| A | 16.6±2.0 | 77.7±1.6 | -8.761±0.089 | 61.1±2.4 | **53.0±5.9** | 19 |
| C | 5.4±3.8 | 72.3±3.4 | -8.597±0.155 | 66.9±4.8 | **43.6±6.7** | 19 |
| D | 0.3±1.2 | 23.7±1.8 | -7.570±0.174 | 23.4±2.0 | **9.2±2.2** | 19 |
| E | 24.2±3.3 | 58.9±2.7 | -8.831±0.263 | 34.6±4.0 | **45.2±3.1** | 29 |
| F | 2.4±2.1 | 55.9±3.4 | -7.565±0.140 | 53.5±3.7 | **22.8±5.0** | 19 |
| G | 4.5±1.9 | 63.7±1.8 | -8.420±0.086 | 59.2±2.5 | **36.5±5.7** | 19 |
| H | 2.3±1.3 | 35.8±2.5 | -7.267±0.142 | 33.5±2.6 | **13.1±2.9** | 19 |
| I | 7.7±3.6 | 87.6±3.2 | -8.577±0.123 | 79.9±4.6 | **53.0±7.9** | 19 |
| K | 6.1±4.3 | 82.0±4.4 | -8.277±0.156 | 75.8±5.9 | **45.3±7.6** | 19 |
| M | 10.4±2.1 | 81.4±1.6 | -8.883±0.081 | 71.0±2.5 | **54.0±6.9** | 19 |
| N | 5.8±1.7 | 55.8±1.8 | -8.200±0.094 | 50.0±2.4 | **30.9±4.8** | 19 |
| P | 1.8±0.7 | 55.4±10.2 | -6.198±0.179 | 53.5±10.0 | **8.7±2.6** | 19 |
| R | 1.3±1.6 | 24.2±3.3 | -7.197±0.263 | 22.9±3.4 | **8.4±2.2** | 19 |
| S | 15.5±5.2 | 84.0±5.4 | -8.228±0.207 | 68.5±7.1 | **50.2±7.1** | 19 |
| T | 3.8±3.0 | 65.2±2.9 | -8.407±0.131 | 61.5±3.9 | **36.9±6.1** | 19 |
| V | 5.6±1.7 | 88.0±1.4 | -8.828±0.058 | 82.4±2.1 | **55.6±7.9** | 19 |
| W | **0.9±0.5** | **N.C.** | **N.C.** | **N.C.** | 2.4±0.7 | 19 |
| Y | 3.0±1.2 | 30.0±3.3 | -6.889±0.199 | 27.0±3.3 | **9.8±2.2** | 19 |
| CysLTR2 wt-1D4 |  |  |  |  |  |  |
| WT | 126.7±30.7 | 1792.0±54.8 | -7.754±0.065 | 1665.0±58.1 | **704.8±122.5** | 26 |
| WT N301A | **49.0±4.5** | **N.C.** | **N.C.** | **N.C.** | 51.6±3.5 | 27 |
| WT Y305F | 48.5±12.4 | 961.1±79.7 | -6.705±0.102 | 912.7±77.4 | **201.0±45.0** | 27 |
| WT/NA/YF | **N.C.** | **49.3±3.5** | **N.C.** | **N.C.** | 48.5±2.7 | 27 |
| L129Q-1D4 | **1156.0±19.5** | **1236.0±88.9** | **-6.874±1.458** | **80.5±86.6** | 1171.0±14.7 | 27 |
| LQ/N301A | 968.0±22.8 | 1279.0±50.8 | -7.409±0.298 | 310.7±51.7 | **1061.0±21.9** | 48 |
| LQ/Y305F | 974.4±27.7 | 1607.0±37.6 | -7.987±0.138 | 632.8±43.1 | **1238.0±37.4** | 55 |
| LQ/N301A/Y305F | 359.0±15.2 | 1144.0±43.5 | -7.193±0.090 | 785.0±43.2 | **561.6±50.6** | 27 |

**Bold** = Preferred model is sigmoidal dose-response TH add amplitude, Normal = Preferred model is horizontal line.
a) Degrees of Freedom, N.C. = Non-converging

**Suppl. Table S2.** Black-Leff Model

| | log KA | log $\tau$ | Basal | E<sub>max</sub> | Dof <sup>a)</sup> |
| --- | --- | --- | --- | --- | --- |
| CysLTR2 wt-1D4 (Fig. 1A) |  |  |  |  |  |
| 11ng |  | 0.259 ± 0.153 | 22.9 ± 0.013 |  |  |
| 3.6ng |  | -0.080 ± 0.112 | 22.9 ± 0.009 |  |  |
| 1.21ng |  | -0.357 ± 0.093 | 22.9 ± 0.022 |  |  |
| 0.4ng |  | -0.629 ± 0.085 | 22.9 ± 0.024 |  |  |
| 0.1ng |  | -1.027 ± 0.092 | 22.9 ± 0.080 |  |  |
| Global (shared) | -7.780 ± 0.081 |  | 22.9 ± 0.019 | 718.0 ± 92.5 | 232 |
| CysLTR2-GFP10 wt (Fig. S3A) |  |  |  |  |  |
| 11ng |  | 0.038 ± 0.237 | 38.9 ± 0.027 |  |  |
| 3.6ng |  | -0.235 ± 0.193 | 38.9 ± 0.065 |  |  |
| 1.21ng |  | -0.468 ± 0.170 | 38.9 ± 0.023 |  |  |
| 0.4ng |  | -0.828 ± 0.153 | 38.9 ± 0.025 |  |  |
| 0.1ng |  | -1.229 ± 0.161 | 38.9 ± 0.022 |  |  |
| Global (shared) | -7.454 ± 0.104 |  | 38.9 ± 0.030 | 608.6 ± 154.6 | 232 |

a) Degrees of Freedom

**Suppl. Table S3.** Slack-Hall Operational Model

| | log KA | log $\chi$ | log $\epsilon$ | Basal | E<sub>max</sub> | log $\tau$ | Dof <sup>a)</sup> |
| --- | --- | --- | --- | --- | --- | --- | --- |
| CysLTR2 wt-1D4 (Fig. 1A) |  |  |  |  |  |  |  |
| 11ng |  | -1.503 ± 0.135 |  |  |  | 0.048 ± 0.198 |  |
| 3.6ng |  | -1.780 ± 0.127 |  |  |  | -0.230 ± 0.157 |  |
| 1.21ng |  | -2.022 ± 0.129 |  |  |  | -0.472 ± 0.135 |  |
| 0.4ng |  | -2.263 ± 0.137 |  |  |  | -0.713 ± 0.121 |  |
| 0.1ng |  | -2.606 ± 0.158 |  |  |  | -1.056 ± 0.113 |  |
| Global (fixed) |  |  |  |  |  |  |  |
| Global (shared) | 7.847 ± 0.083 |  | 1.550 ± 0.186 | 12.3 ± 4.5 | 877.9 ± 200 |  | 231 |
| CysLTR2-L129Q-1D4 (Fig. 1B) |  |  |  |  |  |  |  |
| 11ng |  | -0.323 ± 0.015 |  |  |  | -0.287 ± 0.017 |  |
| 3.6ng |  | -0.494 ± 0.017 |  |  |  | -0.458 ± 0.018 |  |
| 1.21ng |  | -0.755 ± 0.022 |  |  |  | -0.719 ± 0.023 |  |
| 0.4ng |  | -1.024 ± 0.033 |  |  |  | -0.989 ± 0.033 |  |
| 0.1ng |  | -1.484 ± 0.080 |  |  |  | -1.449 ± 0.080 |  |
| Global (fixed) |  |  |  | = 34.75 | = 697.4 |  |  |
| Global (shared) | 8.433 ± 1.158 |  | 0.035 ± 0.018 |  |  |  | 232 |
| CysLTR2-GFP10 wt (Fig. S3A) |  |  |  |  |  |  |  |
| 11ng |  | -1.828 ± 0.238 |  |  |  | -0.113 ± 0.299 |  |
| 3.6ng |  | -2.070 ± 0.222 |  |  |  | -0.355 ± 0.254 |  |
| 1.21ng |  | -2.284 ± 0.219 |  |  |  | -0.569 ± 0.228 |  |
| 0.4ng |  | -2.617 ± 0.225 |  |  |  | -0.901 ± 0.205 |  |
| 0.1ng |  | -2.972 ± 0.250 |  |  |  | -1.257 ± 0.197 |  |
| Global (fixed) |  |  |  |  |  |  |  |
| Global (shared) | 7.494 ± 0.106 |  | 1.715 ± 0.277 | 34.8 ± 2.9 | 697.4 ± 280.3 |  | 231 |
| CysLTR2-GFP10-L129Q (Fig. S3B) |  |  |  |  |  |  |  |
| 11ng |  | -0.186 ± 0.019 |  |  |  | -0.125 ± 0.018 |  |
| 3.6ng |  | -0.349 ± 0.020 |  |  |  | -0.288 ± 0.019 |  |
| 1.21ng |  | -0.575 ± 0.023 |  |  |  | -0.514 ± 0.022 |  |
| 0.4ng |  | -0.919 ± 0.035 |  |  |  | -0.859 ± 0.034 |  |
| 0.1ng |  | -1.330 ± 0.073 |  |  |  | -1.270 ± 0.073 |  |
| Global (fixed) |  |  |  | = 34.75 | = 697.4 |  |  |
| Global (shared) | 8.879 ± 0.732 |  | 0.061 ± 0.020 |  |  |  | 233 |
| pcDNA |  |  |  |  |  |  |  |
| WT | 7.825 ± 0.025 | -1.753 ± 0.101 | 1.722 ± 0.101 |  |  | -0.031 ± 0.008 | 77 |
| Q | 8.338 ± 0.320 | -0.333 ± 0.019 | 0.151 ± 0.024 |  |  | -0.182 ± 0.016 | 77 |
| A | 8.585 ± 0.090 | -1.043 ± 0.056 | 0.846 ± 0.057 |  |  | -0.197 ± 0.015 | 17 |
| C | 8.414 ± 0.158 | -1.557 ± 0.313 | 1.310 ± 0.311 |  |  | -0.247 ± 0.032 | 17 |
| D | 7.516 ± 0.175 | -2.900 ± 2.020 | 2.027 ± 2.014 |  |  | -0.873 ± 0.038 | 17 |
| E | 8.735 ± 0.263 | -0.861 ± 0.067 | 0.481 ± 0.070 |  |  | -0.380 ± 0.028 | 27 |
| F | 7.427 ± 0.144 | -1.919 ± 0.397 | 1.508 ± 0.393 |  |  | -0.411 ± 0.036 | 17 |
| G | 8.263 ± 0.088 | -1.641 ± 0.187 | 1.311 ± 0.187 |  |  | -0.330 ± 0.018 | 17 |
| H | 7.186 ± 0.146 | -1.940 ± 0.244 | 1.279 ± 0.241 |  |  | -0.662 ± 0.037 | 17 |
| I | 8.344 ± 0.127 | -1.398 ± 0.211 | 1.289 ± 0.210 |  |  | -0.109 ± 0.028 | 17 |
| K | 8.062 ± 0.161 | -1.499 ± 0.315 | 1.341 ± 0.314 |  |  | -0.159 ± 0.040 | 17 |
| M | 8.679 ± 0.082 | -1.259 ± 0.091 | 1.096 ± 0.091 |  |  | -0.164 ± 0.015 | 17 |
| N | 8.071 ± 0.095 | -1.525 ± 0.132 | 1.112 ± 0.131 |  |  | -0.413 ± 0.019 | 17 |
| P | 6.061 ± 0.208 | -2.031 ± 0.176 | 1.614 ± 0.175 |  |  | -0.417 ± 0.111 | 17 |
| R | 7.144 ± 0.267 | -2.191 ± 0.533 | 1.331 ± 0.527 |  |  | -0.861 ± 0.067 | 17 |
| S | 8.026 ± 0.213 | -1.077 ± 0.158 | 0.937 ± 0.161 |  |  | -0.140 ± 0.048 | 17 |
| T | 8.244 ± 0.133 | -1.718 ± 0.349 | 1.403 ± 0.347 |  |  | -0.315 ± 0.028 | 17 |
| V | 8.588 ± 0.060 | -1.541 ± 0.139 | 1.437 ± 0.139 |  |  | -0.105 ± 0.012 | 17 |
| W | N.C. | -1.925 ± 0.145 | N.C. |  |  | N.C. | 17 |
| Y | 6.825 ± 0.205 | -1.824 ± 0.174 | 1.071 ± 0.171 |  |  | -0.754 ± 0.057 | 17 |
| Global (fixed) |  |  |  | = 0.000 | = 200.0 |  |  |
| CysLTR2 wt-1D4 |  |  |  |  |  |  |  |
| WT | 7.445 ± 0.075 | -1.412 ± 0.109 | 1.459 ± 0.108 |  |  | 0.047 ± 0.028 | 24 |
| WT N301A | N.C. | -1.812 ± 0.031 | N.C. |  |  | N.C. | 25 |
| WT Y305F | 6.567 ± 0.114 | -1.840 ± 0.112 | 1.436 ± 0.111 |  |  | -0.404 ± 0.050 | 25 |
| WT/NA/YF | N.C. | N.C. | N.C. |  |  | N.C. | 25 |
| L129Q-1D4 | 6.858 ± 1.472 | -0.288 ± 0.011 | 0.045 ± 0.048 |  |  | -0.243 ± 0.049 | 25 |
| LQ/N301A | 7.350 ± 0.304 | -0.400 ± 0.014 | 0.180 ± 0.029 |  |  | -0.220 ± 0.028 | 46 |
| LQ/Y305F | 7.855 ± 0.142 | -0.396 ± 0.017 | 0.349 ± 0.024 |  |  | -0.047 ± 0.019 | 53 |
| LQ/N301A/Y305F | 7.063 ± 0.096 | -0.928 ± 0.021 | 0.633 ± 0.029 |  |  | -0.295 ± 0.025 | 25 |
| Global (fixed) |  |  |  | = 0.000 | = 3400 |  |  |

a) Degrees of Freedom

**Suppl. Table S4.** Ehlert Two-State Allosteric Model

| | Basal | E<sub>max</sub> | log $\tau$ | log K<sub>q</sub> | log $\alpha$ | log K<sub>a</sub> | Dof<sup>a)</sup> |
| --- | --- | --- | --- | --- | --- | --- | --- |
| CysLTR2 wt-1D4 (Fig. 1A) |  |  |  |  |  |  |  |
| 11ng |  |  | 0.097 ± 0.169 |  |  |  |  |
| 3.6ng |  |  | -0.192 ± 0.133 |  |  |  |  |
| 1.21ng |  |  | -0.442 ± 0.114 |  |  |  |  |
| 0.4ng |  |  | -0.689 ± 0.104 |  |  |  |  |
| 0.1ng |  |  | -1.041 ± 0.103 |  |  |  |  |
| Global (fixed) |  |  |  |  | = 4.480 | = 5.000 |  |
| Global (shared) | 14.1 ± 2.9 | 843.0 ± 149.5 |  | -1.637 ± 0.082 |  |  | 232 |
| CysLTR2-L129Q-1D4 (Fig. 1B) |  |  |  |  |  |  |  |
| 11ng |  |  | -0.345 ± 0.013 |  |  |  |  |
| 3.6ng |  |  | -0.496 ± 0.015 |  |  |  |  |
| 1.21ng |  |  | -0.720 ± 0.018 |  |  |  |  |
| 0.4ng |  |  | -0.937 ± 0.025 |  |  |  |  |
| 0.1ng |  |  | -1.239 ± 0.043 |  |  |  |  |
| Global (fixed) | = 14.1 | = 843.0 |  |  |  |  |  |
| Global (shared) |  |  |  | 1.138 ± 0.231 |  |  | 233 |
| CysLTR2-GFP10 wt (Fig. S3A) |  |  |  |  |  |  |  |
| 11ng |  |  | -0.025 ± 0.252 |  |  |  |  |
| 3.6ng |  |  | -0.283 ± 0.209 |  |  |  |  |
| 1.21ng |  |  | -0.508 ± 0.186 |  |  |  |  |
| 0.4ng |  |  | -0.854 ± 0.168 |  |  |  |  |
| 0.1ng |  |  | -1.230 ± 0.171 |  |  |  |  |
| Global (fixed) |  |  |  |  | = 4.480 | = 5.000 |  |
| Global (shared) | 36.7 ± 1.7 | 656.7 ± 191.0 |  | -2.004 ± 0.104 |  |  | 232 |
| CysLTR2-GFP10-L129Q (Fig. S3B) |  |  |  |  |  |  |  |
| 11ng |  |  | -0.044 ± 0.017 |  |  |  |  |
| 3.6ng |  |  | -0.220 ± 0.018 |  |  |  |  |
| 1.21ng |  |  | -0.458 ± 0.022 |  |  |  |  |
| 0.4ng |  |  | -0.817 ± 0.034 |  |  |  |  |
| 0.1ng |  |  | -1.246 ± 0.077 |  |  |  |  |
| Global (fixed) | = 36.7 | = 656.7 |  |  | = 4.480 | = 5.000 |  |
| Global (shared) |  |  |  | 0.792 ± 0.162 |  |  | 234 |
| pcDNA |  |  |  |  |  |  |  |
| WT |  |  | -0.029 ± 0.013 | -1.655 ± 0.066 |  |  |  |
| Q |  |  | -0.206 ± 0.009 | 0.411 ± 0.067 |  |  |  |
| A |  |  | -0.199 ± 0.022 | -0.798 ± 0.081 |  |  |  |
| C |  |  | -0.241 ± 0.024 | -1.118 ± 0.099 |  |  |  |
| D |  |  | -0.871 ± 0.080 | -1.963 ± 0.330 |  |  |  |
| E |  |  | -0.390 ± 0.019 | -0.348 ± 0.076 |  |  |  |
| F |  |  | -0.415 ± 0.040 | -1.999 ± 0.154 |  |  |  |
| G |  |  | -0.328 ± 0.027 | -1.218 ± 0.114 |  |  |  |
| H |  |  | -0.669 ± 0.065 | -2.194 ± 0.237 |  |  |  |
| I |  |  | -0.104 ± 0.023 | -1.162 ± 0.086 |  |  |  |
| K |  |  | -0.160 ± 0.025 | -1.380 ± 0.098 |  |  |  |
| M |  |  | -0.153 ± 0.022 | -0.912 ± 0.081 |  |  |  |
| N |  |  | -0.418 ± 0.030 | -1.304 ± 0.131 |  |  |  |
| P |  |  | -0.452 ± 0.232 | -3.288 ± 0.450 |  |  |  |
| R |  |  | -0.869 ± 0.094 | -2.240 ± 0.345 |  |  |  |
| S |  |  | -0.162 ± 0.024 | -1.177 ± 0.093 |  |  |  |
| T |  |  | -0.312 ± 0.027 | -1.255 ± 0.113 |  |  |  |
| V |  |  | -0.090 ± 0.022 | -1.060 ± 0.081 |  |  |  |
| W |  |  | -0.550 ± 4.717 | -4.116 ± 5.557 |  |  |  |
| Y |  |  | -0.770 ± 0.094 | -2.479 ± 0.307 |  |  |  |
| Global (fixed) | = 0.0 | = 200.0 |  |  |  | = 5.000 |  |
| Global (shared) | | | | | $4.480 \pm 0.062$ | | 489 |
| CysLTR2 wt-1D4 |  |  |  |  |  |  |  |
| WT | | | $0.149 \pm 0.034$ | $-1.989 \pm 0.073$ | | | 25 |
| WT N301A | | | $-1.747 \pm 0.039$ | $1.067 \pm 1.092$ | | | 26 |
| WT Y305F | | | $-0.362 \pm 0.051$ | $-2.792 \pm 0.112$ | | | 26 |
| WT/NA/YF | | | $-1.774 \pm 0.032$ | $1.088 \pm 0.929$ | | | 26 |
| L129Q-1D4 | | | $-0.186 \pm 0.011$ | $1.206 \pm 0.416$ | | | 26 |
| LQ/N301A | | | $-0.223 \pm 0.016$ | $0.454 \pm 0.131$ | | | 47 |
| LQ/Y305F | | | $-0.040 \pm 0.020$ | $-0.119 \pm 0.075$ | | | 54 |
| LQ/N301A/Y305F | | | $-0.448 \pm 0.042$ | $-0.637 \pm 0.155$ | | | 26 |
| Global (fixed) | = 0.0 | = 3000 |  |  | = 4.480 | = 5.000 |  |

a) Degrees of Freedom

**Suppl. Table S5.**
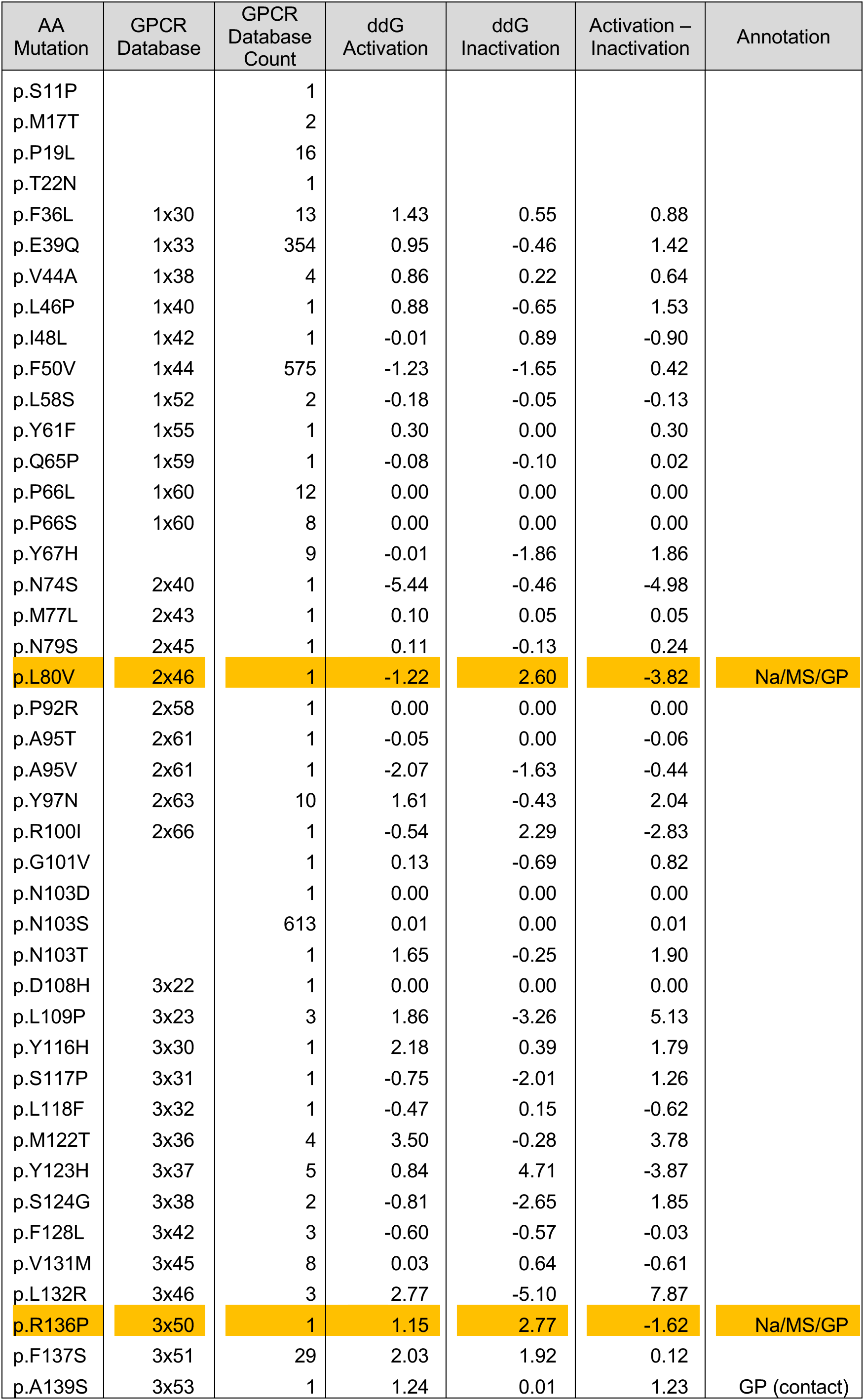
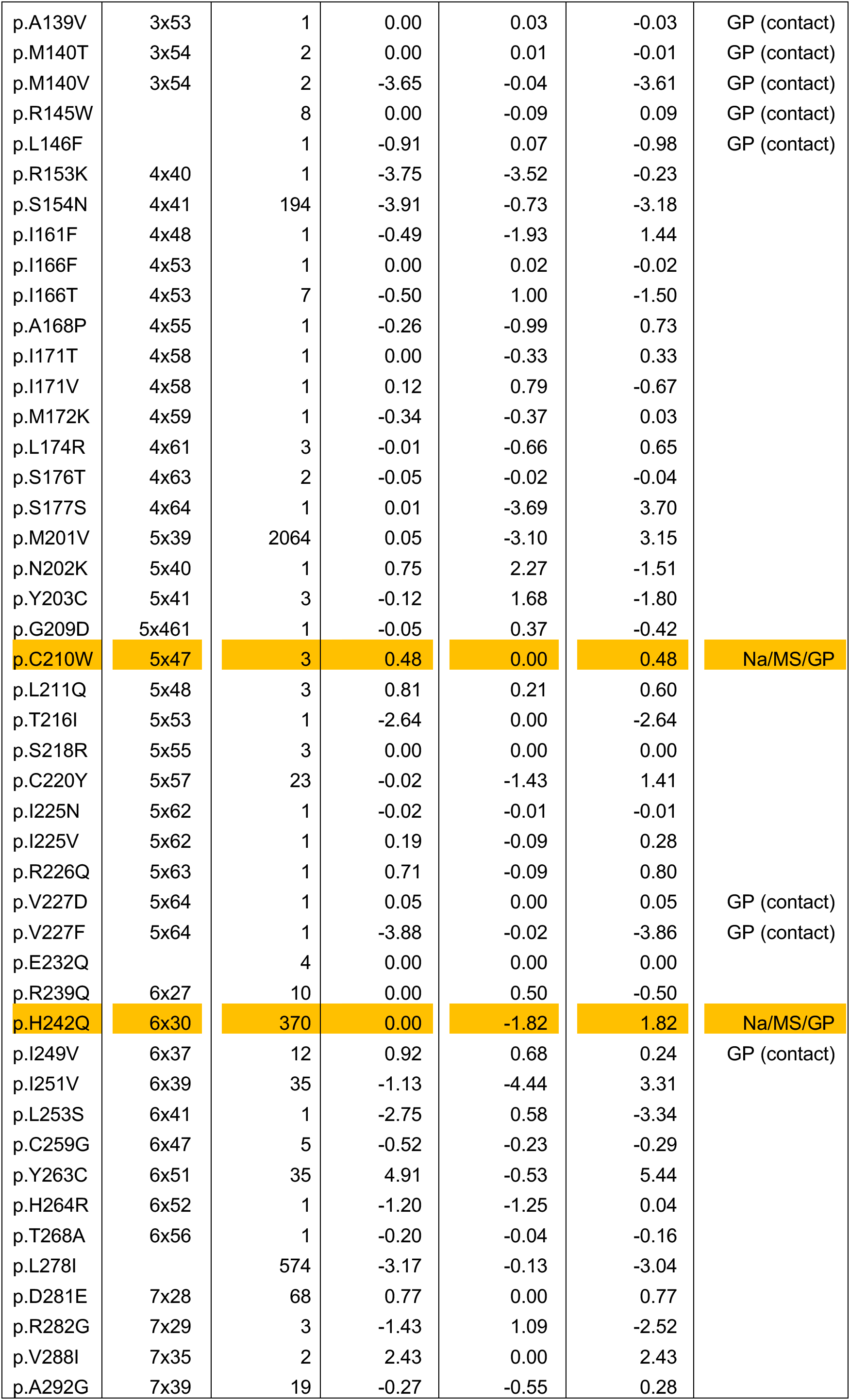
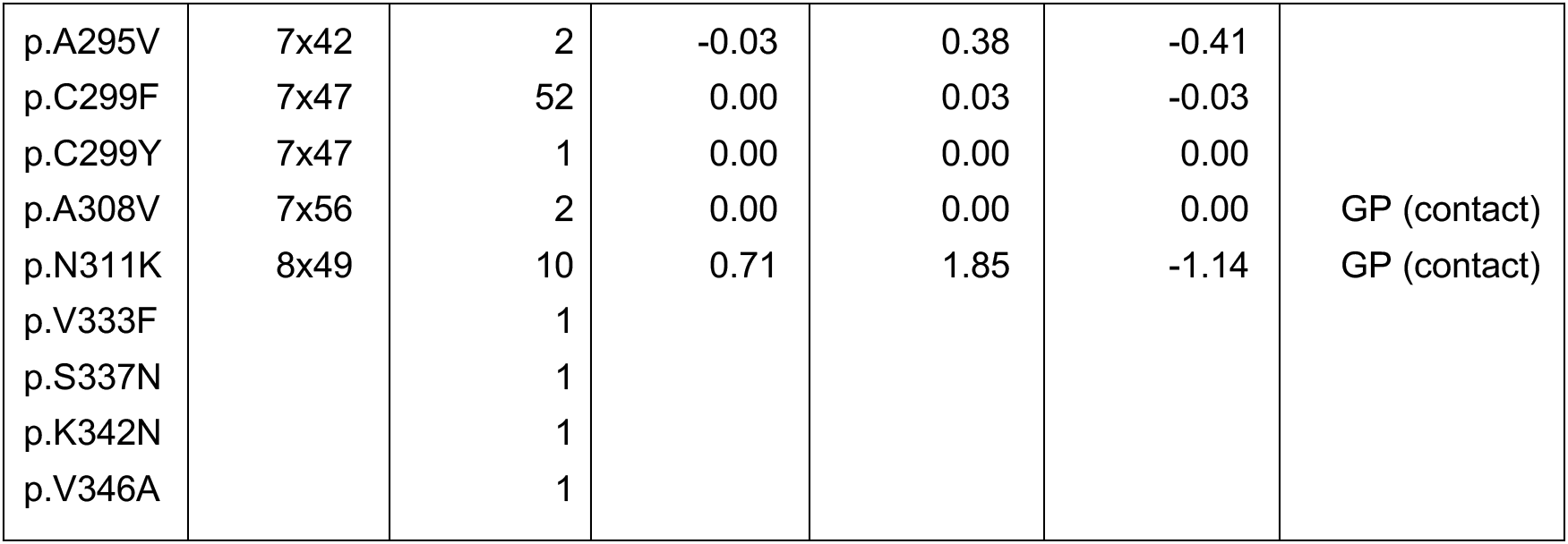
“Normal” Mutations found only as germline MVs listed in the GPCRdb

**Suppl. Table S6.** Mutations found “both” as germline and somatic variants

| AA Mutation | GPCR Database | GPCR Database Count | COSMIC +TCGA | COSMIC Count | TCGA Count | ddG Activation | ddG Inactivation | Activation – Inactivation | Annotation |
| --- | --- | --- | --- | --- | --- | --- | --- | --- | --- |
| p.S70F |  | 1 | 1 |  | 1 | -1.00 | -1.54 | 0.53 |  |
| p.Y119H | 3x33 | 1 | 1 |  | 1 | 3.36 | 4.00 | -0.65 |  |
| p.R136C | 3x50 | 2 | 1 | 1 |  | 2.41 | 2.12 | 0.29 | Na/MS/GP |
| p.R145Q |  | 24 | 1 |  | 1 | 0.40 | 0.03 | 0.37 |  |
| p.A205T | 5x43 | 1 | 1 |  | 1 | 0.64 | 1.96 | -1.32 |  |
| p.S236L | 6x24 | 1139 | 1 | 1 |  | 5.54 | 0.26 | 5.28 |  |
| p.V269I | 6x57 | 6 | 1 | 1 |  | -1.43 | 0.32 | -1.76 |  |
| p.A298T | 7x46 | 1 | 1 | 1 |  | 0.28 | 1.45 | -1.18 |  |
| p.R315K | 8x53 | 265 | 1 |  | 1 | 0.97 | 1.60 | -0.63 |  |
| p.V14I |  | 5 | 2 | 2 |  |  |  |  |  |
| p.P42L | 1x36 | 1 | 2 | 2 |  | -4.54 | -5.37 | 0.83 |  |
| p.P42S | 1x36 | 1 | 2 | 1 | 1 | 0.15 | 0.30 | -0.14 |  |
| p.V75I | 2x41 | 1 | 2 | 1 | 1 | 1.31 | 0.35 | 0.96 |  |
| p.T90M | 2x56 | 19 | 3 | 1 | 2 | -6.25 | -0.18 | -6.07 |  |
| p.M114I | 3x28 | 4 | 3 | 2 | 1 | 3.14 | -0.50 | 3.64 |  |
| p.R136H | 3x50 | 2 | 2 |  | 2 | 0.52 | -1.39 | 1.92 | Na/MS/GP |
| p.R226W | 5x63 | 3 | 4 | 3 | 1 | -3.05 | -1.38 | -1.67 |  |
| p.R239W | 6x27 | 4 | 3 | 2 | 1 | -2.42 | -7.46 | 5.04 |  |
| p.T272M | 6x60 | 17 | 5 | 2 | 3 | -4.02 | -1.98 | -2.04 |  |

**Suppl. Table S7.**
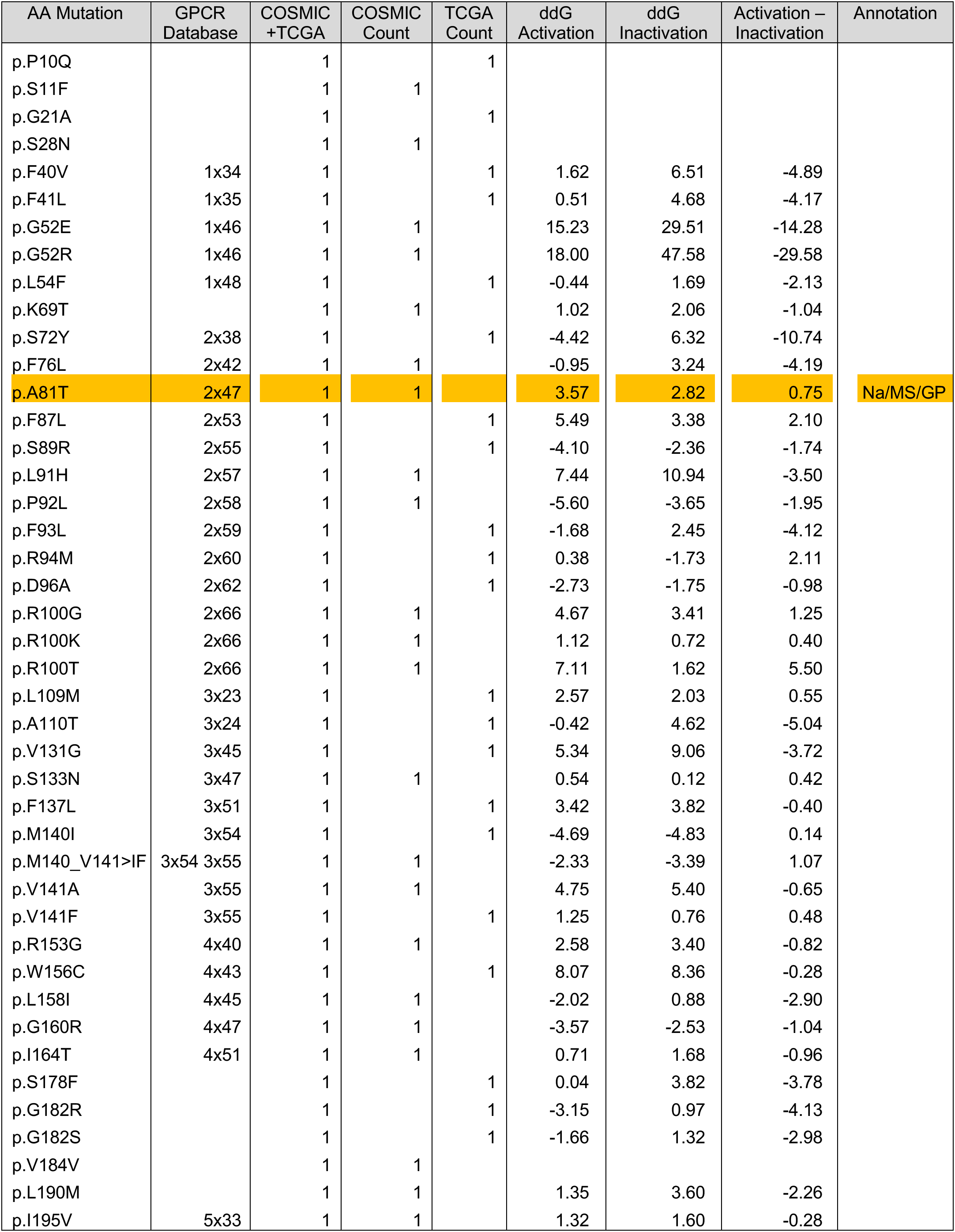
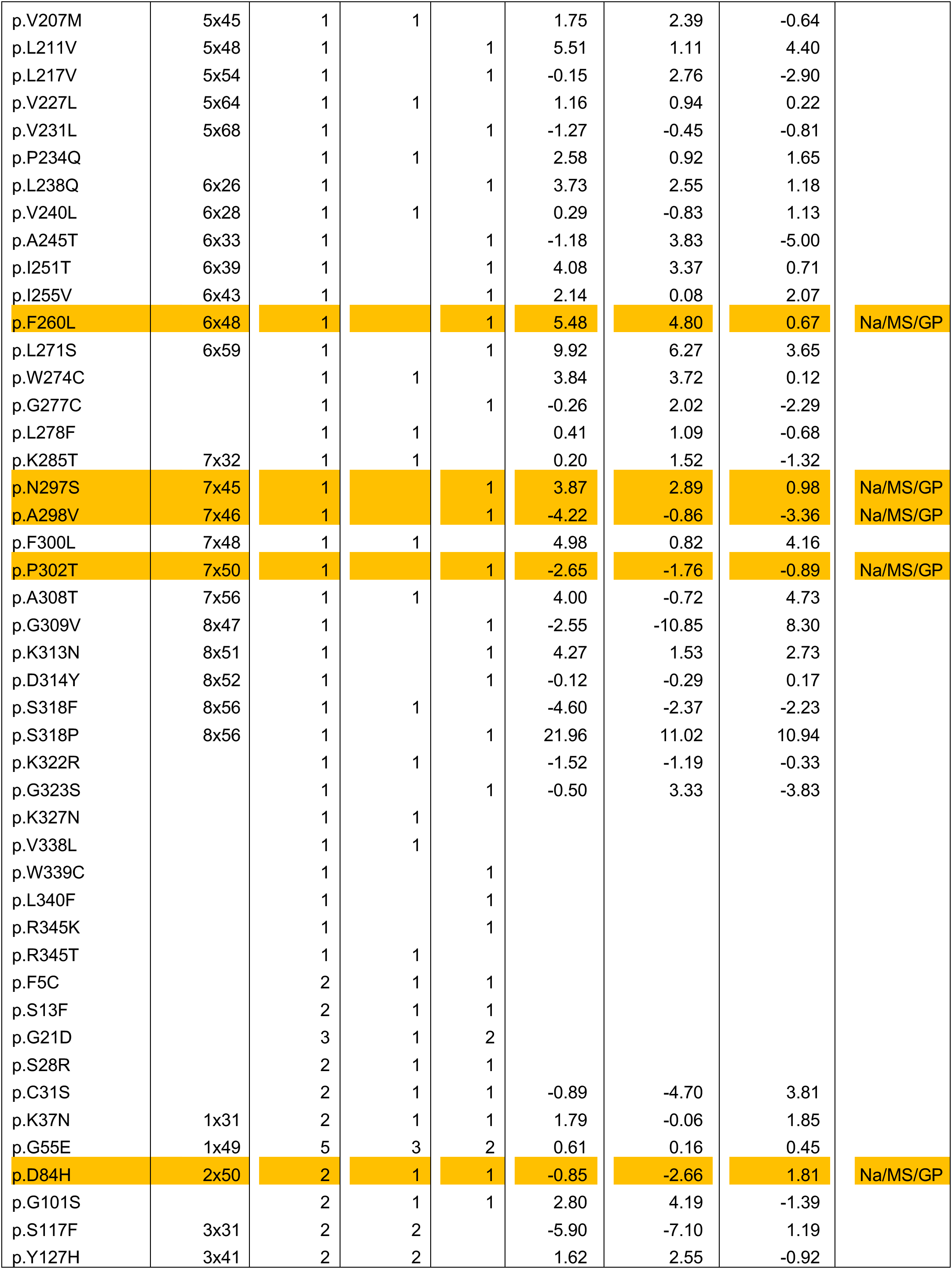
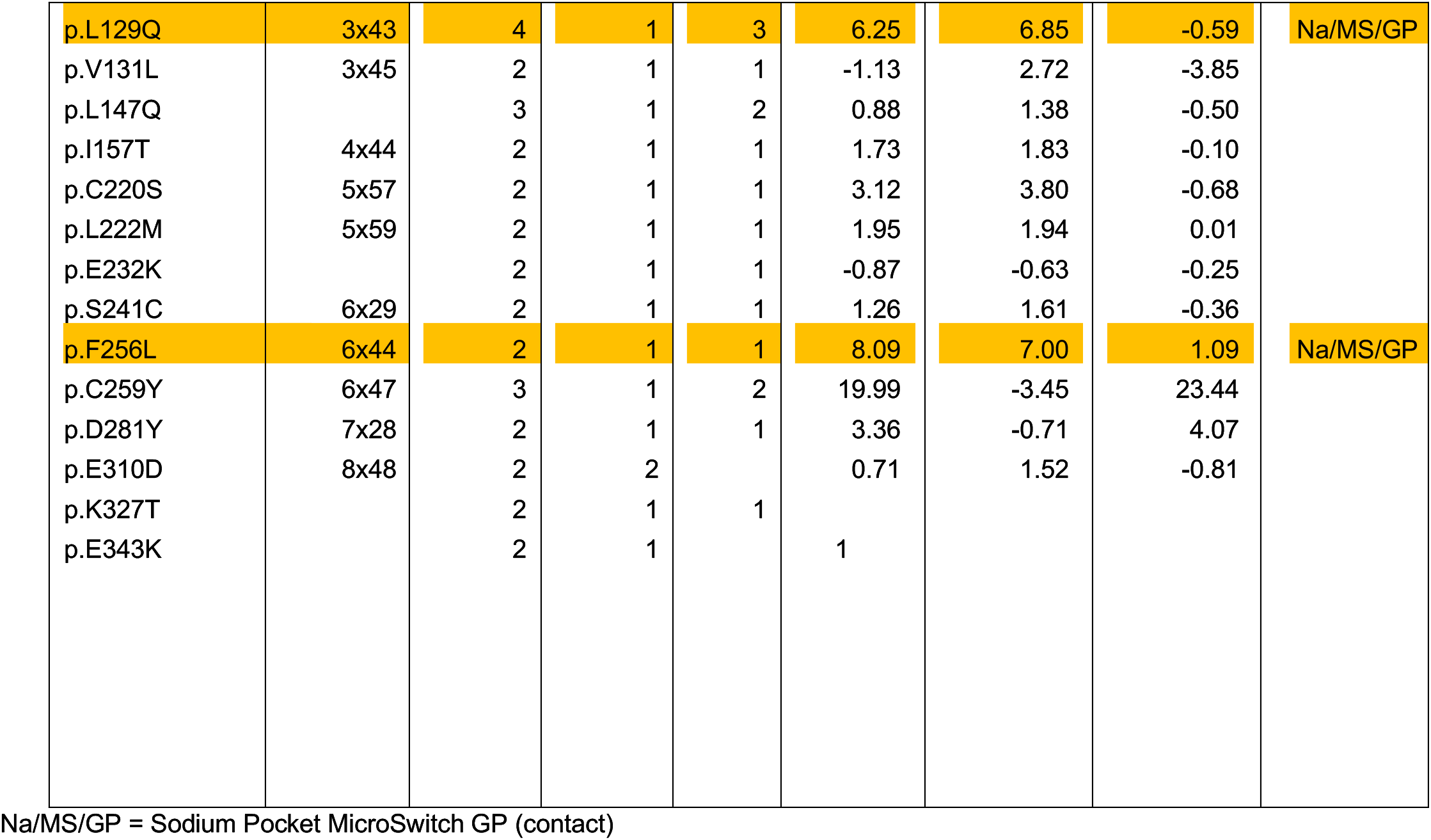
“Cancer” mutations found as somatic variants in the TCGA and COSMIC Databases

**Suppl. Table S8.**
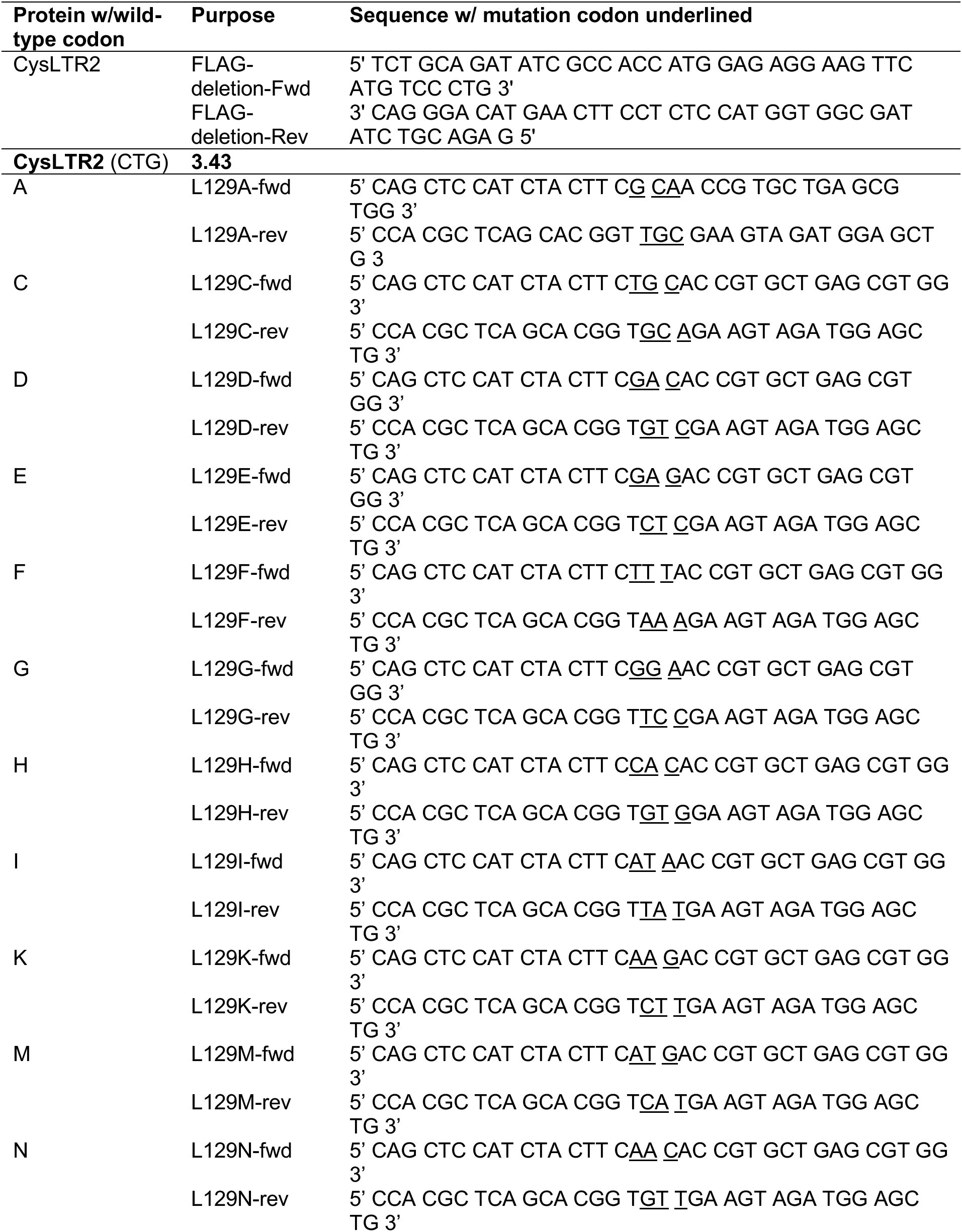
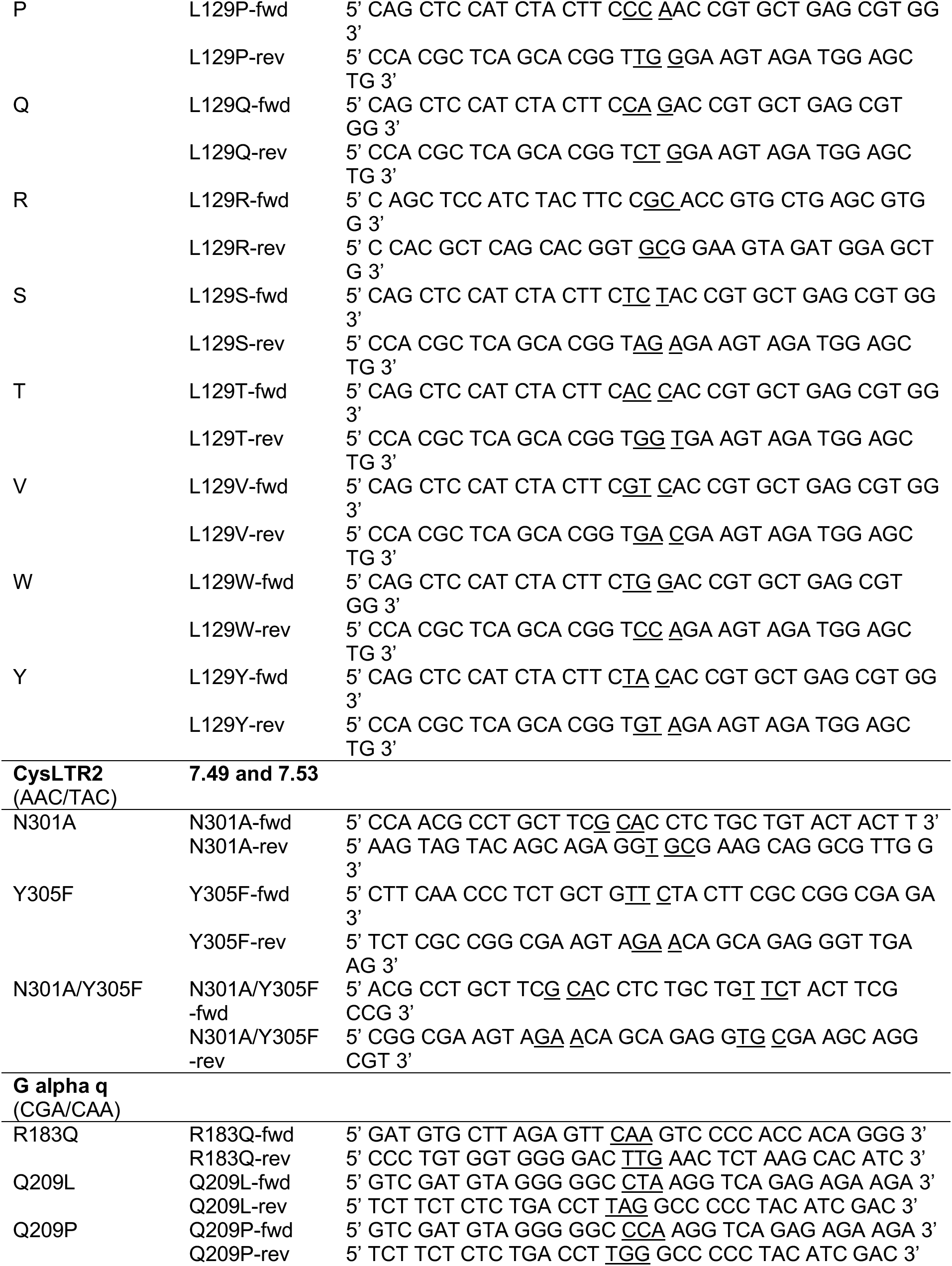
Primers used to generate CysLTR2 3.43, 7.49, and 7.53 mutants and Gαq protein mutants

**Suppl. Table S9.**
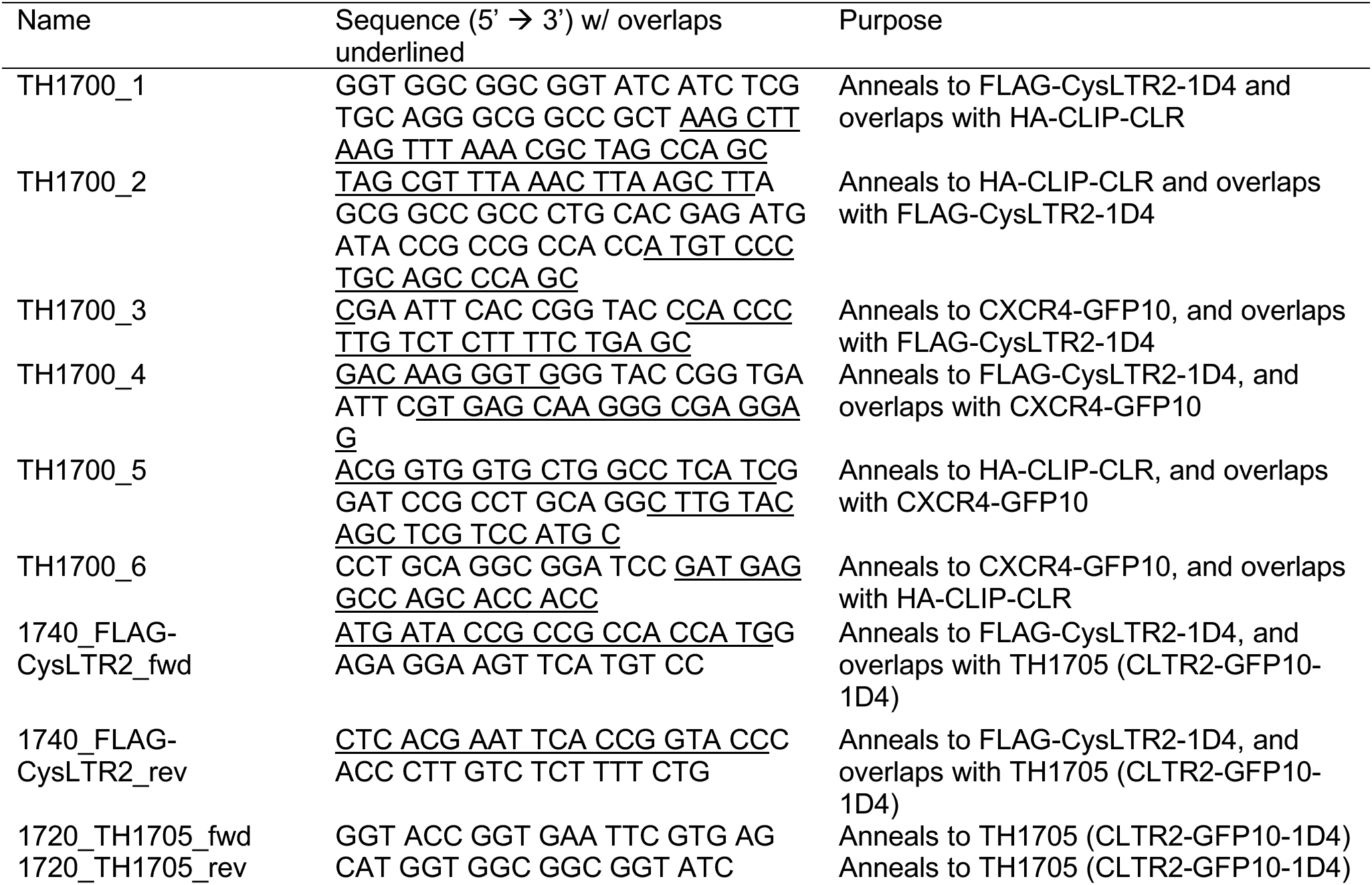
Primers used in building BRET2 acceptors

## References

1. Vass M, Kooistra AJ, Yang D, Stevens RC, Wang MW, de Graaf C. Chemical Diversity in the G Protein-Coupled Receptor Superfamily. Trends Pharmacol Sci 39, 494–512 (2018).

2. Oprea TI, et al. Unexplored therapeutic opportunities in the human genome. Nat Rev Drug Discov 17, 317–332 (2018).

3. Hauser AS, et al. Pharmacogenomics of GPCR Drug Targets. Cell 172, 41–54 e19 (2018).

4. Kan Z, et al. Diverse somatic mutation patterns and pathway alterations in human cancers. Nature 466, 869–873 (2010).

5. Prickett TD, et al. Exon capture analysis of G protein-coupled receptors identifies activating mutations in GRM3 in melanoma. Nat Genet 43, 1119–1126 (2011).

6. Moore AR, et al. Recurrent activating mutations of G-protein-coupled receptor CYSLTR2 in uveal melanoma. Nat Genet 48, 675–680 (2016).

7. Li MM, et al. Standards and Guidelines for the Interpretation and Reporting of Sequence Variants in Cancer: A Joint Consensus Recommendation of the Association for Molecular Pathology, American Society of Clinical Oncology, and College of American Pathologists. J Mol Diagn 19, 4–23 (2017).

8. Richards S, et al. Standards and guidelines for the interpretation of sequence variants: a joint consensus recommendation of the American College of Medical Genetics and Genomics and the Association for Molecular Pathology. Genet Med 17, 405–424 (2015).

9. Moller I, et al. Activating cysteinyl leukotriene receptor 2 (CYSLTR2) mutations in blue nevi. Mod Pathol 30, 350–356 (2017).

10. van de Nes JAP, et al. Activating CYSLTR2 and PLCB4 Mutations in Primary Leptomeningeal Melanocytic Tumors. J Invest Dermatol 137, 2033–2035 (2017).

11. Kusters-Vandevelde HVN, et al. Whole-exome sequencing of a meningeal melanocytic tumour reveals activating CYSLTR2 and EIF1AX hotspot mutations and similarities to uveal melanoma. Brain Tumor Pathol 35, 127–130 (2018).

12. Smit MJ, et al. Pharmacogenomic and structural analysis of constitutive g protein-coupled receptor activity. Annu Rev Pharmacol Toxicol 47, 53–87 (2007).

13. Stoy H, Gurevich VV. How genetic errors in GPCRs affect their function: Possible therapeutic strategies. Genes Dis 2, 108–132 (2015).

14. Vazquez-Prado J, Bracho-Valdes I, Cervantes-Villagrana RD, Reyes-Cruz G. Gbetagamma Pathways in Cell Polarity and Migration Linked to Oncogenic GPCR Signaling: Potential Relevance in Tumor Microenvironment. Mol Pharmacol 90, 573–586 (2016).

15. Fukami M, Suzuki E, Igarashi M, Miyado M, Ogata T. Gain-of-function mutations in G-protein-coupled receptor genes associated with human endocrine disorders. Clin Endocrinol (Oxf) 88, 351–359 (2018).

16. Heise CE, et al. Characterization of the human cysteinyl leukotriene 2 receptor. J Biol Chem 275, 30531–30536 (2000).

17. Black JW, Leff P. Operational models of pharmacological agonism. Proc R Soc Lond B Biol Sci 220, 141–162 (1983).

18. Slack RJ, Hall DA. Development of operational models of receptor activation including constitutive receptor activity and their use to determine the efficacy of the chemokine CCL17 at the CC chemokine receptor CCR4. Br J Pharmacol 166, 1774–1792 (2012).

19. Zhou B, Hall DA, Giraldo J. Can Adding Constitutive Receptor Activity Redefine Biased Signaling Quantification? Trends Pharmacol Sci 40, 156–160 (2019).

20. Kenakin T, Watson C, Muniz-Medina V, Christopoulos A, Novick S. A simple method for quantifying functional selectivity and agonist bias. ACS Chem Neurosci 3, 193–203 (2012).

21. Luttrell LM, Lefkowitz RJ. The role of beta-arrestins in the termination and transduction of G-protein-coupled receptor signals. J Cell Sci 115, 455–465 (2002).

22. Yan D, Stocco R, Sawyer N, Nesheim ME, Abramovitz M, Funk CD. Differential signaling of cysteinyl leukotrienes and a novel cysteinyl leukotriene receptor 2 (CysLT(2)) agonist, N-methyl-leukotriene C(4), in calcium reporter and beta arrestin assays. Mol Pharmacol 79, 270–278 (2011).

23. Leduc M, et al. Functional selectivity of natural and synthetic prostaglandin EP4 receptor ligands. J Pharmacol Exp Ther 331, 297–307 (2009).

24. Berchiche YA, Sakmar TP. CXC Chemokine Receptor 3 Alternative Splice Variants Selectively Activate Different Signaling Pathways. Mol Pharmacol 90, 483–495 (2016).

25. Hamdan FF, Audet M, Garneau P, Pelletier J, Bouvier M. High-throughput screening of G protein-coupled receptor antagonists using a bioluminescence resonance energy transfer 1-based beta-arrestin2 recruitment assay. J Biomol Screen 10, 463–475 (2005).

26. Hulme EC, Trevethick MA. Ligand binding assays at equilibrium: validation and interpretation. Br J Pharmacol 161, 1219–1237 (2010).

27. Pfleger KD, Eidne KA. Illuminating insights into protein-protein interactions using bioluminescence resonance energy transfer (BRET). Nat Methods 3, 165–174 (2006).

28. Mercier JF, Salahpour A, Angers S, Breit A, Bouvier M. Quantitative assessment of beta 1- and beta 2-adrenergic receptor homo- and heterodimerization by bioluminescence resonance energy transfer. J Biol Chem 277, 44925–44931 (2002).

29. Cagnol S, Chambard JC. ERK and cell death: mechanisms of ERK-induced cell death--apoptosis, autophagy and senescence. FEBS J 277, 2–21 (2010).

30. Nishihara E, et al. Sporadic congenital hyperthyroidism due to a germline mutation in the thyrotropin receptor gene (Leu 512 Gln) in a Japanese patient. Endocr J 53, 735–740 (2006).

31. Lu ZL, Hulme EC. The functional topography of transmembrane domain 3 of the M1 muscarinic acetylcholine receptor, revealed by scanning mutagenesis. J Biol Chem 274, 7309–7315 (1999).

32. Tao YX, Abell AN, Liu X, Nakamura K, Segaloff DL. Constitutive activation of G protein-coupled receptors as a result of selective substitution of a conserved leucine residue in transmembrane helix III. Mol Endocrinol 14, 1272–1282 (2000).

33. Fredriksson R, Lagerstrom MC, Lundin LG, Schioth HB. The G-protein-coupled receptors in the human genome form five main families. Phylogenetic analysis, paralogon groups, and fingerprints. Mol Pharmacol 63, 1256–1272 (2003).

34. Pandy-Szekeres G, et al. GPCRdb in 2018: adding GPCR structure models and ligands. Nucleic Acids Research 46, D440–D446 (2018).

35. Zheng Y, et al. Structure of CC Chemokine Receptor 5 with a Potent Chemokine Antagonist RevealsMechanisms of Chemokine Recognition and Molecular Mimicry by HIV. Immunity 46, 1005-+ (2017).

36. Taniguchi R, et al. Structural insights into ligand recognition by the lysophosphatidic acid receptor LPA(6). Nature 548, 356-+ (2017).

37. Kellogg EH, Leaver-Fay A, Baker D. Role of conformational sampling in computing mutation-induced changes in protein structure and stability. Proteins 79, 830–838 (2011).

38. Zhang C, et al. High-resolution crystal structure of human protease-activated receptor 1. Nature 492, 387–392 (2012).

39. Katritch V, Fenalti G, Abola EE, Roth BL, Cherezov V, Stevens RC. Allosteric sodium in class A GPCR signaling. Trends Biochem Sci 39, 233–244 (2014).

40. Liu W, et al. Structural basis for allosteric regulation of GPCRs by sodium ions. Science 337, 232–236 (2012).

41. Takasaki J, et al. A novel Galphaq/11-selective inhibitor. J Biol Chem 279, 47438–47445 (2004).

42. Yoo JH, et al. ARF6 Is an Actionable Node that Orchestrates Oncogenic GNAQ Signaling in Uveal Melanoma. Cancer Cell 29, 889–904 (2016).

43. Shenoy SK, Lefkowitz RJ. beta-Arrestin-mediated receptor trafficking and signal transduction. Trends Pharmacol Sci 32, 521–533 (2011).

44. Dores MR, Trejo J. Endo-lysosomal sorting of G-protein-coupled receptors by ubiquitin: Diverse pathways for G-protein-coupled receptor destruction and beyond. Traffic 20, 101–109 (2019).

45. de Rubio RG, Ransom RF, Malik S, Yule DI, Anantharam A, Smrcka AV. Phosphatidylinositol 4-phosphate is a major source of GPCR-stimulated phosphoinositide production. Sci Signal 11, (2018).

46. van den Bout I, Divecha N. PIP5K-driven PtdIns(4,5)P2 synthesis: regulation and cellular functions. J Cell Sci 122, 3837–3850 (2009).

47. Bruntz RC, Lindsley CW, Brown HA. Phospholipase D signaling pathways and phosphatidic acid as therapeutic targets in cancer. Pharmacol Rev 66, 1033–1079 (2014).

48. Vaque JP, et al. A genome-wide RNAi screen reveals a Trio-regulated Rho GTPase circuitry transducing mitogenic signals initiated by G protein-coupled receptors. Mol Cell 49, 94–108 (2013).

49. Feng X, et al. Hippo-independent activation of YAP by the GNAQ uveal melanoma oncogene through a trio-regulated rho GTPase signaling circuitry. Cancer Cell 25, 831–845 (2014).

50. Yu FX, et al. Mutant Gq/11 promote uveal melanoma tumorigenesis by activating YAP. Cancer Cell 25, 822–830 (2014).

51. D’Souza-Schorey C, Chavrier P. ARF proteins: roles in membrane traffic and beyond. Nat Rev Mol Cell Biol 7, 347–358 (2006).

52. Giguere P, et al. ARF6 activation by Galpha q signaling: Galpha q forms molecular complexes with ARNO and ARF6. Cell Signal 18, 1988–1994 (2006).

53. Laroche G, Giguere PM, Dupre E, Dupuis G, Parent JL. The N-terminal coiled-coil domain of the cytohesin/ARNO family of guanine nucleotide exchange factors interacts with Galphaq. Mol Cell Biochem 306, 141–152 (2007).

54. Grossmann AH, et al. The small GTPase ARF6 stimulates beta-catenin transcriptional activity during WNT5A-mediated melanoma invasion and metastasis. Sci Signal 6, ra14 (2013).

55. Moore AR, et al. GNA11 Q209L Mouse Model Reveals RasGRP3 as an Essential Signaling Node in Uveal Melanoma. Cell Rep 22, 2455–2468 (2018).

56. Chen X, et al. RasGRP3 Mediates MAPK Pathway Activation in GNAQ Mutant Uveal Melanoma. Cancer Cell 31, 685–696 e686 (2017).

57. Terrillon S, Bouvier M. Receptor activity-independent recruitment of betaarrestin2 reveals specific signalling modes. EMBO J 23, 3950–3961 (2004).

58. Luttrell LM, et al. Manifold roles of beta-arrestins in GPCR signaling elucidated with siRNA and CRISPR/Cas9. Sci Signal 11, (2018).

59. Trinquet E, et al. D-myo-inositol 1-phosphate as a surrogate of D-myo-inositol 1,4,5-tris phosphate to monitor G protein-coupled receptor activation. Anal Biochem 358, 126–135 (2006).

60. Lorenzen E, et al. Multiplexed Analysis of the Secretin-like GPCR-RAMP Interactome. bioRxiv, 597690 (2019).

61. Hamdan FF, Percherancier Y, Breton B, Bouvier M. Monitoring protein-protein interactions in living cells by bioluminescence resonance energy transfer (BRET). Curr Protoc Neurosci Chapter 5, Unit 5 23 (2006).

62. Koehler Leman J, Mueller BK, Gray JJ. Expanding the toolkit for membrane protein modeling in Rosetta. Bioinformatics 33, 754–756 (2017).

63. Leaver-Fay A, et al. ROSETTA3: an object-oriented software suite for the simulation and design of macromolecules. Methods Enzymol 487, 545–574 (2011).

64. Alford RF, et al. The Rosetta All-Atom Energy Function for Macromolecular Modeling and Design. J Chem Theory Comput 13, 3031–3048 (2017).

65. Humphrey W, Dalke A, Schulten K. VMD: visual molecular dynamics. J Mol Graph 14, 33–38, 27-38 (1996).

66. Wunder F, et al. Pharmacological characterization of the first potent and selective antagonist at the cysteinyl leukotriene 2 (CysLT(2)) receptor. Br J Pharmacol 160, 399–409 (2010).

67. Ehlert FJ. Analysis of Biased Agonism. Prog Mol Biol Transl Sci 160, 63–104 (2018).

68. Samama P, Cotecchia S, Costa T, Lefkowitz RJ. A mutation-induced activated state of the beta 2-adrenergic receptor. Extending the ternary complex model. J Biol Chem 268, 4625–4636 (1993).

69. Kenakin T. Theoretical Aspects of GPCR-Ligand Complex Pharmacology. Chem Rev 117, 4–20 (2017).

